# LTP induction boosts glutamate spillover by driving withdrawal of astroglia

**DOI:** 10.1101/349233

**Authors:** Christian Henneberger, Lucie Bard, Aude Panatier, James P. Reynolds, Olga Kopach, Nikolay I. Medvedev, Daniel Minge, Michel K. Herde, Stefanie Anders, Igor Kraev, Janosch P. Heller, Sylvain Rama, Kaiyu Zheng, Thomas P. Jensen, Inmaculada Sanchez-Romero, Colin Jackson, Harald Janovjak, Ole Petter Ottersen, Erlend Arnulf Nagelhus, Stephane H.R. Oliet, Michael G. Stewart, U. Valentin Nägerl, Dmitri A. Rusakov

**Affiliations:** UCL Institute of Neurology, University College London, UK; Institute of Cellular Neurosciences, University of Bonn Medical School, Germany; Inserm U1215, Neurocentre Magendie, Bordeaux, France; Université de Bordeaux, Bordeaux, France; Life Sciences, The Open University, Milton Keynes, UK; Institute of Science and Technology Austria (IST Austria), 3400 Klosterneuburg, Austria; Institute of Basic Medical Sciences, University of Oslo, 0317 Oslo, Norway; Interdisciplinary Institute for Neuroscience, CNRS UMR 5297, Bordeaux, France; German Center for Neurodegenerative Diseases (DZNE), Bonn, Germany; Research School of Chemistry, Australian National University, Canberra, Australia

**Keywords:** synaptic transmission, long-term potentiation, glutamate spillover, STED, dSTORM, astroglial glutamate transporters, optical glutamate sensor, inter-synaptic crosstalk, signal integration

## Abstract

Extrasynaptic actions of glutamate are limited by high-affinity transporters on perisynaptic astroglial processes (PAPs), which helps to maintain point-to-point transmission in excitatory circuits. Memory formation in the brain is associated with synaptic remodelling, but how this remodelling affects PAPs and therefore extrasynaptic glutamate actions is poorly understood. Here we used advanced imaging methods, *in situ* and *in vivo*, to find that a classical synaptic memory mechanism, long-term potentiation (LTP), triggers withdrawal of PAPs from potentiated synapses. Optical glutamate sensors combined with patch-clamp and 3D molecular localisation reveal that LTP induction thus prompts a spatial retreat of glial glutamate transporters, boosting glutamate spillover and NMDA receptor-mediated inter-synaptic cross-talk. The LTP-triggered PAP withdrawal involves NKCC1 transporters and the actin-controlling protein cofilin but does not depend on major Ca^2+^-dependent cascades in astrocytes. We have therefore uncovered a mechanism by which synaptic potentiation could alter signal handling by multiple nearby connections.

## INTRODUCTION

The plasma membrane of brain astroglia is densely packed with high-affinity transporters that rapidly take up glutamate released by excitatory synapses (Danbolt, 2001). Perisynaptic astroglial processes (PAPs) often occur in close proximity to the synaptic cleft (Grosche et al., 1999; Heller and Rusakov, 2015; Ventura and Harris, 1999), to hinder released glutamate from activating its receptors on neighbouring synapses. However, astroglia-dependent extrasynaptic glutamate escape, or glutamate ‘spillover’, has long been acknowledged to have a significant physiological impact (Dittman and Regehr, 1997; Kullmann and Asztely, 1998; Lozovaya et al., 1999; Rusakov et al., 1999). In the hippocampus, glutamate spillover has been mechanistically implicated in a co-operative action (including ‘priming’) of dendritic NMDA receptors (NMDARs) (Chalifoux and Carter, 2011; Hires et al., 2008), functional inter-synaptic cross-talk (Arnth-Jensen et al., 2002; Lozovaya et al., 1999; Scimemi et al., 2004), heterosynaptic potentiation and depression (Vogt and Nicoll, 1999), and remote activation of metabotropic glutamate receptors (Min et al., 1998; Scanziani et al., 1997), among other phenomena. Glutamate escape underlies signalling between mitral cells in the olfactory bulb (Isaacson, 1999), and between climbing fibres and interneurons (Coddington et al., 2013; Szapiro and Barbour, 2007), as well as between parallel fibres and stellate cells (Carter and Regehr, 2000) in the cerebellum. At the behavioural level, a causal relationship has been demonstrated between changes in the astroglia-dependent extrasynaptic glutamate escape and cognitive decline (Pereira et al., 2014), fear conditioning behaviour (Tanaka et al., 2013; Tsvetkov et al., 2004), heroin and cocaine relapse (Shen et al., 2014; Smith et al., 2017), among other effects. However, whether and how the degree of glutamate spillover controlled by PAPs is regulated by neural activity has remained poorly understood.

Astrocytes have also emerged as a source of molecular signals that regulate synaptic transmission (Jourdain et al., 2007; Navarrete and Araque, 2010; Pascual et al., 2005; Santello et al., 2011) and contribute to the long-term modifications of synaptic circuitry associated with memory formation (Adamsky et al., 2018; Henneberger et al., 2010; Min and Nevian, 2012; Shigetomi et al., 2013). Molecular exchange between astrocytes and synapses is thought to depend on the occurrence and function of PAPs (Panatier et al., 2006; Panatier et al., 2011). It has been therefore a long-standing question of whether PAPs undergo activity-dependent remodelling, which affects the functioning of local circuitry. Electron microscopy (EM) studies in fixed tissue have reported increased astroglial coverage of synapses in samples that underwent induction of synaptic long-term potentiation (LTP) (Bernardinelli et al., 2014; Lushnikova et al., 2009; Wenzel et al., 1991). An increase in the PAP occurrence has also been found in animals reared in complex environment (Jones and Greenough, 1996). In contrast, synaptic coverage by PAPs was shown to decrease following memory consolidation tasks (Ostroff et al., 2014) or during lactation (Oliet et al., 2001). Nevertheless, EM cannot follow in real time physiological events and could be susceptible to distortions of PAP morphology during tissue fixation (Korogod et al., 2015).

These factors necessitate evidence in live cells. Obtaining such evidence has been a challenge because of the small PAP dimensions. Several previous studies have employed confocal or two-photon excitation (2PE) microscopy to monitor fine changes in PAPs using fluorescent labelling (Bernardinelli et al., 2014; Haber et al., 2006; Hirrlinger et al., 2004; Perez-Alvarez et al., 2014). However, the apparent dynamics of fluorescent shapes in a conventional microscope is subject to interpretation. Firstly, the size of PAPs and inter-PAP space gaps are beyond the light diffraction limit, potentially giving rise to spurious morphologies (Rusakov, 2015). Secondly, cell-permeable fluorescent tracers appear to underreport astroglial structure compared to soluble cytosolic dyes (Reeves et al., 2011). Finally, sub-micron sample drift or local re-arrangement (or photobleaching) of the fluorescent label could be mistaken for changes in PAP shape or motility.

To avoid such uncertainties, we set out to monitor PAPs with advanced microscopy methods that are not limited by diffraction of light, under several LTP induction protocols, in hippocampal slices and in the barrel cortex *in vivo*. We used optical glutamate sensors and patch-clamp electrophysiology to determine how the LTP-associated change in PAPs affects extrasynaptic glutamate escape. We probed multiple astrocytic signalling cascades that could underpin such changes, and identified the key players. The results unveil the mechanism by which plasticity-inducing patterns of neural activity trigger local PAP remodelling and thus alter the postsynaptic integration of excitatory inputs.

## RESULTS

### LTP induction reduces PAP volume

First, we monitored astrocytes that were whole-cell loaded with the soluble morphological tracer Alexa Fluor 594 (AF 594), in the CA1 *stratum radiatum* of acute hippocampal slices (Methods). In these settings, the average fluorescence intensity *F_ROI_* collected over a selected ROI (within a ∼1 µm think optical section) is proportional to the tissue volume fraction (VF) occupied by dye-filled PAPs within this ROI (Figure 1A, *left*). Because astrocytes occupy non-overlapping tissue domains (Bushong et al., 2002), there will be no PAPs from other astroglia in the sampled ROI. In each such ROI, the absolute VF of PAPs can be obtained by relating *F_ROI_* to the fluorescence intensity over the somatic region *F_S_,* which represents 100% VF (Figure 1A, *right*; Supplementary Figure 1A-B), as shown previously (Medvedev et al., 2014; Savtchenko et al., 2018). These measurements returned an average PAP VF value of 6-7% (cell bodies excluded; Figure S1B-C). This value is similar to that obtained previously in area CA1 (Savtchenko et al., 2018) and in the *dentate gyrus* (Medvedev et al., 2014), and is consistent with stereological EM data in CA1 neuropil (Lehre and Rusakov, 2002).

**Figure 1.**
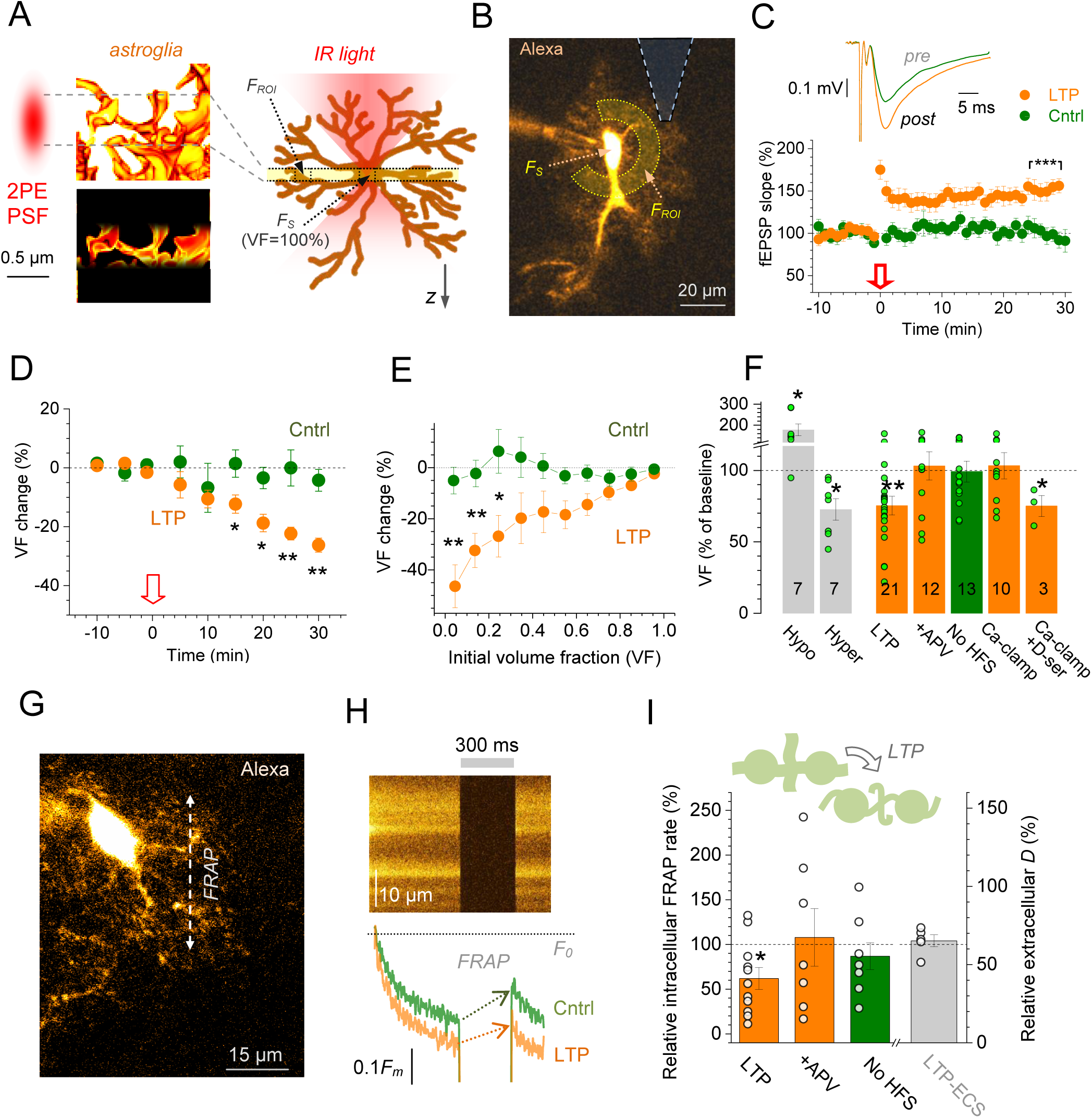
Reduced PAP presence after LTP induction at CA3-CA1 synapses. (A) *Left*: An illustration of the 2PE point-spread function (PSF, red) that excites dye-filled PAPs (yellow, 3D EM modified from (Medvedev et al., 2014)) only within a ∼1 µm focal section (dotted lines; *bottom*). *Right*: Local PAP fluorescence (*F_ROI_*) scales with their local VF, which reaches ∼100% inside the 5-7 µm wide soma (*F_S_*). (B) CA1 astrocyte held in whole-cell (single focal section; λ_x_^2p^ = 800 nm; AF 594; gap junctions blocked with 50 µM carbenoxolone); dashed cone, extracellular recording pipette. VF readout: average fluorescence over ROI (*F_ROI_*; dashed circular segment) relative to that within soma (*F_S_*). (C) *Traces*, Schaffer collateral-evoked fEPSPs near a patched astrocyte, before (*pre*) and ∼25 min after LTP induction (*post*); *Graph*, time course of normalised fEPSP (mean ± SEM, here and thereafter; arrow, LTP induction; ***p < 0.001 (potentiation 25-30 min post-induction: 151.0 ± 6.7% compared to baseline, *n =* 18). (D) Relative change in PAP VF (%) in control conditions (green; *n =* 9) and after LTP induction (orange; *n =* 13 cells; *p *<* 0.05; **p < 0.01, compared to control). (E) Relative change in PAP VF (%, sampled over 0.3 x 0.3 µm ROIs across cell arbour, averaged for individual cells) plotted against the initial PAP VF, in control (*n =* 8) and ∼25 min after LTP induction (orange; *n =* 13; *p *<* 0.05, **p *<* 0.01, compared to control). (F) *Grey*, Relative change in PAP VF (%, sample size *n* shown) in hypo-osmotic (220 mOsm/l) and hyper-osmotic (420 mOsm/l) solutions, as shown. *Green and orange*: Relative PAP VF change 25-30 min after LTP induction in control (LTP, −25 ± 7%), in 50 µM APV (+APV, 3.1 ± 9.9%), with no HFS (−0.8 ± 7.3%), under intra-astrocyte Ca^2+^ clamp (Ca-clamp, 6.8 ± 9.5%) (Henneberger et al., 2010), under Ca^2+^ clamp with 10 µM D-serine added (Ca-clamp+ D-ser, −24 ± 7%); **p < 0.01; *p < 0.05; dots, individual tests. (G) Evaluating diffusion coupling inside CA1 astroglia using FRAP of intracellular AF 594 (Methods; single focal section; ∼80 µm depth); arrow, example linescan position. (H) *Top*: Example linescan as in G (baseline conditions; grey segment, shutter closed). *Bottom*: The corresponding fluorescence intensity time course, before and ∼20 min after LTP induction, as indicated; *F_0_*, initial intensity; arrows, FRAP during shutter-on period (full recovery takes 39-40 s). (I) Summary of FRAP experiments (G-H); diagram, LTP induction reduces thinner parts of PAPs thus lowering diffusion coupling. Bar graph, FRAP rate relative to baseline (left ordinate): ∼25 min after LTP induction (LTP, 62 ± 12%, *n =* 11; *p *<* 0.05); in the presence of APV (108 ± 32%, *n =* 7); no-HFS control (87 ± 15%, *n =* 7). *Grey bar* (right ordinate): relative extracellular diffusivity ∼25 min after LTP induction (LTP-ECS; 104 ± 7%, *n =* 5; Figure S1F-H); dots, individual test.

Next, we induced LTP at CA3-CA1 synapses using the classical protocol of high-frequency stimulation (HFS) applied to Schaffer collaterals while monitoring PAP VF and fEPSPs in the astrocyte arbour area, as shown previously (Henneberger et al., 2010) (Figure 1B-C; Methods). LTP induction prompted a progressive VF decrease lasting for at least 30 min (Figure 1D). No such changes occurred in control conditions (Figure 1D), ruling out confounding effects, such as photobleaching. Interestingly, astroglial areas showing the smallest initial PAP VF underwent the strongest VF reduction (Figure 1E). We also documented an LTP-associated PAP VF reduction in EGFP-expressing astroglia, in similar tests: the effect was smaller (most likely due to restricted EGFP diffusion, compared to AF). Furthermore, no changes were detected when use the protocol to induce long-term depression (LTD, Supplementary Figure 1D-E).

The VF decrease was blocked when LTP induction was suppressed using either the NMDAR antagonist APV, or by clamping Ca^2+^ in the patched astrocyte (Figure 1F): Ca^2+^ clamp suppresses release of the NMDAR co-agonist D-serine from astroglia, which in turn curtails NMDAR activation by glutamate (Henneberger et al., 2010). However, both LTP and the VF reduction under Ca^2+^ clamp could be rescued by adding the NMDAR co-agonist D-serine to the bath (10 µM, Figure 1F), consistent with earlier findings (Adamsky et al., 2018; Henneberger et al., 2010). These tests suggested that the observed VF changes were specific to LTP induction rather than to the stimulation protocol *per se*.

### LTP induction reduces diffusion connectivity among astroglial processes

Fluorescence recovery after photobleaching (FRAP) of the soluble intracellular indicator Alexa can be used to monitor internal diffusion coupling among astrocyte processes, by applying linescan FRAP in the middle of the ‘spongiform’ astrocyte arbour (Figure 1G), as we showed earlier (Anders et al., 2014; Savtchenko et al., 2018). Here, we found that after LTP induction, the FRAP of AF was slower, with no such changes in control conditions (Figure 1H-I). These observations were consistent with the reduced diffusional coupling among PAP structures after LTP induction, possibly due to their partial shrinkage. At the same time, LTP induction had no effect on diffusion in the extracellular space (Figure 1I), right ordinate), assessed using a fluorescence point-source technique (Supplementary Figure 1F-H) as detailed earlier (Zheng et al., 2008). This was not surprising because *stratum radiatum* astroglia occupy only 6-7% of the tissue volume (Supplementary Figure 1C) (Savtchenko et al., 2018), of which 15-20% is taken by the extracellular space (Sykova and Nicholson, 2008). Thus, a 20-30% decrease in PAP VF would add on average only 5-10% to the extracellular space volume (1-2% overall tissue volume) within the affected area.

### STED imaging reveals decreased PAP presence near spines upon LTP induction

STED microscopy enables monitoring nanoscopic cellular compartments in live preparations, far beyond the optical diffraction limit (Tonnesen et al., 2018; Tonnesen et al., 2014). We therefore turned to two-colour STED imaging, combined with electrophysiology, in organotypic hippocampal slices (Panatier et al., 2014) (Methods, Supplementary Figure 2A; we used acute slices in all other methods). To chromatically separate CA1 pyramidal cells and astroglia, we used the Thy1-YFP transgenic mice (neurons, ‘red’ channel) and whole-cell loaded astrocytes AF 488 using a patch pipette (green channel).

We were thus able to monitor dendritic spines of CA1 pyramidal cells and local PAPs before and after LTP induction, with ∼70 nm resolution in the focal plane (Figure 2A). Again, to avoid subjective assessment of PAP changes, we employed volumetric readout, the ratio of green (astroglia) versus red (neuron) pixels (*G/R* values) in the 1.5 µm ROI centred at individual spine heads (Figure 2A). This ratio was decreased by 31 ± 10% following LTP induction (*n =* 22, p *<* 0.001; Figure 2B). This result corroborates the LTP-associated reduction in PAPs seen in acute slices (Figure 1A-F). Here, the unchanged red-pixel count (Figure 2B) rules out Thy1-YFP photobleaching; in the astroglia channel photobleaching was prevented by continuous AF dye dialysis.

**Figure 2.**
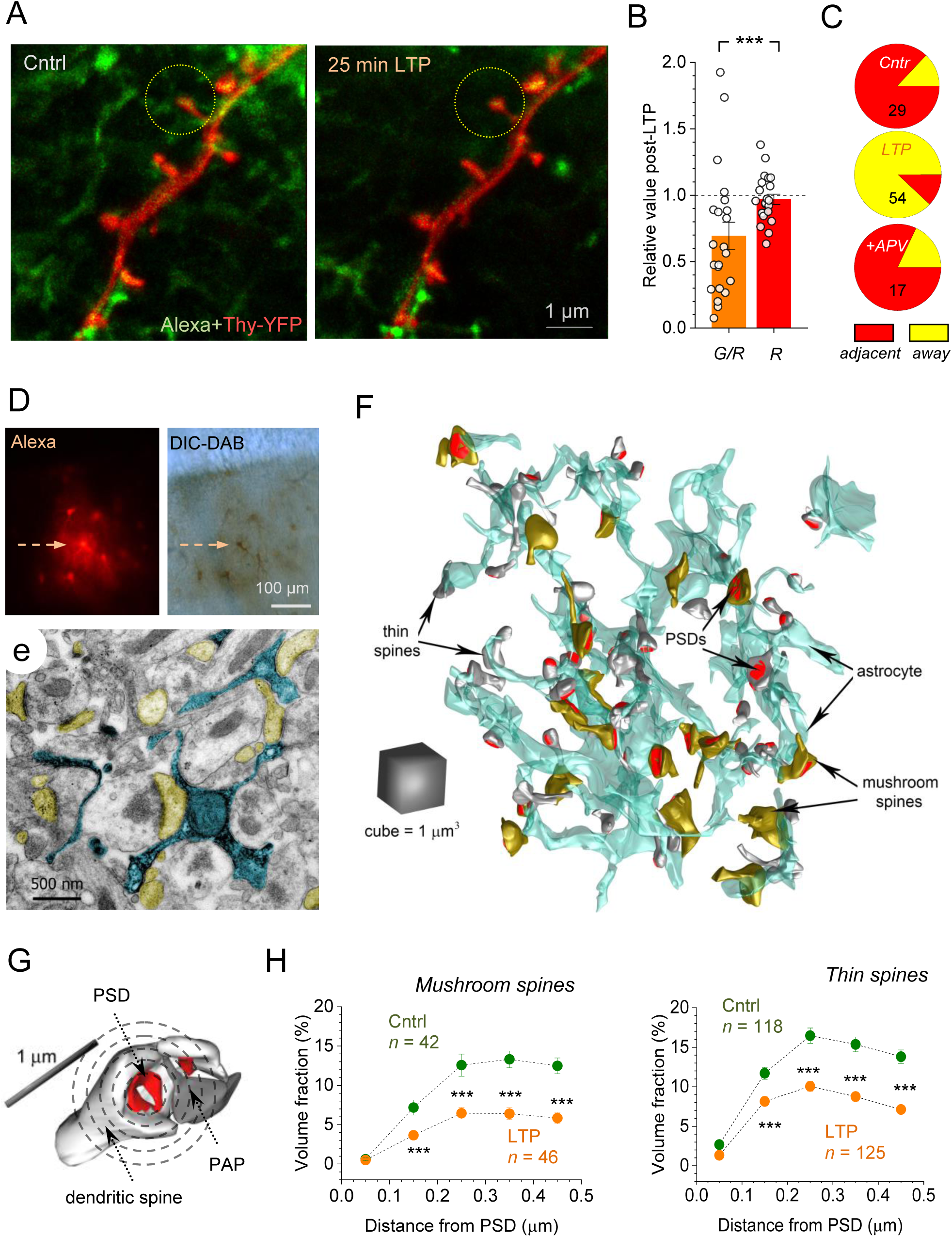
Live STED and correlational 3D EM report PAP withdrawal after LTP induction. (A) STED images of dendritic spines in a CA1 pyramidal neuron (red, Thy1-YFP) and nearby astroglia (green; 600 μM AF 488, whole-cell), before and ∼25 min after LTP induction, as indicated (Methods); dotted circles, ROIs centred at spine heads. (B) LTP induction reduces the green/red (astroglia/neuron) pixel-count ratio within spine-head centred ROIs (*G/R*; mean ± SEM; 31 ± 10%, *n =* 22, ***p *<* 0.001), with no effect on neuronal pixel count (*R*; −3.1 ± 3.8%); dots, individual test data. (C) Proportion of dendritic spines that appear adjacent to (red) and away from (yellow) PAPs, in baseline conditions (Cntr), 20-25 min after LTP induction (LTP), and the latter with 50 µM APV (+APV); numbers of spines shown (Figure S2A-D). (D) *Top*: patched astrocyte (arrow) loaded with biocytin (neighbouring astroglia stained through gap junctions), shown in fluorescence (*left*) and in DIC channel after DAB conversion (*right*). (E) Electron micrograph showing PAPs of the patched astrocyte (arrow in d) filled with precipitate (blue), and adjacent dendritic spines (yellow) featuring PSDs. (F) Fragment of a recorded astrocyte (as in D, cyan) reconstructed in 3D from ∼60 nm serial sections, including adjacent thin (white) and mushroom (yellow) dendritic spines containing PSDs (red; Figure S2E). (G) Integrative (volumetric) measure of synaptic astroglial coverage: VF of PAPs is calculated within 100 nm-wide concentric 3D shells (circles, not to scale) centred at the PSD (red). (H) PAP VF around PSDs (mean ± SEM), for adjacent thin and mushroom dendritic spines, in baseline conditions and ∼30 min after LTP induction, as indicated; sample sizes shown; ***p < 0.001.

STED images in the Thy1-YFP channel revealed subtle changes in some dendritic spines during LTP (Figure S2B). To explore this further while minimising photodamage during live STED, we compared randomised groups of spines between control and potentiated samples. Here, the fraction of spines in close physical contact with astroglial processes was reduced five-fold in the potentiated compared to control samples (Figure 2C). Interestingly, spines from which PAPs dissociated had larger heads in the potentiated sample, which also had a greater fraction of distinctly large spine heads (>500 nm wide, 12/54) overall, compared to control (3/29, Figure S2C-D). These observations suggest a complex picture of concurrent structural changes at the level of PAPs and spine heads following the HFS LTP induction protocol.

### Correlational 3D EM shows reduced occurrence of PAPs after LTP induction

To understand further the LTP-associated PAP changes on the nanoscale, we employed correlational quantitative 3D EM. We patched an astrocyte, in baseline conditions or 15-20 min after LTP induction at local synapses (Figure 1B) and loaded it with biocytin (Figure 2d). This was followed by rapid slice submersion into fixative and DAB conversion for EM identification (Figure 2D-E; Methods). The slices were cut into 60-70 nm thick serial sections and examined visually until the patched astrocyte (arrow in Figure 2D) could be found (Figure 2E). 200-300 serial sections were used to reconstruct an astrocyte fragment of interest and the adjacent synapses identified by their characteristic morphology (Figure 2F, Figure S2E), as detailed previously (Medvedev et al., 2014; Savtchenko et al., 2018).

To evaluate the extent of synaptic PAP coverage (in a shape-insensitive fashion), we calculated the PAP VF inside 100 nm–wide concentric spherical shells centred at individual PSDs (Figure 2G, Methods). Thus, we obtained the distribution of PAP coverage within ∼0.5 µm from the synapse, which is the average nearest-neighbour distance between CA3-CA1 synapses (Rusakov and Kullmann, 1998). Given that ‘thin’ and ‘mushroom’ spines in CA1 pyramidal cells may have distinct functional features (Matsuzaki et al., 2001), we treated these two morphological types separately. The analysis indicated clear astroglial withdrawal (or shrinkage away from spines) following LTP induction, for both spine types (Figure 2H). Our 3D EM data were consistent with our live imaging data, arguing that these results are unlikely to be biased by fixation artefacts (Korogod et al., 2015) (see Discussion).

### LTP-induced PAP withdrawal depends on activation of NKCC1

What are the cellular mechanisms underlying PAP withdrawal in these experiments? First, we examined whether one of the major astroglial Ca^2+^-signalling cascade that is known to engage metabotropic glutamate receptor (mGluR)-dependent activation of IP_3_ receptors (Porter and McCarthy, 1997; Volterra et al., 2014), and to alter PAP motility (Perez-Alvarez et al., 2014), is involved in PAP shrinkage. To test this, we spot-uncaged IP_3_ inside astrocyte branches using 2PE: this evoked robust local Ca^2+^ rises (Figure 3A-B) but had no effect on PAP VF (Figure 3C). Pressure-puff application of the wide-spectrum mGluR agonist DHGP gave a similar result (Figure 3C). PAP VF remained unchanged upon application of WIN55, an agonist of cannabinoid CB1 receptor (which contributes prominently to astroglial function (Navarrete and Araque, 2010)). Similarly, the GABA_A_ receptor agonist muscimol (which triggered slight shrinkage of sulforhodamine-101 stained astroglia (Florence et al., 2012) had no effect on PAP VF (Figure 3C).

**Figure 3.**
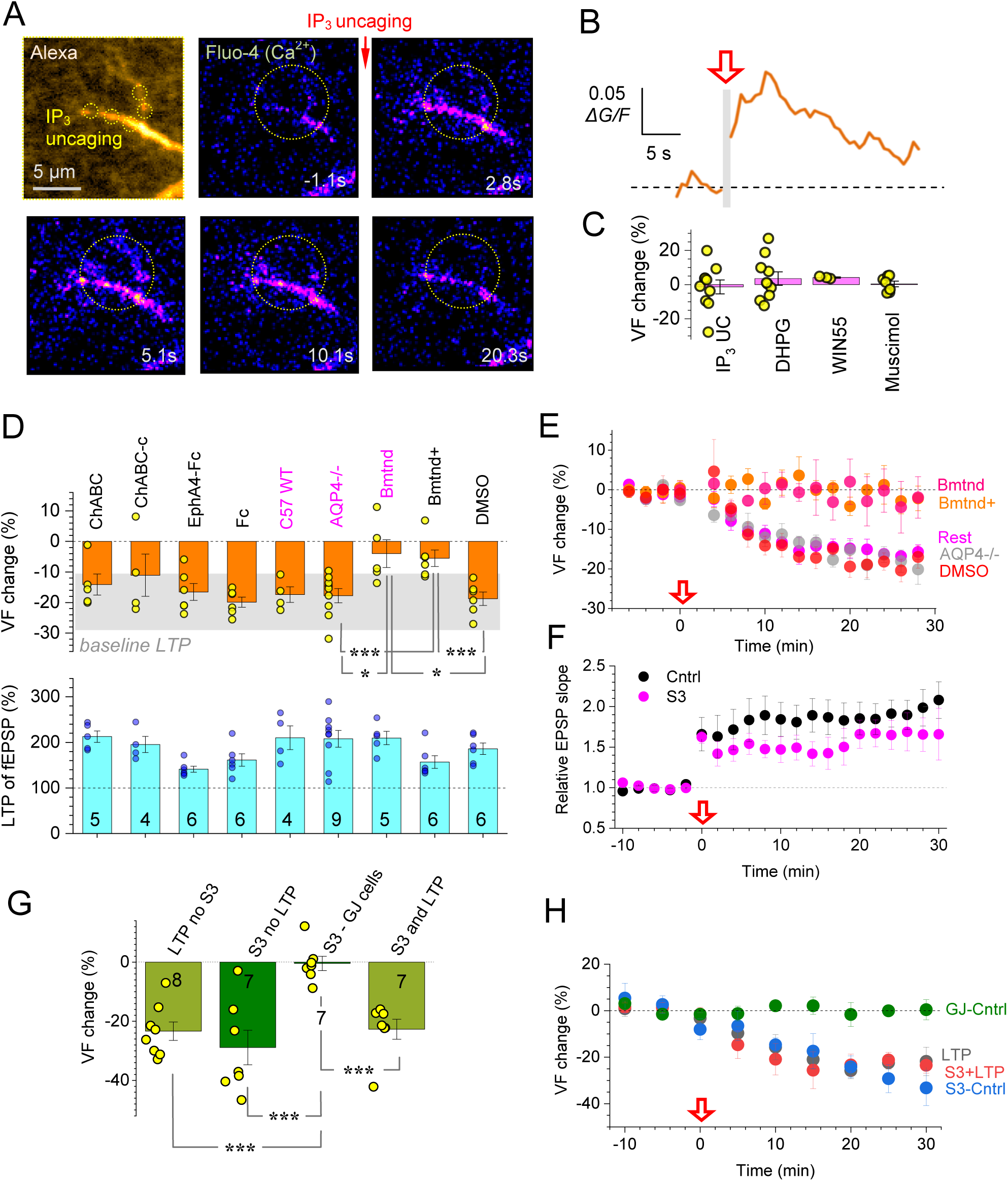
PAP withdrawal after LTP induction depends on NKCC1 and cofilin. (A) *Top left:* astrocyte fragment (5 µm z-stack), with uncaging spots (whole-cell 400 µM NPE-IP_3_; AF 594 channel, λ_x_^2P^ = 840 nm). *Other panels:* Time-lapse frame-scans of Ca^2+^ response (whole-cell Fluo-4; false colours) to IP_3_ spot-uncaging (at *t* = 0; five 5 ms pulses at 5Hz; λ_u_^2P^ = 720 nm); time shown; dotted circle, ROI for Ca^2+^ monitoring. (B) Time course of intracellular astroglial Ca^2+^ signal (*ΔF/G*) in the ROI shown in **a**; one-cell example; red arrow (grey segment), IP_3_ uncaging. (C) Relative change in PAP VF (%, mean ± SEM, here and thereafter) 25 min after: spot-uncaging of intracellular IP_3_ (−1.4 ± 4.1%, *n =* 10), pressure-puff application of group I mGluR agonist DHPG (300 µM, 3.5 ± 3.9%, *n =* 10), CB1 receptor agonist WIN55 (1 µM, 4.1 ± 0.4%, *n =* 3), and GABA receptor agonist muscimol (20 µM, 0.4 ± 1.7%, *n =* 6). (D) Relative change in PAP VF (%; *top*) ∼25 min after LTP induction, and the corresponding potentiation level (%; *bottom,* sample size shown): in the presence of 0.5-0.7 U/ml Chondroitinase ABC (ChABC, −14 ± 3%), control ChABC-c (−11 ± 7%), 10 µg/ml EphA4-Fc (−17 ± 3%), 10 µg/ml Fc control (−20 ± 2%), wild-type C57BI6 mice (−17 ± 3%), AQP4^-/-^ knockout mice (−18 ± 2%), 20 µM intracellular bumetanide (Bmtnd, −4 ± 4.5%), 50 µM intracellular bumetanide + 100 µM extracellular TGN-020 (Bmtnd+, −5.5 ± 2.7%), DMSO control 0.2% external + 0.05% internal (−19 ± 2%); blue text, data from mice; grey shadow, 95% confidence interval for PAP VF change after LTP induction in control conditions; *p < 0.02, **p < 0.01, ***p < 0.005 (*t*-test or Mann-Whitney independent sample tests). (E) Time course of PAP VF (mean ± SEM) during LTP induction (arrow) in key tests shown in (D), as indicated, including summary data for the rest of experiments (Rest). (F) Time course of fEPSP slope during LTP induction at CA3-CA1 synapses in control (*n =* 10) and with S3 peptide (200 µM) inside the adjacent astroglia (*n =* 6), as shown (Figure S3B-C). (G) Occlusion experiment: Relative change in PAP VF (sample size shown): ∼25 min post LTP induction in control (LTP no S3; −23 ± 3 %), no LTP induction with whole-cell loaded S3 (S3 no LTP; −29 ± 6 %); same test but recorded in gap-junction connected astrocytes devoid of S3 (S3 - GJ cells; −0.4 ± 2.4 %); and ∼25 min after LTP induction with S3 (S3 and LTP; −27 ± 3%). (H) Time course of PAP VF (relative to baseline) in the occlusion experiments shown in (G); notations as in (E-G).

We next tested morphogenic agents which have previously been associated with astroglia-dependent synaptic remodelling or stabilisation, such as the extracellular matrix (ECM) chondroitin sulfate (Dityatev and Schachner, 2003) and the ephrin/Eph cascades (Filosa et al., 2009; Murai et al., 2003; Nishida and Okabe, 2007). Neither the removal of chondroitin sulfate (with chondroitinase ABC) (Kochlamazashvili et al., 2010), nor the blockade of EphA4 activity with EphA4-Fc (Filosa et al., 2009; Murai et al., 2003) had a detectable effect on the LTP-induced reduction in PAP VF (Figure 3D).

We then turned to astroglial mechanisms of ion and water exchange, in which aquaporin-4 (AQP4) plays a prominent role (Nagelhus and Ottersen, 2013). However, in the AQP4 KO mice (Thrane et al., 2011), the LTP-associated changes in PAPs remained intact (Figure 3D-E). Another key player in cell volume regulation is the Na^+^-K^+^-2Cl^-^ cotransporter NKCC1, which is widely expressed in astroglia (Hoffmann et al., 2009; Kaila et al., 2014). To ensure cell specificity of our probing, we dialyzed individual astrocytes (whole-cell mode) with the NKCC1 blocker bumetanide (20 µM). Strikingly, bumetanide blocked changes of PAP VF while preserving LTP induction (Figure 3d-e): at the same time, basal PAP VF was not affected by bumetanide or by its vehicle DMSO (Figure S3A). This result was confirmed in rats with 50 µM intracellular bumetanide (Figure 3D-E; in these tests, 100 µM AQP4 blocker TGN-020 (Igarashi et al., 2011) was added to bath medium, to approach conditions of AQP4 KO, although see (Tradtrantip et al., 2017)). These results indicate that the LTP-induced PAP withdrawal depended on the activation of NKCC1.

### Activating cofilin cascade occludes LTP-induced changes in PAPs

What could be the downstream signal of NKCC1? In glioblastoma cells, NKCC1 was found to serve as a protein scaffold regulating the phosphorylation of the small, freely-diffusible protein cofilin-1 (Schiapparelli et al., 2017). In neurons, ion transporter KCC2 plays a similar role in cofilin control (Llano et al., 2015). Cofilin-1 is a well-established pH-dependent regulator of actin filament polymerization, which in turn controls formation and retrieval of thin cell protrusions, such as dendritic spines (Bravo-Cordero et al., 2013; Ethell and Pasquale, 2005). To probe this signalling cascade, we turned to intracellular dialysis of astroglia with a specific inhibitor peptide (S3) of cofilin-1 phosphorylation (Liu et al., 2016),(Aizawa et al., 2001). We confirmed that loading astrocytes with a solution containing P3 fully preserved LTP induction (Methods; Figure 3F, Figure S3B-C), as in the case of bumetanide.

Surprisingly, the peptide S3 dialysis alone, without LTP induction, triggered the PAP VF reduction similar to that in LTP experiments (Figure 3G). In the same experiment, astrocytes connected via gap-junctions to the patched cell showed no PAP changes, confirming a cell-specific effect (S3 - GJ cells; Figure 3G). However, when we induced LTP near the S3-containing astrocyte (Figure S3B-C), the magnitude and the time course of PAP shrinkage were indistinguishable from those during either LTP induction alone or under S3 dialysis alone (Figure 3H). If LTP induction and peptide S3 engaged separate cellular cascades, the effects of the two together on PAP VF should have been additive, in their amplitude and/or rate. However, the action of peptide S3 fully occluded the effect of LTP induction on PAP VF, thus indicating a shared molecular mechanism (see Discussion).

### Single-synapse LTP induction prompts local PAP retraction

Inducing LTP in the bulk of tissue will synchronously potentiate multiple synapses. However, memory trace formation in the brain may require synaptic changes to occur independently, at individual connections. It was therefore important to test whether the induction of LTP at individual synapses would affect local PAPs a similar way to the bulk-induced LTP.

Here, we modified the well-established protocol where LTP at individual CA3-CA1 connections is induced by glutamate spot-uncaging near a dendritic spine of the postsynaptic cell held in whole-cell (Harvey and Svoboda, 2007; Matsuzaki et al., 2004; Yasuda et al., 2003). First, we positioned the uncaging spot near a visualised dendritic spine (Figure 4A), and adjusted the uncaging laser power (pulse duration 1 ms) to generate a typical single-synapse EPSC waveform in the postsynaptic cell (Figure 4B). Second, we switched to current clamp while maintaining V_m_ at −60 to −65 mV, the range characteristic of CA1 pyramidal cells in freely-moving animals (Epsztein et al., 2010). In these conditions, we applied the spot-uncaging sequence aiming to mimics the HFS protocol but also to ensure that the postsynaptic cell is sufficiently depolarised to allow postsynaptic Ca^2+^ entry (Supplementary Figure 4A-B). Switching back to voltage clamp revealed clear potentiation of single-pulse EPSCs (Figure 4B), with LTP robustly induced at every recorded synapse (7 out of 7 cells, Figure 4B-C).

**Figure 4.**
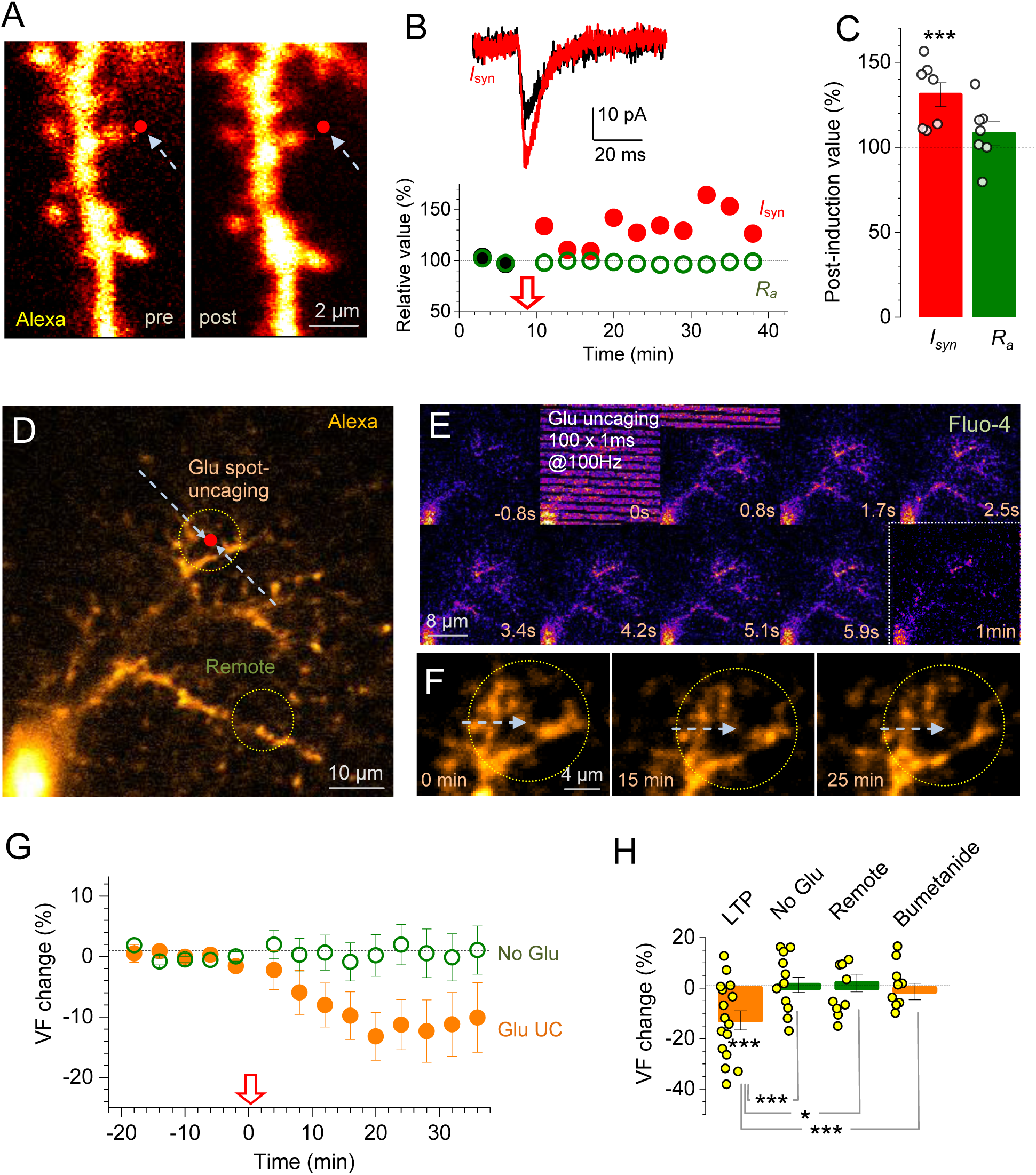
LTP induction at individual CA3-CA1 synapses reduces local PAP presence. (A) Dendritic fragment, CA1 pyramidal cell (AF channel), showing glutamate spot-uncaging position (red dot; 2.5 mM bath-applied MNI-glutamate) before (pre) and ∼20 min after LTP induction by spot-uncaging (post). (B) *Traces:* EPSCs (*I_syn_*, voltage-clamp) during baseline (black) and ∼30 min after LTP induction (red) in tests shown in A (Methods; dendritic Ca^2+^ dynamics shown in Figure S4A-B). *Graph*: one-spine example, time course of uncaging-evoked EPSC amplitude (*I_syn_*; black and red circles) and cell access resistance (*R_a_*, green) in the same experiment; red arrow, LTP induction onset. (C) Statistical summary of experiments depicted in (A-B) (mean ± SEM; *n =* 7, ***p < 0.005); notations as in (B); dots, individual tests. (D) Example, astrocyte fragment (whole-cell AF 594, single focal section); red dot, glutamate uncaging spot; circles, ROIs for PAP VF monitoring near the spot and at a remote site, as shown. (E) Time-lapse frames (area shown in D): astrocyte Ca^2+^ response (Fluo-4, λ_x_^2P^ = 840 nm) to the spot-uncaging LTP induction protocol (bleed-through at 0-0.8s frame; λ_u_^2P^ = 720 nm). (F) Astrocyte fragment near uncaging spot (as in D; arrow) immediately after (0 min), at 15 min and 25 min after LTP induction (∼9 µm *z*-stack average); PAP retraction could be seen at 15-25 min post-stimulus (Video 1; Figure S4C-E). (G) Time course of PAP VF change (%, mean ± SEM) in experiments shown in (D-E) (Glu, *n =* 11), and in control (no MNI-glutamate, No Glu, *n =* 11; arrow, uncaging onset). (H) Summary: PAP VF change (%, mean ± SEM) after LTP induction inset (LTP, −13 ± 4%, ***p < 0.005, *n =* 16), with no MNI-glutamate present (No Glu, 1.3 ± 3.0%, *n =* 9), in remote ROI (as in D; 2.0 ± 3.5%, *n =* 11), and with whole-cell loaded 20 µM bumetanide (−1.4 ± 3.3%, *n =* 9); *p < 0.05; ***p < 0.005; dots, individual tests; see Figure S4f-H for similar tests but with membrane-bound GFP.

Because CA3-CA1 synapses are only ∼0.5 µm apart (Rusakov and Kullmann, 1998), this protocol (Figure 4A-B) should potentiate at least one synapse near the uncaging spot, whether or not the unclamped postsynaptic cell is visualised. We therefore applied this protocol in *stratum radiatum* while monitoring PAP VF and Ca^2+^ in astrocytes loaded whole-cell with AF 594 and OGB-1, in the proximity of the uncaging spot (Figure 4D). The HFS uncaging sequence in most cases evoked a local Ca^2+^ rise in PAPs (Figure 4D-E, Figure S4D), which served as an indicator of robust glutamate release. In such cases, this protocol induced a progressive PAP VF reduction near the uncaging spot (Figure 4F-G; Figure S4C-E; Video 1). No detectable VF changes were found either in remote ROIs (>3 µm away from the spot, Figure 4D), when the protocol was applied without MNI-glutamate (Figure 4G-H). Importantly, the blockade of astroglial NKCC1 using whole-cell loaded bumetanide completely blocked the LTP-associated PAP changes (Figure 4H). A complementary experimental strategy, in which astrocytes were transduced with a membrane-bound GFP (AAV5.GfaABC1D.Pi.lck-GFP.SV40), rather than dialysed with a cytosolic AF dye, produced a qualitatively identical result, with the PAP withdrawal after LTP induction lasting for up to 100-120 min (Figure S4F-H).

### LTP induction increases distances between release machinery and GLT-1 transporters

Because PAP membranes are packed with the high-affinity glutamate transporter GLT-1 (Danbolt, 2001), we hypothesise that the LTP-associated PAP withdrawal alters perisynaptic arrangement of transporter molecules. To test this, we turned to dSTORM, a super-resolution microscopy technique that we adapted previously (Heller et al., 2017). Here, we mapped the 3D co-ordinates of the presynaptic protein bassoon, the postsynaptic density protein Homer1, and perisynaptic GLT-1. In order to potentiate the majority of synapses in the neuropil, we used the classical chemically-induced LTP (cLTP) protocol in acute hippocampal slices (Otmakhov et al., 2004), with full electrophysiological control (Figure S5A).

Three-color 3D dSTORM revealed a detailed spatial pattern for many hundreds of GLT-1 molecules occurring near individual synaptic contacts (Figure 5A; Figure S5B). We found that in the potentiated tissue GLT-1 occurred consistently further away from protein bassoon than in control (Figure 5B). Because bassoon is a key molecular partner of the synaptic vesicle release machinery (Gundelfinger et al., 2016), this result suggests that glutamate released from potentiated synapses has to travel, on average, comparatively larger distances before being taken up by GLT-1. Interestingly, we could not detect a similar trend for the distances between GLT-1 and Homer1 (Supplementary Figure 5C), suggesting that the withdrawal of PAPs from the presynaptic, axonal side could be more pronounced compared to postsynaptic withdrawal.

**Figure 5.**
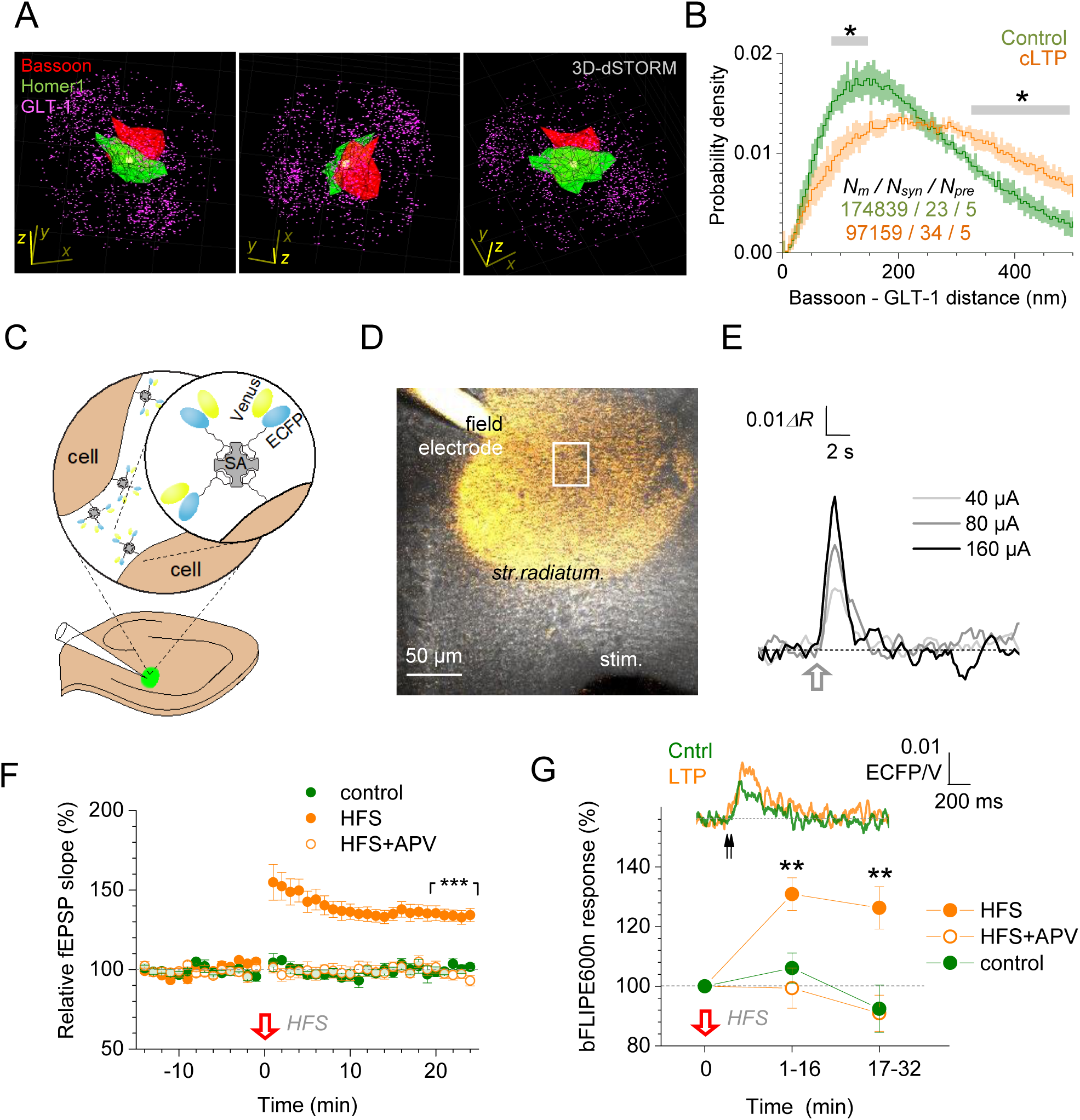
LTP induction triggers withdrawal of glial glutamate transporters boosting extracellular glutamate transient. (A) Spatial patterns of presynaptic bassoon (CF-568, red cluster), postsynaptic Homer 1 (Atto-488, green cluster), and glutamate transporter GLT-1 (Alexa-647, magenta dots) molecules visualised with 3D dSTORM; one-synapse example, three viewing angles shown (bassoon and Homer1 shown as clusters for clarity); *x-y-z* scale bars, 500 nm (Methods). (B) Average distribution (probability density, mean ± SEM) of the nearest-neighbour distances (<500 nm) between GLT-1 and presynaptic Bassoon molecules, in control tissue and ∼30 min after cLTP induction (Figure S5A-B; Methods); sample sizes: *N_m_*, inter-molecular distances; *N_syn_*, synapses; *N_pre_*, preparations (slices); SEM reflects the variance among *N_pre_* = 5 preparations; *p < 0.05 (grey segments, intervals of significant difference between the curves). (C) Diagram, extracellular immobilisation of the ratiometric glutamate sensor bFLIPE600n (Venus and ECFP attachments indicated) via biotinylation and attachment to streptavidin (SA, Figure S5C-D; Methods) in *s. radiatum* (pressurized delivery pipette shown). (D) Experimental design: fEPSPs evoked by electrical stimulation of Schaffer collaterals (stim) are recorded with a sensor-injecting pipette (field) while bFLIPE600n fluorescence is monitored within an adjacent ROI (white rectangle). (E) Example of extrasynaptic glutamate transients reported by bFLIPE600n (*ΔR*, ECFP/Venus intensity ratio) in response to Schaffer collateral HFS (100 Hz for 1 s, red arrow; 10 µM NBQX, 50 µM D-APV in the bath) in *stratum radiatum* (input-output calibration, Figure S5E). (F) The fEPSP slope relative to baseline (%, mean ± SEM) in control (green, *n =* 8 slices), during LTP induction (*n* = 14, orange), and with APV present (*n =* 7, orange empty); ***p < 0.001, difference over 20-25 min post-induction. (G) Traces, bFLIPE600n response to two afferent stimuli (50 ms apart, arrows) in control (green) and ∼25 min after LTP induction (orange). Plot, statistical summary (notations as in F); **p < 0.01, difference between LTP and either control or APV data sets.

### Induction of LTP extends extracellular exposure of released glutamate

To test whether the LTP-associated withdrawal of the GLT-1-containing PAPs indeed implies increased extracellular travel of released glutamate, we employed the optical glutamate sensor FLIPE600n (Okumoto et al., 2005). The sensor was modified so that it could be immobilised in the extracellular space (Okubo et al., 2010), as described previously (Whitfield et al., 2015) (Figure 5C, Figure S5D; Methods). The sensor was highly sensitive to glutamate, *in vitro* and *in situ* (Supplementary Figure 5E) and could be delivered to CA1 *stratum radiatum* using a recording patch-pipette (Figure 5C-D). Burst stimulation of Schaffer collaterals induced a clear, stimulus strength-dependent optical FLIPE600n response (Figure 5E, Figure S5F). This response was significantly increased after LTP induction (Figure 5F-G). In similar settings, LTP induction has been reported to have no effect on the amount of released glutamate (Diamond et al., 1998; Luscher et al., 1998). Therefore, the increased bFLIPE600n response suggests a greater exposure of the sensor to the extrasynaptic glutamate transient. To test this at the level of individual synapses we carried out two complementary experiments involving optical glutamate sensors.

### LTP induction widens spatial extracellular transients of released glutamate

In the first experiment, we expressed the glutamate sensor iGluSnFR (Marvin et al., 2013) in hippocampal area CA1, as described previously (Jensen et al., 2019), in either neurons or astroglia (Methods). We confirmed that optical responses of iGluSnFR faithfully reflected both synaptic field responses and changes in their paired-pulse ratios (PPRs, Supplementary Figure 6A-B). In these tests, the PPR was not affected by LTP induction (Supplementary Figure 6C), suggesting that the average release probability remained unchanged, in line with previous reports (Diamond et al., 1998; Luscher et al., 1998).

To probe extracellular glutamate escape from a sub-microscopic point-source (akin to synapses), we monitored fluorescent responses of iGluSnFR to 2PE glutamate spot-uncaging, either near a CA1 pyramidal cell dendrite (Figure 6A) or within an imaged astrocyte territory (Figure S6A). The spatial spread (FWHM) of iGluSnFR responses to 1 ms uncaging pulses was recorded using linescan mode (Figure 6B, Figure S6F), before and 10-30 min after the spot-uncaging LTP induction protocol described above (Figure 4A-B). We found that LTP induction led to the widening of the spatial iGluSnFR signal (Figure 6C-D; *n =* 12). Importantly, the signal spread was unaffected by LTP induction when the visualised astroglia was dialysed whole-cell with the NKCC1 blocker bumetanide (20 µM; Figure 6D; Figure S6F-G).

**Figure 6.**
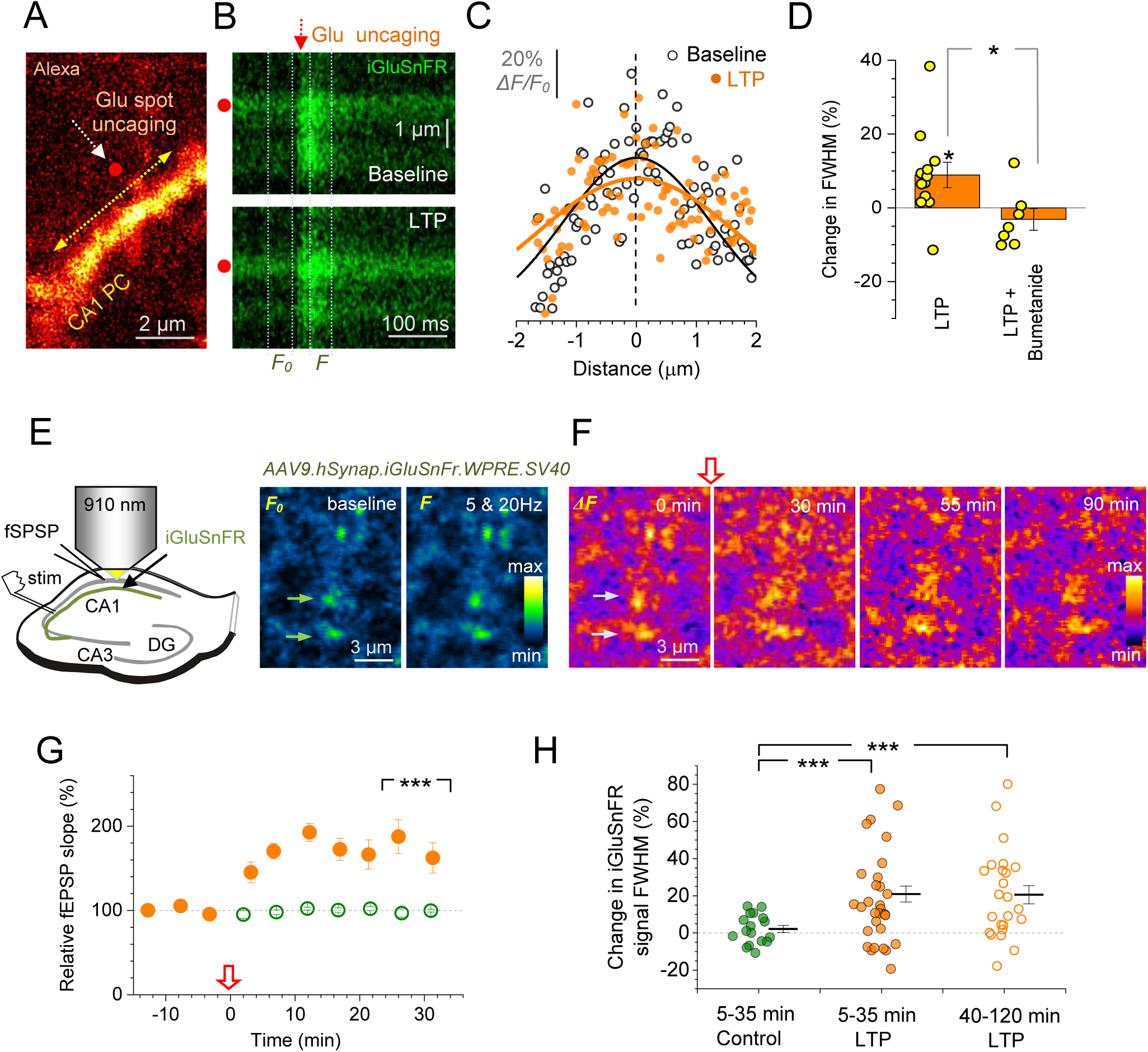
LTP induction expands synaptic glutamate transients in the extrasynaptic space. (A) Dendritic fragment, CA1 pyramidal cell (2PE, AF 594 channel); red dot, glutamate uncaging spot; yellow arrow, line scan positioning for iGluSnFR monitoring; see Figure S6A-C for iGluSnFR validation. (B) Line scans (positioned as in A; iGluSnFR channel) showing extracellular fluorescence transients in response to 1 ms glutamate spot-uncaging (arrow, onset; red dot, position), before (*top*) and 20-25 min after spot-uncaging LTP protocol (*bottom*); dotted lines, sampling time windows for baseline (*F_0_*) and evoked (*F*) fluorescence profiles, giving signal profile *ΔF* = *F - F_0_* (Methods). (C) Spatial iGluSnFR fluorescence profiles (dots, pixel values) from test in (B); zero, uncaging spot position; black and orange lines, best-fit Gaussian approximation. (D) Summary of experiments in (A-C): relative change (%, mean ± s.e.m) in the full-width-at-half-magnitude (FWHM) of glutamate signal, at ∼25 min after LTP induction (LTP, 9.0 ± 3.4%; *n =* 12; *p = 0.027) and with 20 µM bumetanide inside astroglia (LTP+Bumetanide; −3.1 ± 3.0%; *n =* 7; *p < 0.02, difference with LTP data); dots, individual tests. (E) Experiment diagram: monitoring afferent stimulation-evoked glutamate release from Schaffer collateral axonal boutons transduced with iGluSnFR, in acute slices. Image panels (*stratum radiatum* ROI): iGluSnFR fluorescence *F_0_* in resting conditions (left) and fluorescence *F* averaged over five pulses at 20 Hz (right); arrows, two tentative axonal boutons (which show a distinct evoked glutamate signal), false colour scale. (F) Time-lapse frames showing the landscape of the evoked iGluSnFR signal (*ΔF = F - F_0_*; ROI as in E) just before (0 min, time point shown in E) and at three time points after LTP induction (red arrow, onset); false colour scale (see Figure S6H for signal measurement procedure). (G) Time course of fEPSP slope relative to baseline (%, mean ± SEM, *n =* 8 slices) in experiments shown in (E-F); red arrow, HFS LTP induction onset; ***p < 0.001 (difference with no-HFS control, *n =* 4; over 25-35 min post-induction). (H) Average change in the FWHM of evoked axonal iGluSnFR signals, over 5-35 min (*n =* 31 boutons) and 40-120 min (*n =* 21) after LTP induction, as shown; dots, individual experiments; bars, mean ± s.e.m; ***p < 0.001 (difference with no-HFS control group, *n =* 17, recorded 5-35 min post-induction).

To test whether we can detect similar phenomena near active, glutamate-releasing axonal boutons (rather than near the glutamate uncaging spot), in the second experiment we took advantage of slices with relatively sparse expression of iGluSnFR in *stratum radiatum*. Here, we monitored individual (tentative) presynaptic boutons that showed fluorescent iGluSnFR responses to remote electric stimulation of Schaffer collaterals (five pulses 50 ms apart; Figure 6E). In this test, we recorded the spatial spread of the evoked iGluSnFR signal, before and up to 90-120 min after LTP induction (Figure 6F; Figure S6H). As before, LTP induction led to a significant increase in the spatial iGluSnFR signal FWHM, for up to 120 min (Figure 6H). Notably, a proportion of synapses did not show such changes (Figure 6H): these synapses probably reflected non-potentiated connections. In any case, the overall effect was markedly larger than that under spot-uncaging (Figure 6D), likely because a brief train of stimuli amplified glutamate escape (Lozovaya et al., 1999). These results lend further support to the hypothesis that LTP induction boosts extrasynaptic glutamate escape from potentiated synapses, consistent local withdrawal of GLT-1-containing PAPs.

### Whisker stimulation induced LTP reduces PAP presence near firing axons

To understand better the physiological relevance of our observations, we turned to tests in a living animal. We focused on the well-established protocol of LTP induced at the thalamocortical synapses of the barrel cortex (layer *II/III*) by contralateral rhythmic whisker stimulation (RWS) (Gambino et al., 2014; Megevand et al., 2009). Building upon our previous *in vivo* imaging protocols (Mishra et al., 2016; Reynolds et al., 2018; Savtchenko et al., 2018; Zheng et al., 2015), we used viral transduction to express the green Ca^2+^ indicator (hSyn) GCaMP6f in the ventral posteromedial nucleus (VPM) that sends axonal projections to the barrel cortex (Figure 7A). In parallel, barrel cortex astroglia were transduced to sparsely express the red-shifted, cytosol-soluble indicator (GfaABC1D) tdTomato (Figure 7B). This enabled us to monitor, through an implanted cranial window, fine astroglial morphology as well as Ca^2+^ dynamics in individual axonal boutons representing thalamocortical projections (Figure 7C-D).

**Figure 7.**
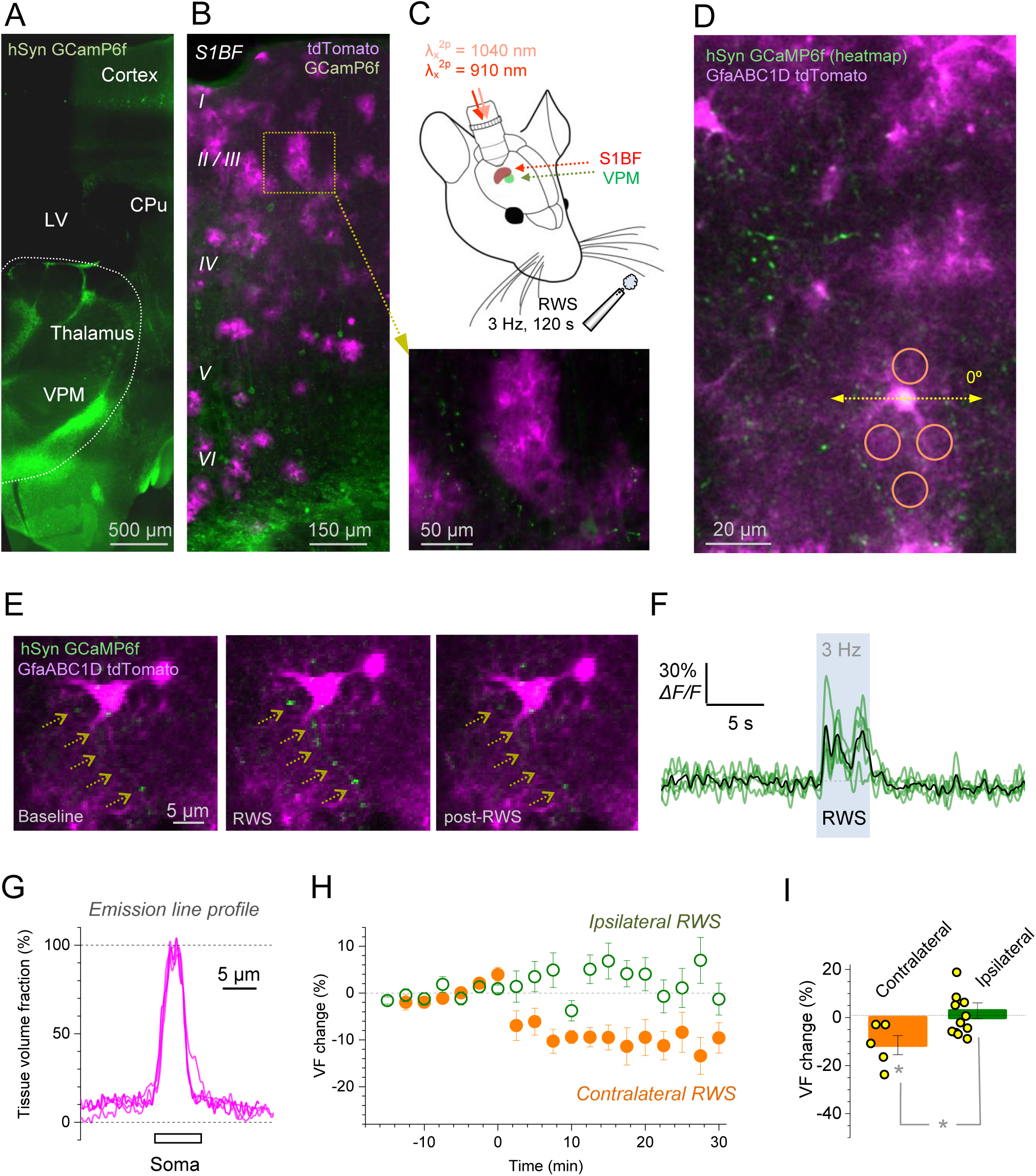
Whisker-stimulation LTP protocol in the barrel cortex *in vivo* triggers volume reduction in astroglia trespassed by stimulated axons. (A) Expression of the Ca^2+^ indicator GCaMP6f three weeks post-transfection (AAV9) into the mouse ventral posteromedial nucleus (VPM), coronal section shown; LV, lateral ventricle; CPu caudate putamen; wide-field image in fixed tissue. (B) Composite image, barrel cortex area (coronal section), with astroglia expressing GfaABC1D tdTomato (magenta; AAV5 transfection) and neuronal structures expressing GCaMPf6 (green); dotted rectangle and magnified inset (yellow arrow), two astrocytes (magenta) with axonal bouton projections occurring nearby (green); λ_x_^2p^ = 1040 nm (tdTomato) and λ_x_^2p^ = 910 nm (GCamPf6). (C) Experiment diagram: 2PE imaging of the barrel cortex (S1BF) through a cranial window, with two fs laser as indicated. The LTP induction protocol uses RWS (5 Hz air-puff stimuli for 120 s) on the contralateral side. (D) Barrel cortex view (S1BF) through the cranial window (λ_x_^2p^ = 1040 and 910 nm, single focal section). Green (GCaMPf6), heat map of axonal signals firing in response to whisker stimulation; magenta (tdTomato), local astroglia. Dashed arrow, sampling tdTomato emission intensity profile (astroglia VF readout; see D below); circles: ROIs for PAP VF readout in proximity to whisker-responding axons (green). (E) Example of individual axonal boutons in S1BF (dashed arrows; GCaMP6f, green) crossing local astroglia (tdTomato, magenta) while responding to a short RWS burst (3 Hz, 5s) with Ca^2+^ elevations. (F) Time course of Ca^2+^ signal (GCaMP6f) at five individual axonal boutons (green traces) shown in (E); black trace, average. (G) Example, astroglial VF profile along the sampling line shown in D (0°), and also at 45°, 90°, and 135° (tdTomato, relative to somatic signal). The profile is similar to that in astroglia loaded whole cell with AF (Figure S1B). (H) Time course of PAP VF change (%, mean ± SEM), during RWS LTP induction protocol, in astroglia crossed by the axons responding to contralateral RWS (orange, *n =* 5 cells, three animals), and in control astroglia during ipsilateral RWS (*n =* 12 cells, four animals). (I) Summary of experiments (shown in G): VF change (%, mean ± SEM) over 15-30 min after the LTP induction onset; dots, data from individual cells; *p < 0.04.

First, we confirmed that the readout of PAP VF using tdTomato was similar to that using whole-cell dialysis of AF (Figure 7D and G; compare to Figure S1A-B). Next, we identified axonal boutons that occurred within the territory of (’trespassing’) tdTomato-expressing astrocytes and responded to a brief RWS train by Ca^2+^ elevations (Figure 7E-F). This enabled us to monitor PAP VF near active axonal boutons, before and after LTP induction by RWS (3 Hz air stimuli, 100 ms pulse width, for 120 s, Figure 7C). We found that LTP induction by contralateral RWS led to a reduction in PAP VF in astroglia trespassed by active boutons whereas applying the same protocol at the ipsilateral side had no effect (Figure 7H-I).

We took advantage of this imaging design to carry out a complementary test in acute hippocampal slices. We dialysed a CA3 pyramidal cell with the Ca^2+^ indicator OGB-1 and traced its axon into area CA1, which was populated with tdTomato-expressing astroglia (Supplementary Figure 7A-B). We then paired presynaptic spikes (triggered by somatic depolarization pulses) with the repetitive brief periods of postsynaptic CA1 pyramidal cell depolarisation induced by an extracellular electrode (Supplementary Figure 7C-D). This LTP-inducting pairing protocol reduced PAP VF near activated axonal boutons by 12 ± 2% (*n =* 5) compared to control. The VF reduction did not occur in the areas that were away from the firing axon (change 3.4 ± 1%, *n =* 10; difference at p < 0.01; Figure S7E).

### LTP induction prompts NMDAR-mediated cross-talk among synapses

If LTP induction enhances extrasynaptic glutamate escape it should also boost activation of high-affinity receptors, such as NMDARs, further away from the synaptic cleft, including neighbouring synapses. Multiple studies have shown that this may have far-reaching implications for synaptic signal integration in local circuitry, and ultimately behaviour (see Introduction). To test whether LTP induction indeed enhances the NMDAR-mediated inter-synaptic cross-talk, we used a protocol that was established specifically to monitor such cross-talk among independent CA3-CA1 synapses (Scimemi et al., 2004). It takes advantage of the use-dependent NMDAR inhibitor MK801, which blocks the receptor only upon its activation. Therefore, if NMDARs at non-active (silent) synapses get progressively blocked by MK801 they must have been activated by glutamate molecules escaping from nearby active synapses.

First, we employed paired-pulse testing to establish presynaptic independence of two Schaffer collateral pathways (Figure S8A). Second, we recorded baseline AMPA receptor-mediated EPSCs (AMPAR EPSCs), then baseline NMDAR EPSCs (in the presence of NBQX) in both pathways (Figure 8A). Third, we applied MK801 and recorded NMDAR EPSCs in one (active) pathway only, while keeping the other pathway silent (Figure 8A). When stimulation resumed in the silent pathway, its NMDAR EPSCs were close to their baseline amplitude (Figure 8A, top dotted line; Figure S8B, *no-LTP, silent*). Thus, the silent pathway had little cross-activation of its NMDARs by synaptic discharges in the active pathway.

**Figure 8.**
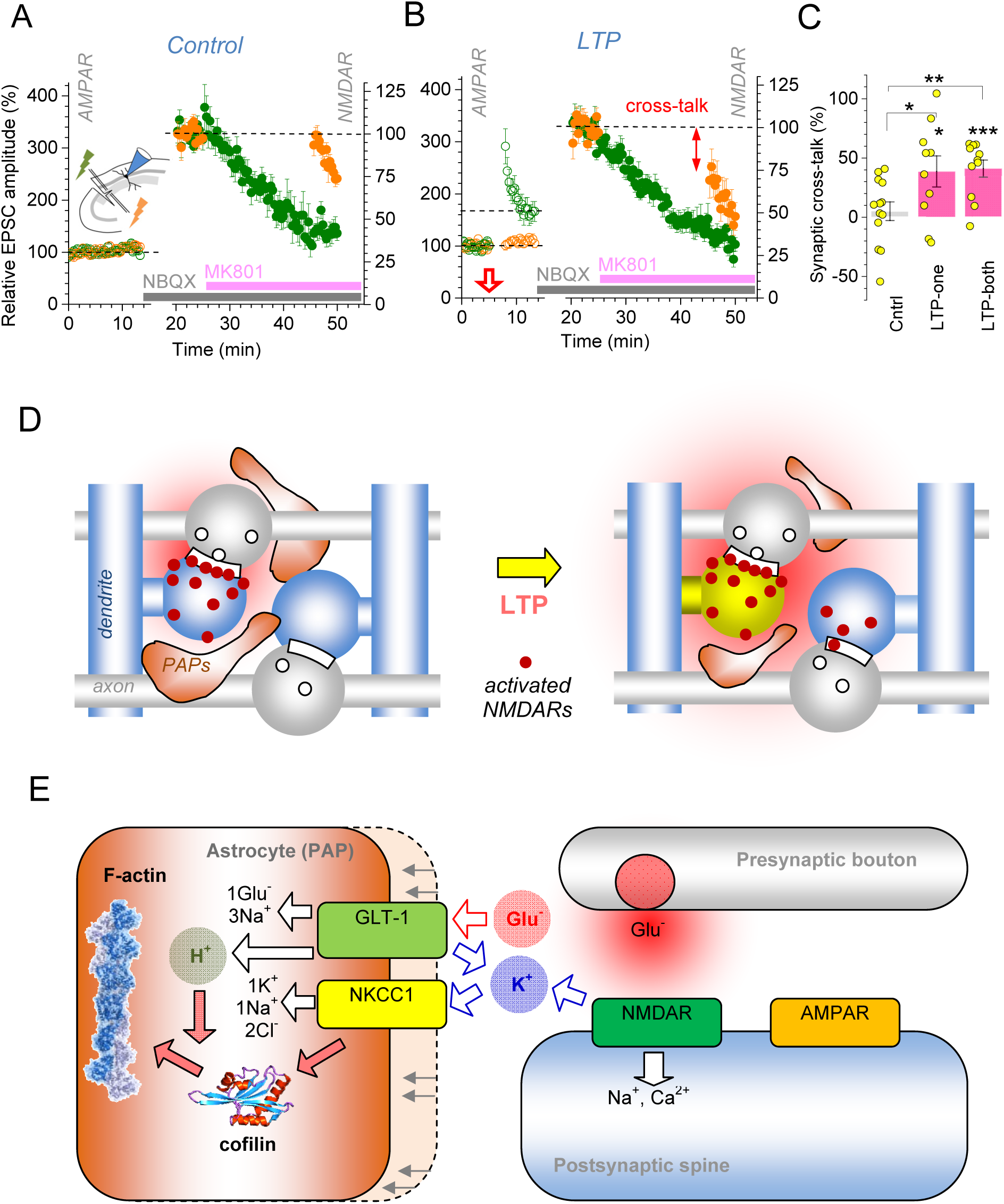
LTP induction boosts NMDAR-mediated inter-synaptic cross-talk. (A) *Inset diagram*, a previously established test (Scimemi et al., 2004) for NMDAR-mediated cross-talk between two independent CA3-CA1 afferent pathways (green and orange lightning) (Methods; Figure S8A). *Plot*, time course of EPSC amplitude (mean ± SEM, *n =* 13), with single stimuli, 20 s apart, applied alternately to the two pathways (green and orange). AMPAR EPSCs are recorded for 12-15 min (V_m_ = −70 mV; left ordinate), then NMDAR EPSCs for ∼5 min (10 µM NBQX, V_m_ = −20 mV; right ordinate). Once MK801 is added, NMDAR EPSCs are recorded in active (green) pathway only. Resuming stimulation in the silent (orange) pathway shows little change in the NMDAR EPSC amplitude compared to baseline (dotted line). (B) Experiment as in (A) but with LTP induced in the active pathway (red arrow; *n =* 7). The reduced amplitude of NMDAR EPSCs in the silent (orange) pathway upon resumed stimulation (arrow, cross-talk) indicates increased NMDAR activation by glutamate escaping from the active (green), potentiated pathway. (C) Summary of experiments shown in (A-B). The extent of cross-talk (percentage of NMDARs in one pathway activated by glutamate escaping from the other pathway; mean ± SEM) is shown, in control conditions (Cntrl, *n =* 13), with LTP induced either in one (LTP-one, *n =* 10) or both (LTP-both, *n =* 11; Figure S8C-D) pathways, prior to NMDAR EPSC recordings; *p *<* 0.05, **p *<* 0.01, ***p *<* 0.001. (D) The proposed scenario of changes in the PAP microenvironment after LTP induction. In baseline conditions (*left*), PAPs restrict glutamate action to the synaptic cleft and some extrasynaptic NMDARs (red dots). Following LTP induction (*right*), some PAPs withdraw, widening the pool of activated extrasynaptic NMDARs, including neighbouring connections. (E) Diagram, a candidate cellular mechanism of LTP-driven PAP withdrawal. LTP induction involves release of glutamate, activating postsynaptic NMDARs and engaging astroglial GLT-1 transporters. This generates a local extracellular potassium hotspot, activating the NKCC1 - cofilin-1 pathway that engages, in a pH-sensitive manner (probably influences by GLT-1 activity), actin polymerisation responsible for morphogenesis. See Discussion for detail.

The result was different when we induced LTP of AMPAR EPSCs in the active pathway (Figure 8B, left ordinate). Here, resuming stimulation of the silent pathway revealed significantly reduced NMDAR EPSCs (Figure 8B, double headed arrow). Thus, a proportion of NMDARs in the silent pathway must have been activated by glutamate escaping from synapses in the active pathway (see Discussion for quantitative estimates). LTP induction in the silent pathway or in both pathways produced a similar boost of NMDAR-dependent inter-pathway cross-talk (Figure 8C, Figure S8C). We confirmed that the trial-to-trial decay of the NMDAR EPSC amplitude was similar among potentiated and non-potentiated pathways, suggesting no effects of LTP induction on the average release probability (Figure S8D), as reported here (Figure S6A-C) and earlier (Manabe and Nicoll, 1994).

## DISCUSSION

This study was prompted by multiple findings highlighting the important role of variable extrasynaptic glutamate escape (spillover) in controlling signal integration in brain circuits (see Introduction). Based on converging lines of evidence, our study unveils a novel regulatory mechanism of synaptic function where LTP induction prompts local withdrawal of PAPs, which in turn boosts extrasynaptic glutamate escape thus increasing activation of NMDARs further away from the release site, potentially involving neighbouring synapses (Figure 8D). To assess whether this sequence of events is biophysically plausible, we employed a detailed Monte-Carlo model of CA3-CA1 synapses (Figure S8E) (Zheng et al., 2008) and simulated three ‘competing’ scenarios that reflect our empirical observations. In these scenarios, the GLT-1 enriched PAPs (i) withdraw with the same numbers of GLT-1, (ii) withdraw while losing some GLT-1, or (iii) re-arrange laterally with the same amounts of GLT-1 (Figure S8F), thus partly exposing extrasynaptic NMDARs. After multiple test runs (example in Video 2), it appeared that scenario (i) was most likely to boost extrasynaptic NMDAR activation (Figure S8G), compared to the other two cases. This is indeed in line with the most parsimonious explanation of our experimental findings.

### Cellular mechanisms of LTP-dependent PAP withdrawal

We found that the PAP withdrawal following LTP induction depends on the ion exchanger NKCC1, a key morphology regulator in brain cell migration (Garzon-Muvdi et al., 2012; Haas and Sontheimer, 2010). In glioma cells, NKCC1 mediates hydrodynamic volume changes and thus prompts dramatic morphological transformations that enable invasion of intact brain tissue (Watkins and Sontheimer, 2011). In these cells, NKCC1 activity can accumulate intracellular chloride up to 140 mM, triggering prominent (up to 35%) cellular shrinkage (Habela et al., 2009). In glioblastoma cells, NKCC1 regulates phosphorylation of cofilin-1 (Schiapparelli et al., 2017), which controls, in a pH-dependent manner, actin filament polymerization, and thus formation and retrieval of thin cell processes, such as dendritic spines (Bravo-Cordero et al., 2013; Ethell and Pasquale, 2005). Indeed, interfering with cofilin-1 phosphorylation by dialyzing individual astrocytes with peptide S3 (Aizawa et al., 2001; Liu et al., 2016) completely occluded LTP-induced PAP shrinkage, suggesting a shared molecular pathway. What activates perisynaptic astroglial NKCC1 upon LTP induction remains to be ascertained. One possibility is that activation of postsynaptic NMDARs and local astroglial GLT-1 during LTP induction leads to a sustained efflux (extracellular hotspot) of potassium. Classically, NKCC1 transport is activated by excess of external potassium (Russell, 2000) whereas proton transport by the GLT-1 transporter could help to boost cofilin-dependent actin assembly. While this provides a plausible explanation (Figure 8E), a full understanding of the cellular mechanisms engaging NKCC1 and cofilin in astroglia will require a separate study.

### 3D EM: faithful representation of live tissue?

The suitability of EM analyses in fixed brain tissue has recently been questioned in elegant tests showing that chemical fixation *in vivo* may produce drastic tissue shrinkage (∼18% linear contraction), with the extracellular space shrinking from ∼15% to ∼2% of tissue volume, which severely distorts PAP morphology (Korogod et al., 2015). However, different chemical fixation protocols produce different outcomes. Our earlier studies reported 5-6% linear tissue shrinkage upon fresh hippocampal slice fixation by submersion (Rusakov et al., 1998), and in another test the estimated extracellular space fraction in area CA1 was ∼12% (Rusakov and Kullmann, 1998), only slightly smaller than the 15% measured in live tissue (Sykova and Nicholson, 2008). In chemically fixed CA1 tissue, astroglia occupied ∼9% of tissue volume (Lehre and Rusakov, 2002), which if anything is towards the higher end of 5-10% (depending in the inclusion of somata and large processes) estimated here with live VF monitoring. Finally, in the correlational EM studies employing rapid hippocampal slice fixation, PAP VF values in the dentate gyrus were undistinguishable between fixed-tissue EM and live imaging data (both at ∼8%) (Medvedev et al., 2014). Here, we used rapid slice fixation by submersion, and made no attempts to assess PAP shapes or describe their exact position. This, in addition to the protocol differences, might explain an apparent discrepancy with some previous studies: for instance, smaller PAPs that occur closer to synapses might well count as an increase in PAP occurrence (Lushnikova et al., 2009; Wenzel et al., 1991) even though their VF decreases.

### PAP withdrawal and extrasynaptic glutamate actions

Changes in PAP geometry on the nanoscale should not affect total glutamate uptake by astroglia because all released glutamate molecules will still be rapidly taken up by the same astrocyte. Furthermore, since glutamate escape and local transporter binding occur on the sub-millisecond scale, subtle PAP rearrangement after LTP induction will have no detectable effect on the uptake kinetics, especially when measured at the astrocyte soma (Diamond et al., 1998; Luscher et al., 1998). With the reduced PAP presence, glutamate should dwell longer and travel further in the extracellular space, which is precisely what the present tests employing optical glutamate sensors indicate.

How much would such changes enhance the inter-synaptic cross-talk by glutamate spillover? We examined NMDAR-mediated cross-activation of two independent afferent pathways (converging onto a CA1 pyramidal cell) and found that, following LTP induction, ∼120 stimuli in the active pathway activated ∼40% NMDARs in the silent pathway. At first glance, this suggests that one discharge activates only ∼0.4% of NMDARs at neighbouring synapses. However, these experiments deal with relatively sparse connections because, on average, only 2-3% of CA3-CA1 synapses are activated in either pathway under this protocol (Scimemi et al., 2004). Because the nearest-neighbour inter-synaptic distance in this area is ∼0.5 µm (Rusakov and Kullmann, 1998), 2% of scattered synapses will be separated by 0.5·(0.02^-1/3^) ∼1.8 µm. Increasing the distance from a release site from 0.5 to 1.8 µm will roughly correspond to a >100-fold glutamate concentration drop (over the first 0.5 ms post-release) (Rusakov, 2001; Zheng et al., 2008). Thus, the accumulated cross-talk among 2-3% of synapses over ∼120 trials will probably underestimate the cross-talk between nearest synaptic neighbours following a single discharge. One might argue that in these experiments the two pathways activate two clustered, relatively dense synaptic pools, rather than evenly scattered synapses: if these clusters overlap in the neuropil our data may overestimate spillover. However, the denser (smaller) the clusters the less likely they overlap in space, in which case there will be little chance of any synaptic cross-talk between the pathways, with or without LTP. Thus, the most parsimonious explanation of our observations predicts two overlapped pools of sparsely activated synapses.

The increased exposure of glutamate to the extracellular space after LTP induction might explain, at least in part, why some pioneering LTP studies reported increased extracellular glutamate transients detected with micro-dialysis (Bliss et al., 1986; Errington et al., 2003). This result might also explain the reduced NMDAR EPSC variability during LTP at CA3-CA1 synapses (Kullmann et al., 1996), an enhanced local excitability of pyramidal cell dendrites after LTP induction (Frick et al., 2004), and why LTP at one synapse could lower the NMDAR-dependent LTP induction threshold at the neighbouring synapses (Harvey and Svoboda, 2007). Among other important functional consequences of increased glutamate escape could be a reported boost in dendritic NMDAR-dependent spikes (Chalifoux and Carter, 2011), facilitated plasticity at inactive excitatory connections nearby (Tsvetkov et al., 2004), or increased heterosynaptic depression (Vogt and Nicoll, 1999). In aged animals, increased glutamate spillover in the hippocampus could impair aspects of memory related to spatial reference and fear (Tanaka et al., 2013; Tsvetkov et al., 2004).

Our observations were necessarily limited to 30-90 min after LTP induction. Therefore, they do not preclude the possibility for PAP coverage to re-establish itself on a longer time scale. Indeed, it has been widely acknowledged that the progressive accumulation of synaptic LTP could lead to runway excitation and that synaptic weight re-scaling is more likely to follow LTP induction. By the same token, one might expect a dynamic sequence of use-dependent astroglial remodelling on a longer time scale, which remains an important and intriguing question.

## ACKNOWLEDGEMENTS

This work was supported by the Wellcome Trust Principal Fellowship, European Research Council Advanced Grant, Medical research Council, Biology and Biotechnology Research Council (all UK), BM1001 Cost Action and FP7 ITN EXTRABRAIN Marie Curie Action (European Commission) (D.A.R.); NRW-Rückkehrerpogramm, UCL Excellence Fellowship, German Research Foundation (DFG) SPP1757 and SFB1089 (C.H.); Human Frontiers Science Program (C.H., C.J.J., H.J.); EMBO Long-Term Fellowship (L.B.). We thank J. Angibaud for preparation of organotypic cultures, R. Chereau and J. Tonnesen for technical help with the STED microscope. This work was supported by grants from Marie Curie FP7 PIRG08-GA-2010-276995 (A.P.) and Marie Curie Astromodulation (S.R.); Equipe FRM DEQ 201 303 26519, Conseil Régional d’Aquitaine R12056GG, INSERM (S.H.R.O.); ANR SUPERTri, ANR-13-BSV4-0007-01, Université de Bordeaux, labex BRAIN (S.H.R.O., U.V.N.); CNRS, HFSP, ANR CEXC and France-BioImaging ANR-10-INSB-04 (U.V.N.); FP7 MemStick Project No. 201600 (M.G.S.).The authors declare no conflict of interest.

## AUTHORS CONTRIBUTIONS

D.A.R. and C.H. conceived the study and its research strategies; C.H., L.B., O.K., D.M. and M.K.H. carried out patch-clamp recordings, morphometric studies, glutamate uncaging, and glutamate sensor imaging experiments and analyses; A.P., S.H.R.O. and U.V.N. designed and carried out STED experiments; J.P.R. implemented expression of genetic sensors and labels and carried out *in vivo* experiments and analyses; N.I.M., I.K., and M.G.S. designed and carried out 3D EM studies and analyses; I.S.R., C.J.J., and H.J. designed and provided the modified optical glutamate sensor bFLIPE600n; S.R. performed S3 peptide experiments; J.H. designed and carried out dSTORM studies; K.Z. performed biophysical modelling tests and dSTORM quantification; S.A. performed selected imaging experiments *ex vivo*; T.J. carried out single-axon pairing experiments in slices; O.P.O. and E.A.N. provided expertise and materials pertinent to the AQP4 and pharmacological dissection tests; D.A.R. carried out selected data and image analyses and wrote the paper which was subsequently contributed to by all the authors.

## STAR METHODS

### Animals

All animal procedures were conducted in accordance with the European Commission Directive (86/609/ EEC), the United Kingdom Home Office (Scientific Procedures) Act (1986), and all relevant national (France, Germany) and institutional guidelines. Details on each of the animal models employed are given throughout the text and summarized below. All animals were maintained in controlled environments as mandated by national guidelines, on 12hr light/dark cycles, with food and water provided *ab libitum*.

For *ex vivo* electrophysiology and imaging, a combination of Wistar rats (3 – 5 weeks old, male), Sprague-Dawley rats (3 – 5 weeks old, male), KO and transgenic mice (3 – 5 weeks old, male) were employed. For experiments requiring viral-mediated expression of optical sensors, male and female wildtype C57BL/6 mice (Charles River Laboratories) were injected at 3 - 4 weeks of age with viral vectors and acute slices were obtained 2 – 4 weeks later. AQP KO mice were backcrossed with C57BL/6 mice for five generations before intercrossing to yield KO (-/-) and wildtype (+/+) mice.

For STED microscopy, organotypic hippocampal slice cultures were prepared from 5 – 7 day old Thy1-YFP mice.

For *in vivo* recordings, group-housed male and female wildtype C57BL/6 mice (Charles River Laboratories) were used. Animals served as their own controls through the use of ipsi- and contralateral stimuli as specified below. All animals were injected with viral vectors at 3 – 4 weeks, and cranial windows were implanted 2 weeks later. Imaging was performed at between 6 and 12 weeks of age, at least 3 weeks after injection of viral vectors.

### Preparation of acute slices

350 μm thick acute hippocampal slices were obtained from three- to five week-old male Sprague-Dawley, Wistar rats, wild-type, knockout and transgenic mice (specified below). Slices were prepared in an ice-cold slicing solution containing (in mM): NaCl 75, sucrose 80, KCl 2.5, MgCl_2_ 7, NaH_2_PO_4_ 1.25, CaCl_2_ 0.5, NaHCO_3_ 26, ascorbic acid 1.3, sodium pyruvate 3, and glucose 6 (osmolarity 300-305), stored in the slicing solution at 34°C for 15 minutes before being transferred to an interface chamber for storage in an extracellular solution containing (in mM): NaCl 126, KCl 2.5, MgSO_4_ 1.3, NaH_2_PO_4_ 1, NaHCO_3_ 26, CaCl_2_ 2, and glucose 10 (pH 7.4, osmolarity adjusted to 295-305). All solutions were continuously bubbled with 95% O_2_/ 5% CO_2_. Slices were allowed to rest for at least 60 minutes before recordings started. For recordings, slices were transferred to the submersion-type recording chamber and superfused, at 33-35°C unless shown otherwise. Where required, 50-100 µM picrotoxin and 5 µM CGP52432 were added to block GABA receptors and a cut between CA3 and CA1 was made to suppress epileptiform activity.

### Electrophysiology *ex vivo*

Electrophysiological examination of astrocytes was carried out as previously described (Henneberger et al., 2010; Henneberger and Rusakov, 2012). Briefly, whole-cell recordings in astrocytes were obtained using standard patch pipettes (3-4 MΩ) filled with an intracellular solution containing (in mM) KCH_3_O_3_S 135, HEPES 10, Na_2_- Phosphocreatine or di-Tris-Phosphocreatine 10, MgCl_2_ 4, Na_2_-ATP 4, Na-GTP 0.4 (pH adjusted to 7.2 using KOH, osmolarity 290-295). Cell-impermeable dyes Fluo-4 (200 μM, Invitrogen) and AF 594 hydrazide (20-100 μM) were routinely added to the intracellular solution, unless indicated otherwise. Where specified, bumetanide (20 µM) or S3 peptide fragment (200 μM, Anaspec) was added to the intracellular solution. Passive astrocytes were identified by their small soma size (∼10 μm; visualized in the AF emission channel), low resting potential (below −80 mV without correction for the liquid-junction potential), low input resistance (< 10 MΩ), passive (ohmic) properties and characteristic morphology of the arbor (Figure 1 and Figure S1). Astrocytes were either held in voltage clamp mode at their resting membrane potential or in current clamp. Where specified, the intracellular free Ca^2+^ concentration was clamped to a steady-state level of 50-80 nM by adding 0.45 mM EGTA and 0.14 mM CaCl_2_ to the intracellular solution (calculation by WebMaxChelator, Stanford).

### LTP induction *ex vivo*

Where indicated, an extracellular recording pipette was placed immediately adjacent to the astrocyte under investigation visualized in the AF channel (Figure 1). Synaptic responses were evoked by orthodromic stimulation (100 µs, 20-100 µA) of Schaffer collaterals using either a bipolar or coaxial stimulation electrode placed in the *stratum radiatum* >200 µm away from the recording electrodes. Field EPSPs (fEPSPs) were recorded using a standard patch pipette filled with the extracellular solution. Predominantly AMPAR-mediated fEPSPs (with no NMDAR blockers added) are denoted AMPAR fEPSPs throughout the text. In some experiments, astrocytic EPSCs (a-fEPSCs) or filed EPSPs (a-fEPSPs) were also recorded using the cell patch pipette (Henneberger and Rusakov, 2012): astrocytic readout was fully consistent with extracellular fEPSPs (e.g., Figure S3C). The baseline stimulus intensity was set at ∼50% of the maximal response, stimuli were applied every 30 seconds for at least 10 minutes before LTP was induced using three trains of high-frequency stimulation (HFS, 100 pulses at 100 Hz) 60 seconds apart. The slope of fEPSPs was monitored afterwards for at least 30 minutes. See sections below for LTP induction protocols used in specific experiments, such as through glutamate uncaging or using a ‘chemical cocktail’.

### Two-photon excitation imaging of astroglia *ex vivo*

We used a Radiance 2100 (Zeiss-Biorad), FV10MP (Olympus), Femto3D-RC or Femto2D (Femtonics, Budapest) and a Scientifica imaging system optically linked to femtosecond pulse lasers MaiTai (SpectraPhysics-Newport) or Vision S (Coherent) and integrated with patch-clamp electrophysiology. Once in whole-cell mode, dyes normally equilibrated across the astrocyte tree within 5-10 min. Routinely, in astrocyte morphology time-lapse experiments astrocytes loaded with fluorescence indicators (see above) were imaged in frame mode at a nominal resolution of ∼ 0.1 µm / pixel (512×512 pixels, 25x Olympus objective /NA1.05) in the red emission channel (540LP / 700SP filter; λ_x_^2P^ = 800 nm). To minimize photodamage only a single focal section through the soma (average of three) was acquired at a laser intensity of 3-6 mW under the objective with careful adjustment of the z-position.

### iGluSnFR transduction of hippocampal astroglia and neurons

#### Stereotactic injections: astroglial expression of iGluSnFR

For the expression of the glutamate sensor iGluSnFR (Marvin et al., 2013) in astrocytes, an AAV virus expressing iGluSnFR under a GFAP promoter (AAV1.GFAP.iGluSnFr.WPRE.SV40; Penn Vector Core, PA, USA) was injected bilaterally into the ventral hippocampus. C57BL6/N mice (4 weeks old, Charles Rivers Laboratories) were injected intra-peritoneally with a ketamin/medotomidine anaesthesia (100 and 0.25 mg per kg body weight in NaCl, injection volume 0.1 ml per 10 g body weight, ketamin 10%, betapharm; Cepotir 1 mg/ml, CPPharma). Firstly, the head fur was removed and the underlying skin disinfected. After ensuring that the animal was under deep anesthesia, the head was fixed in a stereotactic frame (Model 901, David Kopf Instruments). After making an incision, bregma was localized. Next, the coordinates for the ventral hippocampus (relative to bregma: anterior −3.5 mm, lateral -/+3 mm, ventral −2.5 mm) were determined and the skull was locally opened with a dental drill. Under control of a micro injection pump (100 nl/min, WPI) 1 µl viral particles were injected with a beveled needle nanosyringe (nanofil 34G BVLD, WPI). After retraction of the syringe, the incision was sutured using absorbable thread (Ethicon). Finally, the anesthesia was stopped by i.p. application of atipamezol (2.5 mg per kg body weight in NaCl, injection volume 0.1 ml per 10 g body weight, antisedan 5 mg/ml, Ventoquinol). To ensure analgesia, carprofen (5 mg/kg in NaCl, injection volume 0.1 ml/20 g body weight, Rimadyl 50 mg/ml, Zoetis) was injected subcutaneously directly, 24h and 48h after the surgery.

#### Stereotactic injections: neuronal expression of iGluSnFR

C57BL/6 mice (3 - 4 weeks of age), male and female, were prepared for aseptic surgery and anaesthetised using isoflurane (5% v/v induction, 1.5 - 2.5% maintenance). The scalp was shaved and disinfected using three washes of topical chlorhexidine. The animal was secured in a stereotaxic frame (David Kopf Instruments, CA, USA) and loss of pedal reflexes was confirmed prior to surgery. Body temperature was maintained at 37.0 ± 0.5 °C using a feedback rectal thermometer and heating blanket. Perioperative analgesics were administered (subcutaneous buprenorphine, 60 µg kg-1, topical lidocaine/prilocaine emulsion, 2.5%/2.5%) before ocular ointment (Lacri-lube, Allergan, UK) was applied to the eyes. A small midline incision was made and superficial tissue resected to expose the skull. A craniotomy of approximately 1 - 2 mm diameter was performed over the right hemisphere using a high-speed hand drill (Proxxon, Föhren, Germany), at a site overlying the medial hippocampus. Stereotactic coordinates were 60 % of the anteroposterior distance from bregma to lambda and 2.5 mm lateral to midline. Upon exposure, a warmed, sterile saline solution was applied to exposed cortical surface during the procedure.

Pressure injections of AAV9 hSyn iGluSnFR (totalling 0.1 - 1 x 1010 genomic copies in a volume not exceeding 200 nL, supplied by Penn Vector Core, PA, USA) were carried out using a pulled glass micropipette stereotactically guided to a depth of 1.3 mm beneath the cortical surface, at a rate of approximately 1 nL sec-1. The total injection volume was delivered in three steps, reducing depth by 100 μm at each step. Once delivery was completed, pipettes were left in place for 5 minutes before being retracted. The surgical wound was closed with absorbable 7-0 sutures (Ethicon Endo-Surgery GmbH, Norderstedt, Germany) and the animal was left to recover in a heated chamber. Meloxicam (subcutaneous, 1 mg kg-1) was subsequently administered once daily for up to two days following surgery. Mice were killed by transcardial perfusion with ice-cold sucrose-enriched slicing medium (in mM, 105 sucrose, 60 NaCl, 2.5 KCl, 1.25 NaH2PO4, 26 NaHCO3, 15 glucose, 1.3 ascorbic acid, 3 Na pyruvate, 0.5 CaCl2 and 7 MgCl2, saturated with 95% O2 and 5% CO2) after a 2 - 4 week AAV incubation period and acute hippocampal slices prepared for imaging and electrophysiological recordings as below.

### Viral transduction of astroglial GFP

An AAV virus expressing the astroglial GFP (AAV5.GfaABC1D.Pi.lck-GFP.SV40, supplied by Penn Vector Core, PA, USA) was injected into the cerebral ventricles of neonates. C57BL/6J mice (P0-P1), male and female, were prepared for aseptic surgery and maintained all time while being away from the mothers in a warm environment to eliminate risk of hypothermia in neonates. Intracerebroventricular (ICV) injections were carried out after a sufficient visualization of the targeted area to ensure a proper injection. Viral particles (totalling 5 x 109 genomic copies in a volume 2 µl) were injected using a glass Hamilton microsyringe, 2 µl/hemisphere, at a rate not exceeding of 0.2 µl/s, 2 mm deep, perpendicular to the skull surface, guided to a location approximately 0.25 mm lateral to the sagittal suture and 0.50–0.75 mm rostral to the neonatal coronary suture. Once delivery was completed, microsyringe was left in place for 20-30 seconds before being retracted. Pups received ICV injections were kept as a group of litters and returned to the mother in their home cage.

### Viral transduction of thalamocortical boutons and astrocytes in the barrel cortex

C57BL/6 mice (3 - 4 weeks of age), male and female, were prepared as above for neuronal expression of iGluSnFR. During the procedure, two craniotomies of approximately 1 - 2 mm diameter were performed over the right hemisphere using a high-speed hand drill (Proxxon, Föhren, Germany), at sites overlying the ventral posteromedial nucleus of the thalamus (VPM) and the barrel cortex (S1BF). The entire microinjection into the VPM was completed prior to performing the second craniotomy over S1BF. Stereotactic coordinates for VPM injections were −1.8 mm and 1.5 mm along the anteroposterior and mediolateral axes, respectively. Two injection boluses was delivered at 3.0 and 3.2 mm beneath the dural surface. For S1BF injections, the coordinates were −0.5 mm and 3.0 mm along the anteroposterior and mediolateral axes, respectively, delivering a single bolus at a depth of 0.6 mm. A warmed saline solution was applied to exposed cortical surface during the procedure.

Pressure injections of AAV9 hSyn.GCaMP6f (totalling 1 x 1010 genomic copies in a volume not exceeding 200 nL, supplied by Penn Vector Core, PA, USA) and AAV5 GfaABC1D tdTomato (0.5 x 1010 genomic copies, in a volume not exceeding 200 nL, supplied by Penn Vector Core, PA, USA) were carried out using a glass micropipette at a rate of 1 nL sec-1, stereotactically guided to the VPM and S1BF, respectively, as outlined above. Once delivery was completed, pipettes were left in place for 5 minutes before being retracted. The surgical wound was closed and the animal recovered as outlined above for neuronal expression of iGluSnFr. Meloxicam (subcutaneous, 1 mg kg-1) was administered once daily for up to two days following surgery. Mice were subsequently prepared for cranial window implantation approximately 2 weeks later.

### Cranial window implantation

Mice were prepared for aseptic surgery and secured in a stereotaxic frame as before during the viral transduction procedure. Once secured and under stable anaesthesia (isoflurane, maintenance at 1.5 - 2%), a large portion of the scalp was removed to expose the right frontal and parietal bones of the skull, as well as the medial aspects of the left frontal and parietal bones. The right temporalis muscles were reflected laterally to expose the squamous suture, to facilitate cement bonding during fixation of the cranial window implant. The exposed skull was coated with Vetbond (3M, MN, USA) and a custom-made headplate was affixed over the S1BF. The asSEMbly was then secured with dental cement (SuperBond, Sun Medical Co. Ltd., Japan). Once the bonding agents had cured, the animal was removed from the stereotaxic frame and it’s headplate was secured in a custom-built head fixation frame. A craniotomy of approximately 4 mm diameter was carried out over the right somatosensory cortex, centred over the S1BF injection site. Immediately prior to removal of the skull flap, the surface was superfused with warmed aCSF (in mM; 125 NaCl, 2.5 KCl, 26 NaHCO3, 1.25 Na2HPO4,18 Glucose, 2 CaCl2, 2 MgSO4; saturated with 95% O2 / 5% CO2, pH 7.4). The dura was resected using a combination of 26G needles (tapped against a hard surface to introduce a curved profile), fine-tipped forceps (11252-40, Fine Science Tools, Germany) and 2.5 mm spring scissors (15000-08, Fine Science Tools, Germany), taking care not to penetrate to the pia mater. Once the dura was removed, a previously-prepared coverslip consisting of a 34 mm diameter round coverglass affixed beneath a 4 mm diameter round coverglass (Harvard Apparatus UK, affixed using a UV-curable optical adhesive (NOA61), ThorLabs Inc., NJ, USA) was placed over the exposed cortex. Slight downward pressure was applied to the coverslip using a stereotactically guided wooden spatula that was previously severed and sanded to allow some flexibility and preclude excessive force. The superfusion was discontinued and excess aCSF was removed using a sterile surgical sponge, taking care not to wick fluid from beneath the cranial window. The coverslip was then secured with VetBond and dental cement, sequentially. Once cured, the animal was recovered in a heated chamber and returned to its homecage when ambulatory. Post-operative care was administered as before during the viral transduction procedure.

### Multiphoton imaging *in vivo*

Two-photon excitation was carried out using a wavelength multiplexing suite consisting of a Newport-Spectraphysics Ti:sapphire MaiTai tunable IR laser pulsing at 80 MHz and a Newport-Spectraphysics HighQ-2 fixed-wavelength IR laser pulsing at 63 MHz. The laser lightpaths were aligned (though not synchronised) before being point-scanned using an Olympus FV1000 with XLPlan N 25x water immersion objective (NA 1.05). During imaging, animals were lightly anaesthetised (fentanyl, 0.03 mg kg-1, midazolam, 3 mg kg-1, and medetomidine, 0.3 mg kg-1) and secured under the objective on a custom-built stage via the previously affixed headplate.

Initial acquisitions were performed with both lasers illuminating the tissue at 910 nm and 1040 nm, respectively, in order to locate active thalamocortical boutons in S1BF within the arbor of tdTomato-positive cortical astrocytes. Brief 5 second, 3 Hz pulses of nitrogen were directed at the contralateral whiskers to determine responsive regions of interest. Measurements were performed in L1 and L2/3, at depths of 50 - 150 nm. For bouton recordings, framescans of 4 - 20 Hz were performed, with a pixel dwell time of 2 μs and a mean laser power of 30 mW at the focal plane. Upon identification of suitable astrocytes, we sampled the baseline VF. Except when needed for illustrative purposes, illumination by the tunable IR laser (910 nm, to excite GCaMP6f) was occluded at this stage, in order to limit photobleaching. High-resolution z-stacks, incorporating 1 or more astrocytes, were taken every 2.5 minutes, for 15 - 20 minutes. Z-stacks were 512 x 512 pixels, with a pixel size of 0.25 - 0.5 μm and an interval size of 1.5 - 2.5 μm. Sensory-evoked synaptic potentiation within the barrel cortex was then induced as previously described (Gambino et al,. 2014), via a contralateral rhythmic whisker stimulation (RWS, 120 sec, 3 Hz). Sampling of z-stacks, covering the same cortical area, was continued for 30 - 45 minutes following the RWS. The same regions were sampled again one week later, before and after an ipsilateral RWS, to serve as control VF measurements. To determine VF in vivo, stacks were coded (to blind experimenters) and motion-corrected using MATLAB. Fluorescence values for the astrocytic soma and 2 - 4 ROIs within its arbor, from the same focal plane, were tabulated. Sampling of fluorescence from the primary astrocytic branches was avoided as pilot data indicated that VF changes within such branches was negligible. Values for each ROI were averaged to give cell-specific ratiometric fluorescence values, which were normalized to yield relative changes in VF.

### Monitoring astrocyte tissue volume fraction

Astrocyte tissue volume fraction (VF) was monitored to detect structural changes of fine astrocyte branches smaller than the diffraction limit (200-300 nm for diffraction-limited 2PE imaging). VF was obtained by normalizing the background-corrected fluorescence of the morphological dye AF 594 to somatic values, where 100% of the tissue is occupied by the astrocyte (Figure 1A-B, Figure S1A-C). The VF values obtained with this approach were not affected by dye escape through gap-junctions or hemichannels (Figure S1C).

### Fluorescence recovery after photobleaching (FRAP) experiments

FRAP of AF 594 was used to quantify changes of intracellular diffusivity in astrocytes. Fluorescence recordings were obtained in line-scan mode (500 Hz, line placed quasi-randomly through the astrocyte arbour) at an increased laser power of 15-20 mW under the objective to induce substantial bleaching of AF 594.

### Optical measurements of extracellular diffusivity

The effective diffusivity of fluorescent dyes was determined using a point-source diffusion method as previously described (Savtchenko and Rusakov, 2005; Zheng et al., 2008). Briefly, a bolus of fluorescent dye (AF 594 hydrazide, 50 µM in extracellular solution) was ejected from a patch pipette into the CA1 *stratum radiatum* neuropil by a pressure pulse (0.8 bar, 2-6 ms). The diffusion spread of the dye was traced by scanning along a line in front of the ejection pipette (∼300-1000 Hz; Figure S2A). Fluorescence life profiles for each time point were fitted with a Gaussian function exp((x - xc)^2^/(4 w)) with *w* = *D_eff_ t* where x is the position within the linescan, xc the puff position, *D_eff_* the effective diffusivity and t the time since the puff. *D_eff_* is then obtained by linear fitting of w(t) (Figure S2B). All analyses were performed using Matlab (Mathworks). Measurements were repeated every 10 minutes. Field EPSPs were evoked by Schaffer collateral stimulation (see above) and recorded through another field pipette less than 150 μm away from the puff pipette. In a subset of recordings LTP was induced after 10 minutes of baseline recording.

### STED microscopy in organotypic slices

Organotypic hippocampal slice cultures were prepared from 5-7 day pups of Thy1-YFP transgenic mice in accordance with the French National Code of Ethics on Animal Experimentation and approved by the Committee of Ethics of Bordeaux (No. 50120199). As described before (Nagerl et al., 2004), cultures were prepared using the roller tube method (Gähwiler method). First, pups were decapitated. Then, brains were removed, hippocampus dissected (in cooled Gey’s Balanced Salt Solution, GBSS) and 350 µm coronal slices were sectioned using a tissue chopper (McIlwain). After 30-60 minutes rest at 4°C in GBSS, each half slice was mounted on a glass coverslip coated with heparinized chicken plasma (10 µl, Sigma). Thrombin (Merck) was added to coagulate the plasma and to allow the slice to adhere to the coverslip. After 30 minutes at room temperature, each coverslip was inserted into a delta tube (Nunc) before adding 750 µl culture medium containing: 50% Basal Medium Eagle (BME, Gibco), 25% Hanks’ Balanced Salt solution (HBSS, Gibco), 25% of heat inactivated horse serum (Gibco) supplemented with glutamine to a final concentration of 1mM and glucose to a final concentration of 11g/l (Sigma). Finally, slices were cultivated during 5-6 weeks in tubes placed on a roller-drum incubator set at 35 °C in dry air with a rotation rate of ∼10 revolutions per hour. The experimental day, the slice was transferred to a submersion-type recording chamber perfused (2 ml/min) with ACSF at 31°C saturated with 95%O_2_/5%CO_2_ and containing (in mM): NaCl 119, KCl 2.5, NaH2PO4 1.25, NaHCO3 26, Trolox 1.5 and 10 glucose (pH 7.4; osmolarity 295-298) in the presence of 1.3 mM Mg^2+^ and 2 mM Ca^2+^.

To enable STED microscopy studies, as described previously (Tonnesen et al., 2011), our home-built STED microscope was constructed around the base of an inverted confocal microscope (DMI 6000 CS Trino, Leica, Mannheim, Germany) using a glycerin objective with a high numerical aperture and equipped with a correction color (PL APO, CORR CS, 63x, NA 1,3; Leica), providing an optical resolution of at least 70 nm in x-y tens up to 50 μm below the tissue surface. A pulsed-laser diode (PDL 800-D, Picoquant, Berlin, Germany) was used to deliver excitation pulses at 485 nm wavelength with 90 ps duration. Furthermore, an optical parametric oscillator (OPO BASIC Ring fs RTP, APE, Berlin, Germany) pumped by a Ti:Sapphire laser (MaiTai, Spectra-Physics, Darmstadt, Germany), operating at 80 MHz produced a pulsed STED beam centred at a wavelength of 592 nm, to quench the fluorescence. The maximal power of the STED beam going into the back aperture of the objective was 12 mW. Both, excitation and STED pulses were synchronized at 80 MHz by externally triggering the laser diode and optimizing the relative delay using an electronic delay generator. The fluorescence signal was first separated from the excitation light by a dichroic mirror (499-nm long-pass), then cleaned with a 525/50 band-pass filter, spectrally separated by a dichroic mirror (514-nm long-pass), and finally imaged onto two multimode optical fibers connected to avalanche photodiodes (SPCM-AQR-13-FC, PerkinElmer, Waltham, MA).

Image acquisition was controlled by the custom-written software IMSpector (www.max-planck-innovation.de/de/industrie/technologieangebote/software/). The pixel dwell time was 15 µs with a pixel size of 19.53 nm. Typically, 2 µm stacks, with nine z-sections, 220 nm apart were acquired. As described before (Tonnesen et al., 2011), YFP (in neurons) and AF 488 (in astrocytes) were spectrally detected using a 514 nm long-pass emission filter. Effective colour separation was achieved offline by linear un-mixing of the fluorescence channels (using a plugin from ImageJ) after deconvolution (3 iterations) using Huygens Professional (SVI). All morphometric analyses were done on deconvolved image sections of the two unmixed colour channels. To determine spine head width, a 3-pixel thick line was manually positioned through the largest part of the spine head, and the full width at half maximum (FWHM) as a measure of spine size was extracted from the line profile. Astrocytic processes and spines were considered to be in close proximity if the visible distance between their edges (as determined by the FWHM of a line profile laid across the point of shortest distance) was equal or less than 20 nm, corresponding to one pixel. Conversely, for separations larger than 1 pixel, the astrocytic process and spine were not considered to be in close proximity.

### Fast fixation and DAB staining of recorded astrocytes

In a subset of experiments, we loaded an astrocyte with biocytin, and after the experiment the slices were rapidly fixed (by submersion) with 1.25% glutaraldehyde and 2.5% paraformaldehyde in 0.1 M PB (phosphate buffer, pH 7.4), to be kept overnight, infiltrated in 10% sucrose in PB for 10 min and then in 20 % sucrose in PB for 30 min. Infiltrated slices were consequentially freeze-thaw in liquid freon and liquid nitrogen for 3 sec each to gently crack intracellular membranes and embedded in 1% low gelling temperature agarose in PB (Sigma-Aldrich, USA). Embedded slices were sectioned at 50 µm on a vibrating microtome (VT1000; Leica, Milton Keynes, UK). 50 µm sections were incubated in 1% H2O2 in PB for 20 min to eliminate blood background, washed with 0.1 M TBS (tris buffer saline, pH 7.4) and incubated with ABC solution (VECTASTAIN ABC, Vector laboratories, USA) for 30 min at room temperature. Next section were washed with 0.1M TB (tris buffer, pH 7.4), pre-incubated with DAB (3,3’-Diaminobenzidine tablets - Sigma-Aldrich, USA) solution (10 mg DAB tablet + 40 ml TB) for 30 min at room temperature in dark and finally incubated with DAB+ H2O2 solution (5 μl of 33% H2O2 + 25 ml of DAB solution) for 10-20 min at room temperature in dark. The DAB stained sections were washed in PB, post-fixed in 2% osmium tetroxide and further processing and embedding protocols were essentially similar to those reported previously (Medvedev et al., 2010). Briefly, the tissue was dehydrated in graded aqueous solutions of ethanol (30-100%) followed by 3 times in 100% acetone, infiltrated with a mixture of 50% epoxy resin (Epon 812 ⁄ Araldite M) and 50% acetone for 30 min at room temperature, infiltrated in pure epoxy resin, and polymerized overnight at 80 °C. Sections in blocks were coded and all further analyses were carried out blind as to the experimental status of the tissue.

### 3D electron microscopy

Serial sections (60–70 nm thick) were cut with a Diatome diamond knife as detailed and illustrated earlier (Medvedev et al., 2010; Popov et al., 2005; Popov et al., 2004), and systematically collected using Pioloform-coated slot copper grids (each series consisted of up to 100 serial sections). Sections were counterstained with 4% uranyl acetate, followed by lead citrate. Finally sections were imaged in *stratum radiatum* area of CA1 (hippocampus) using an AMT XR60 12 megapixel camera in a JEOL 1400 electron microscope. Serial sections were aligned as JPEG images using SEM align 1.26b (software available from http://synapses.clm.utexas.edu/). 3D reconstructions of DAB stained astrocyte processes and the adjacent dendritic spines were performed in Trace 1.6b software (http://synapses.clm.utexas.edu/). Dendritic spines were categorized according to (Harris et al., 1992; Peters and Kaiserman-Abramof, 1970); since 90-95% of excitatory synapses in CA1 area of hippocampus are located on either thin or mushroom dendritic spines only the mushroom (*n =* 88) and thin (*n =* 243) spines were reconstructed and analyzed. 3D reconstructions of segmented astrocytic processes and dendritic spines were imported to 3D-Studio-Max 8 software for rendering of the reconstructed structures.

### Measurements of astroglial coverage in 3D EM

To analyse astroglial coverage of synapses, a set of virtual 100 nm thick concentric spherical shells (Figure 3D) was arranged *in silico* around each reconstructed PSD using custom-made software. The volume of each shell as well as the volume and surface area of astrocytic segments inside each shell were computed to estimate the volume fraction (VF) occupied by astrocyte processes (astrocyte volume / total shell volume) and the surface area of astrocyte, throughout concentric shell between centered at 0-0.5 μm around the centroid of each individual PSD. In some cases, we also carried out additional analyses using curvilinear 3D shells reproducing the contours of each PSD; the results were qualitatively identical. All data from digital reconstructive analyses were evaluated to obtain one value for each individual slice taken from individual animals (there were *n =* 3 preparations in each group), in each data set. ANOVA tests were used to examine differences between specific animal groups (implemented through Origin Pro 7.5). Data were presented as mean ± SEM (*n =* 3 animals per group).

### Immunohistochemistry and three-color 3D dSTORM

We used a modified protocol described by us previously (Heller et al., 2017). Deeply anaesthetized rats (Sprague Dawley, ∼500 g) were perfused with ice-cold 4% PFA in PBS, brains were removed and incubated in 4% PFA in PBS overnight at 4°C; 30 μm coronal sections were prepared and kept free-floating in PBS; non-reacted aldehydes were quenched in 0.1% NaBH_4_ in PBS for 15 min; washed thrice for 5 min with PBS; autofluorescence was quenched with 10 mM CuSO_4_ in 50 mM NH_4_Cl, final pH = 5 for 10 min; washed with H_2_O thrice quickly and once with PBS (5 min). Permeabilisation and blocking was carried out with PBS-S (0.2% saponin in PBS) supplemented with 3% BSA for at least 3 hours; incubated with primary antibody (see below) in PBS-S overnight at 4°C; washed trice with PBS-S; incubated with secondary antibody (see below) in PBS-S for two hours; washed with PBS-S twice for 10 min and with PBS twice for 10 min; post-fixed with 4% PFA in PBS for 30 min; washed with PBS thrice for 10 min; incubated in Scale U2 buffer (Hama et al., 2011) (4 M urea, 30% Glycerol and 0.1% Triton X-100 in water) at 4°C until being prepared for imaging.

Primary antibodies were for: presynaptic protein Bassoon (Mouse, SAP7F407, Recombinant rat Bassoon, Novus, NB120-13249, AB_788125, dilution 1:500), postsynaptic protein Homer1 (Rabbit, polyclonal, Recombinant protein of human homer (aa1-186), Synaptic Systems, 160003, AB_887730, dilution 1:500), glial glutamate transporter GLT-1 (Guinea pig, Polyclonal, Synthetic peptide from the C-terminus of rat GLT-1, Merck, AB1783, AB_90949, dilution 1:500). Secondary antibodies were: anti-mouse IgG (Donkey, CF568-conjugated, Biotium, 20105, AB_10557030, dilution 1:500), anti-rabbit IgG (Goat, Atto488-conjugated, Rockland, 611-152-122S, AB_10893832, dilution: 1:500), anti-guinea pig IgG (Donkey, Alexa647-conjugated, Jackson ImmunoResearch Labs, 706-606-148, AB_2340477, dilution: 1:500).

To obtain spatial patterns of individual proteins in the synaptic microenvironment, we employed the single-molecule localization microcopy (SMLM) technique direct stochastic optical reconstruction microscopy (dSTORM) (Endesfelder and Heilemann, 2015). Images were recorded with a Vutara 350 microscope (Bruker). The targets were imaged using 640 nm (for Alexa647), 561 nm (for CF568) or 488 nm (for Atto488) excitation lasers and a 405 nm activation laser. We used a photoswitching buffer containing 100 mM cysteamine and oxygen scavengers (glucose oxidase and catalase) (Metcalf et al., 2013). Images were recorded using a 60x-magnification, 1.2-NA water immersion objective (Olympus) and a Flash 4.0 sCMOS camera (Hamatasu) with frame rate at 50 Hz. Total number of frames acquired per channel ranged from 5000-20000. Data were analyzed using the Vutara SRX software (version 6.02.05) and a custom-written script for MATLAB. Single molecules were identified by their continued emission frame-by-frame after removing the background. Identified particles were then localized in three dimensions by fitting the raw data with a 3D model function, which was obtained from recorded bead data sets. The experimentally achievable image resolution is 20 nm in the *x-y* plane and 50 nm in the *z* direction; in tissue sections we routinely achieved *x-y* resolution of 58.0 ± 7.1 and *z-*resolution of 73 ± 5.8 nm.

### Chemical induction of long-term potentiation

The classical ‘chemical’ LTP (cLTP) was induced by perfusing the acute slice for 10-15 min with the Mg-free ACSF solution containing 4 mM CaCl_2_ (Sigma), 0.1 μM rolipram (Cayman Chemical Company), 50 μM forskolin (Cayman Chemical Company) and 50 μM picrotoxin (Sigma) (Otmakhov et al., 2004). This treatment increases the level of cAMP and that of network activity leading to a tetanic-like stimulation in bulk that potentiates the majority of excitatory synapses.

### LTP induction by two-photon spot-uncaging of glutamate

We used a combined two-photon uncaging and imaging microscope (Olympus, FV-1000MPE) powered by two Ti:Sapphire pulsed lasers (Chameleon, Coherent, tuned to 720 nm for uncaging and MaiTai, Spectra Physics, tuned to 840 nm for imaging). The intensity of the imaging and uncaging laser beams under the objective was set to 5 mW and 12-17 mW, respectively. CA1 pyramidal neurons and astrocytes were loaded with Fluo-4 (200 µM) and AF 594 (100 µM) and held in current-clamp mode. The MNI-glutamate was applied in the bath at 2.5 mM. The stimulation protocol was delivered >30 µm from the cell soma and included three series of 100 x 1ms pulses at 100Hz, 60 seconds apart. The uncaging spot was placed ∼1µm from the identifiable small process in astrocytes or the dendritic spine head in patched and visualized CA1 pyramidal neurons.

To test whether this protocol elicited LTP, CA1 pyramidal neurons were recorded in whole-cell patch clamp (see above), and EPSCs were elicited by 1 ms uncaging pulses delivered every 3 min. After a 10 min baseline, the neuron was held in current clamp (−60 to −65 mV, as in freely-moving rats (Epsztein et al., 2010)) and LTP was induced using the glutamate uncaging protocol. Once the induction protocol had been completed, EPSCs were monitored in voltage clamp for 30 min.

For IP_3_ uncaging, 400 µM NPE-caged IP_3_ (D-Myo-Inositol 1,4,5-Triphosphate, P4(5)-(1- (2-Nitrophenyl)ethyl) ester, Life Technologies) were added to the internal solution. The uncaging protocol consisted of 3-5 cycles (200 ms apart) of 5-10 ms pulses on 4-5 points, repeated 3 times every 60 s. To test the effect of glutamate and IP_3_ uncaging on astrocyte morphology, astrocytes located in the *stratum radiatum* of CA1 were loaded with Fluo-4 (200 µM) and AF 594 (100 µM).

In baseline conditions and 30-40 min after the glutamate-uncaging LTP induction protocol, Z-stacks of the same region of the astrocyte were collected every 60-120 seconds. The intracellular Ca^2+^ response to glutamate and IP_3_ uncaging was recorded using frame-scans in astrocytes (Figs. 4A, 5E) and linescan recordings in dendritic spines of CA1 pyramidal cells and expressed as Δ*G/R* values (green/red ratio; Fluo-4 fluorescence normalized to the AF 594 signal, Figure S5A-B).

### Probing ephrins and extracellular matrix signalling

The candidate morphogenic signals that could be invoked during LTP induction involve signaling molecules of the extracellular matrix (ECM) (Dityatev and Rusakov, 2011) or the ephrin/Eph-dependent neuron-astrocyte signaling attributed to astrocyte-dependent stabilization of newly formed dendritic protrusions (Nishida and Okabe, 2007). The protocol for catalytic removal of chondroitin sulfate (and side chains of proteoglycans) with Chondroitinase ABC (0.5U/ml, 45 min, 33°C) has been established and validated by us previously (Kochlamazashvili et al., 2010). Similarly, the blockade of EphA4 activity with EphA4-Fc (10 µg/ml) using previously tested protocols was carried out in accord with the reported procedures (Murai et al., 2003). Because degrading the ECM’s hyaluronic acid with hyaluronidases interfered with LTP induction (Kochlamazashvili et al., 2010) such experiments were not included in the present study. Suppressing NKCC1 activity in the recorded astrocyte was performed through intracellular dialysis of bumetanide (20 µM) (Migliati et al., 2009).

### Monitoring extracellular glutamate transients with optical glutamate sensors

We modified FLIPE600n (Okumoto et al., 2005) to contain a biotin tag for immobilization of the sensor in the tissue, as described previously (Whitfield et al., 2015). A nucleotide sequence coding for the biotin tag was synthetized de novo (Epoch Life Science), amplified using PCR and then inserted into pRSET FLIPE-600n (Addgene #13537, courtesy of Wolf B. Frommer) using BamHI restriction site.

bFLIPE600n reports glutamate levels through a FRET mechanism, by changing the fluorescence intensity ratio *R* = ECFP/Venus. Calibration of the bFLIPE600n sensor using 2PE was first done in free solution (Figure S5A-B). bFLIPE600n in PBS (3-4 µM, pH 7.4) was placed in a meniscus under the microscope objective. Increasing amounts of glutamate (dissolved in PBS, pH adjusted to 7.4) were added and changes in the ECFP/Venus emission ratio were calculated offline. For experiments in acute slices, 30-40 µM bFLIPE600n were preincubated with 5-7 µM streptavidin in PBS at 4° C for at least 12 h. A standard patch pipette (2-4 MΩ resistance) was then backfilled with the sensor solution and bFLIPE600n was gently injected into the CA1 s*. radiatum* of biotinylated slices (see above and (Whitfield et al., 2015)) at 70-100 µm depth applying light positive pressure for 10-20 s. Sensor levels were allowed to equilibrate for 15 min before recordings started at a depth of 50-60 µm below the slice surface (∼3 mW excitation intensity at the focal plane). Schaffer collateral stimulation was done as described above except that the stimulation intensity was ∼50% of the one inducing near-maximal fEPSP responses.

### Evaluating the extent of extracellular glutamate transients with iGluSnFR

iGluSnFR was expressed in the CA1 region of the hippocampus as described above. A iGluSnFR-expressing CA1 pyramidal neuron was loaded with 100 µM AF 594 to visualize dendritic spines. The iGluSnFR fluorescence was monitored in linescan mode (λ_x_^2P^ = 910 nm, 500 Hz) following MNI-glutamate uncaging (1 ms pulse, 2.5 mM in the bath). The linescan was positioned near the closest dendritic spines head, parallel to the dendritic stem (Figure 6d). In baseline conditions, three linescan images were recorded 3 min apart and averaged (Figure 6E, top). LTP was induced with 2PE uncaging of glutamate as described above. 5-10 min following LTP induction, five linescan images were recorded every five minutes for averaging (Figure 6E, bottom).

In each linescan image, two ∼30 ms long ROI bands were selected for analyses, one shortly before the spot-uncaging onset (background iGluSnFR fluorescence profile *F_0_* along the linescan axis *x*, *F_0_*(*x,t*)) the and one ∼10 ms after (glutamate-bound iGluSnFR profile *F*(*x,t*); Figure 6E). The pixel brightness values (originally recorded grey scale) in these iGluSnFR linescan images were (i) averaged along the timeline *t*, and (ii) among the pre-uncaging and the post-uncaging groups in each trial, thus giving average profiles *F*_0_*(*x*) and *F\**(*x*), respectively, for trials before and after LTP induction. In each trial therefore the glutamate signal profile was obtained as a pixel-by-pixel image (vector) operation (*F\**(*x*)*-F*_0_*(*x*)) */ F*_0_*(*x*) giving the glutamate-bound iGluSnFR brightness distribution *ΔF/F_0_* (*x*) along a linescan axis near the uncaging spot. The distribution *ΔF/F_0_* (*x*) along *x*-axis (distance) was best-fit approximated with a Gaussian centered at the uncaging spot, with the amplitude *A* and dispersion *σ* being free parameters (OriginPro, Origin Lab Corp, MA).

### Evaluating NMDAR-mediated inter-synaptic cross-talk in a two-pathway experiment

The NMDAR-mediated synaptic cross-talk was probed by taking advantage of the use-dependency of the NMDAR inhibitor MK801, as described in detail earlier (Scimemi et al., 2004). CA1 pyramidal cells where held in voltage clamp to record EPSCs in response to stimulation of two independent synaptic CA3-CA1 pathways (see Figure 4C for an illustration, GABA receptors blocked as described above). While individual pathways displayed a robust (same-pathway) paired-pulse facilitation of 75.4 ± 6.1% (*n =* 54, *P* < 0.001; inter-stimulus interval 50 ms), the facilitation was approximately five times lower between the pathways (16.5 ± 2.9%, *P* < 0.0001) thus indicating that these pathways do not interact presynaptically by more than ∼20%. Separation of pathways was helped by making an additional cut into the *stratum radiatum* in parallel to the pyramidal cell layer. AMPAR-mediated EPSCs were recorded at a holding potential of −70 mV for 10-15 minutes. In a subset of experiments LTP was induced on one or both pathways (HFS, see above). NMDAR-mediated EPSCs of the same pathways were then recorded by clamping the cell to −20 mV and inhibiting AMPAR with 10 µM NBQX. MK801 (4 µM) was bath-applied after another baseline period. Stimulation of the test pathway was then stopped and resumed after 20 minutes. EPSCs were evoked at 0.1 Hz throughout the experiment. Synaptic cross-talk was quantified at the test pathway by calculating the reduction of NMDAR-mediated EPSC amplitudes in the absence of test pathway stimulation relative to baseline.

An LTP-associated increase of the presynaptic release probability (PR) may facilitate cross-talk independent of astrocyte morphology changes. According to the binomial model of release, an increase of PR would decrease the variability of postsynaptic responses (coefficient of variation [CV]). Experiments using LTP induction in a single pathway showed that the CVs for the baseline AMPAR and NMDAR-mediated responses were not different between pathways and within a pathway (1/CV^2^, four paired Student t-tests, p > 0.18). In addition, the rate of blockade of NMDAR-mediated response by MK801 is an indicator PR and was not affected by LTP-induction (Figure S8D).

Recordings were carried out using a Multiclamp 700B (Molecular Devices). Signals were filtered at 3-10 kHz, digitized and sampled through an AD converter (National Instruments or Molecular Devices) at 10-20 kHz, and stored for off-line analysis using pClamp10 software (Molecular Devices). Receptor blockers were purchased from Tocris and Abcam Biochemicals.

### Monte Carlo simulations

#### Monte Carlo simulations of glutamate diffusion, uptake and NMDAR activation in the environment of the CA3-CA1 synapse

The modelling approach was described and validated against experimental data previously (Savtchenko et al., 2013; Zheng et al., 2015; Zheng et al., 2008). In brief, the presynaptic part (Schaffer collateral en-passant boutons) and the postsynaptic part (dendritic spine heads of CA1 pyramidal cells) were represented by the two truncated hemispheres separated by a 300 nm wide 20 nm high apposition zone including a 200 nm wide synaptic cleft (Figure S7), to reflect the typical three-dimensional ultrastructure reported for these synapses (Harris et al., 1992; Lehre and Rusakov, 2002; Shepherd and Harris, 1998; Ventura and Harris, 1999). The synapse was surrounded by 20-30 nm wide extracellular gaps giving an extracellular space fraction α ∼ 0.15. 3000 molecules of glutamate (Savtchenko et al., 2013) were released at the cleft center and allowed to diffuse freely. The diffusion coefficient for glutamate (excluding space tortuosity due to cellular obstacles) was set at 0.4 µm^2^/ms (Zheng et al., 2008). The statistics on activation of extrasynaptic NMDARs were collected using a cluster of receptors placed at 200-250 nm from the synaptic centroid (thus approximately equidistant to the two nearest-neighboring synapses in area CA1 (Rusakov and Kullmann, 1998)). To test four different scenarios pertinent to the astroglial environment of synapses, we distributed glial glutamate transporters (EAAT1-2 type) using four different patterns that occupy four sectors of the extrasynaptic environment (Figure S7). In the control case (baseline conditions) their extracellular density was ∼0.2 mM, to match a membrane surface density of 5-10•10^3^ μm^−2^ (Lehre and Danbolt, 1998) reported earlier. Cases (*i-iii*) thus mimicked possible astroglial re-arrangements following LTP induction. In case (*i*), the transporter density doubled while the astrocyte membrane area occupied by them was reduced two-fold (thus the total transporter number was unchanged); case (*ii*) was similar to (*i*) but with the transporter density unchanged (total number was reduced two-fold); and in the case (*iii*) the transporter-occupied area was rearranged towards one side of the nearby NMDAR cluster. During extensive control simulations we found no interaction between any of the four sectors in terms of transporter or NMDAR activation by released glutamate. In our tests therefore we could compare the four scenarios using the same simulations run (repeated 32 times for a statistical assessment of the stochastic receptor and transporter actions). Our simulations have suggested that, somewhat paradoxically, one factor that could prolong the presence of glutamate near NMDARs and therefore boosting receptor activation could be its stochastic unbinding from local transporters, as suggested earlier (Rusakov, 2001). Simulations were carried out using a dedicated 14-node PC cluster running under Linux (Zheng et al., 2015).

## QUANTIFICATION AND STATISTICAL ANALYSIS

The present study contained no longitudinal or multifactorial experimental designs. In electrophysiological or imaging experiments the main source of biological variance was either individual cells or individual preparations (the latter in case of field measurements in acute slices), as indicated. In accord with established practice, in the *ex vivo* tests we routinely used one cell per slice per animal, which thus constituted equivalent statistical units in the context of sampling, unless indicated otherwise. Statistical hypotheses pertinent to mean comparisons were tested using a standard two-tailed *t*-test, unless the sample showed a significant deviation from Normality, in which case non-parametric tests were used as indicated. The null-hypothesis rejection-level was set at α = 0.05, and the statistical power was monitored to ensure that that the sample size and the population variance were adequate to detect a mean difference (in two-sample comparisons) of 10-15% or less. Group data are routinely reported as mean ± SEM, unless indicated otherwise.

## SUPPLEMENTARY FIGURES

**Figure S1 (related to Figure 1).**
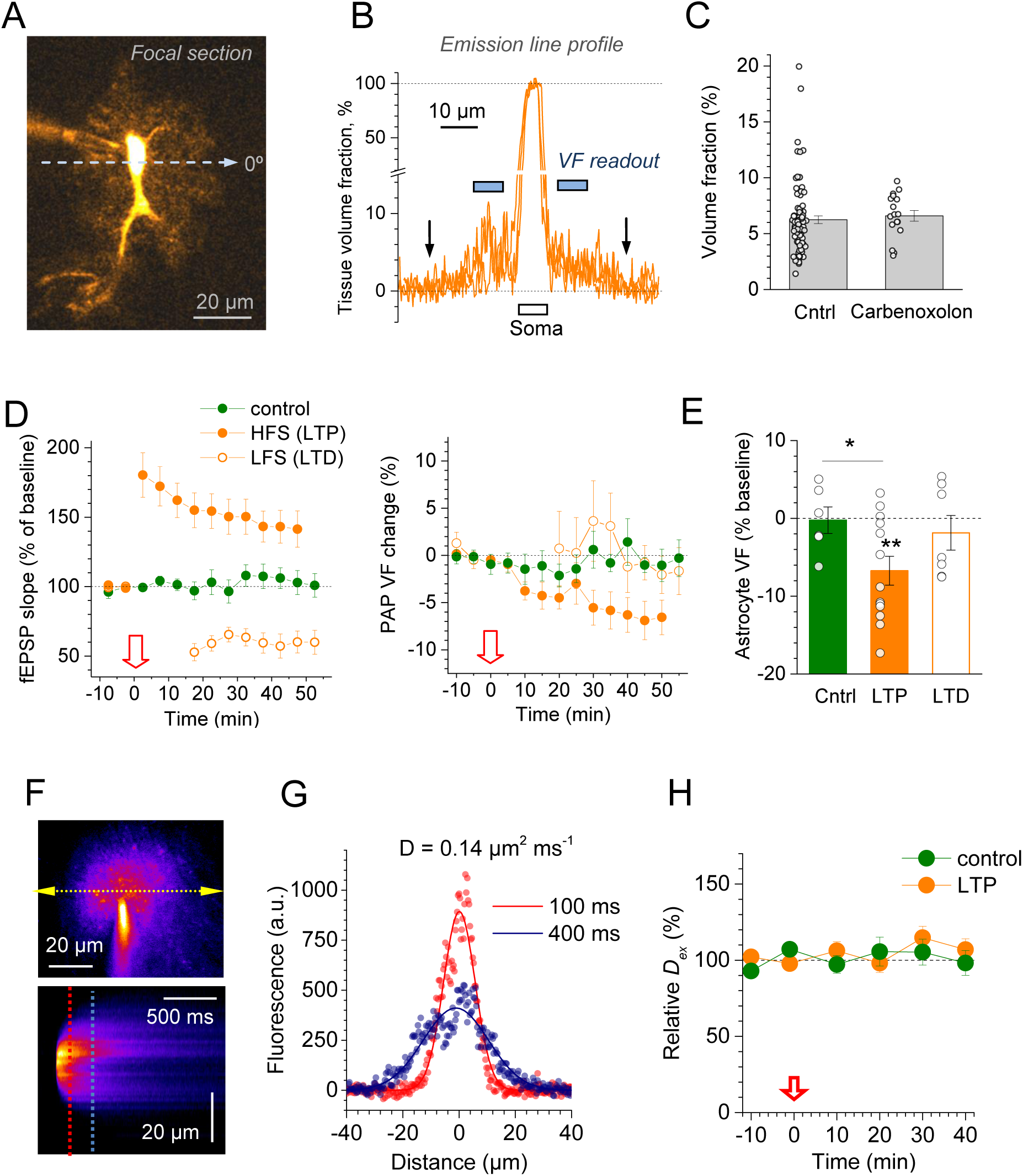
PAP volume fraction and extracellular diffusivity monitored using two-photon excitation imaging. (A) CA1 astrocyte, imaged in a single 2PE section (λ_x_^2p^ = 800 nm; Alexa Fluor 594 hydrazide; 50 µM gap-junction blocker carbenoxolone); dashed line, sampling of the emission intensity profile (an example crossing the soma; angle 0°; false colour scale). (B) Example of three astrocyte fluorescence profiles (measured using the original grey-level image, background-corrected): along the sampling line at 0° (shown in a), at 45°, and 135°. Data normalised to the emission intensity within the soma; arrows, detectable edges of the astrocyte profile; blue segments, typical ROI position for averaging VF readout during LTP induction (as shown in Figure 1B): ROI was selected to avoid thick primary processes near the soma or the uneven edges of the astroglial arbour. (C) Average VF values for astrocyte arbours in control conditions (mean ± SEM: 6.2 ± 0.35%, *n =* 83) were not statistically different from those obtained in the presence of the gap junction blocker carbenoxolone (6.6 ± 0.48%, *n =* 17, p = 0.56, unpaired Student’s t-test). (D) Time course of fEPSP slope (left) and PAP VF (right) relative to baseline (%, mean ± SEM); VF measured using astroglial EGFP expressed under a human GFAP promoter (hGFAP-EGFP) (Nolte et al., 2001); in control conditions (green dots, n = 6), during HFS LTP induction (solid orange circles; red arrow, onset; n = 13), and during the induction of LTD (low-frequency induction protocol, 1800 stimuli at 2 Hz; open orange); red arrow, onset of LTP/LTD induction. (E) Summary of PAP VF changes (%, mean ± SEM) in experiments shown in d, 40-50 min after plasticity induction onset: LTP (−7 ± 2%, n = 13, ** p < 0.004, t-test; p = 0.0081, Wilcolxon Signed Rank Test, W = 9, Z = −2.516), control (+ 0.23 ± 1.7 %, n = 6, difference with LTP at *p = 0.021), LTD (−1.9 ± 2.2 %, n = 7, p = 0.43). Note that the dynamics of PAP VF readout in EGFP-expressing cells, unlike Alexa-loaded cells, is likely to underestimate and fall behind real VF changes because diffusion equilibration of EGFP is much slower slow than that of Alexa (MW are 33 kDa and 760 Da, respectively). (F) Measuring local extracellular diffusivity. *Top:* A snapshot of a pressurised pipette filled with Alexa Fluor 594 and placed in CA1 *s. radiatum* area near recorded astroglia; arrows, linescan position near the pipette tip. *Bottom*: Linescan recording of a brief (2-6 ms) pulse ejection of the dye; dashed lines, sampling the extracellular fluorescence intensity profiles (readout of dye concentration) at 100 ms and 400 ms post-pulse (Zheng et al., 2008). (G) Fluorescence intensity profiles (dye concentration readout, shown in F) approximated with a Gaussian function, as indicated. Effective extracellular diffusion coefficient (indicated) is obtained from the time evolution of such profiles, as described earlier (Zheng et al., 2008). (H) Time course of extracellular diffusivity in experiments with and without LTP induction (arrow), as indicated; average extracellular diffusivity 40 min after LTP induction: 107.0 ± 7.1% of baseline (*n =* 8); 40 min time point in control (no-LTP) experiments was 98.2 ± 8.2% of baseline (*n =* 6).

**Figure S2 (related to Figure 2).**
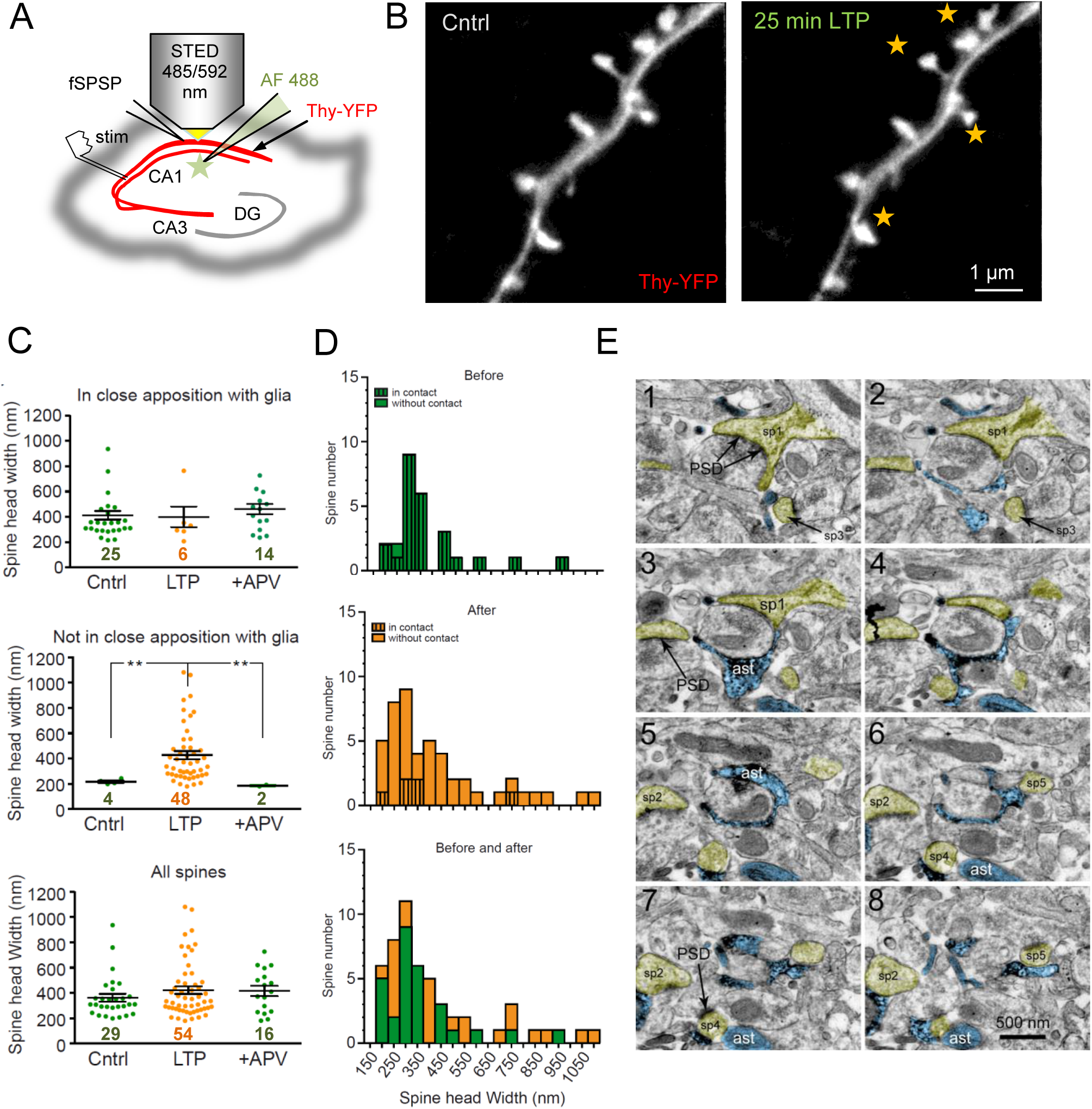
STED imaging and 3D EM of LTP-associated changes in perisynaptic astroglial environment. (A) Experiment diagram: CA3-CA1 LTP is induced in organotypic hippocampal slices, with Thy-YFP-labelled CA1 pyramidal cells (red channel) and whole-cell recorded astroglia (Alexa Fluor 488, green channel) for STED imaging. (B) Characteristic STED images of CA1 pyramidal cell dendritic spines (optimised Thy1- YFP channel), before and ∼25 min after LTP induction, as indicated (Methods). Stars indicate visible alterations in spine head geometry after LTP induction. (C) Effects of LTP induction on the occurrence of close astroglia-spine apposition and on the dendritic spine head sizes, monitored with STED microscopy. Digits, numbers of identified spine heads under pseudo-random inspection of the area of interest; bars, mean ± SEM; LTP, data ∼25 min after LTP induction; +APV, same protocol but in the presence of 50 µM APV; ** p < 0.01 (Kruskall-Wallis ANOVA, sample sizes *n* shown). (D) Frequency histograms of spine head sizes before and 20-25 min after LTP induction (data set as in C), also with the separation between spines which are closely approached (striped columns) and not approached (plain columns) by PAPs, as shown. (E) 3D electron microscopy: Example of serial sections containing profiles of the astroglia recorded and labelled in an acute slice; eight (1-8) 60 nm thick adjacent serial sections are shown, depicting dendritic spines (yellow shade; sp1-sp5) and astroglia (blue, ast; filled with electron dense precipitate from DAB conversion of the recorded astrocyte filled with biocytine); scale bar, 500 nm.

**Figure S3.**
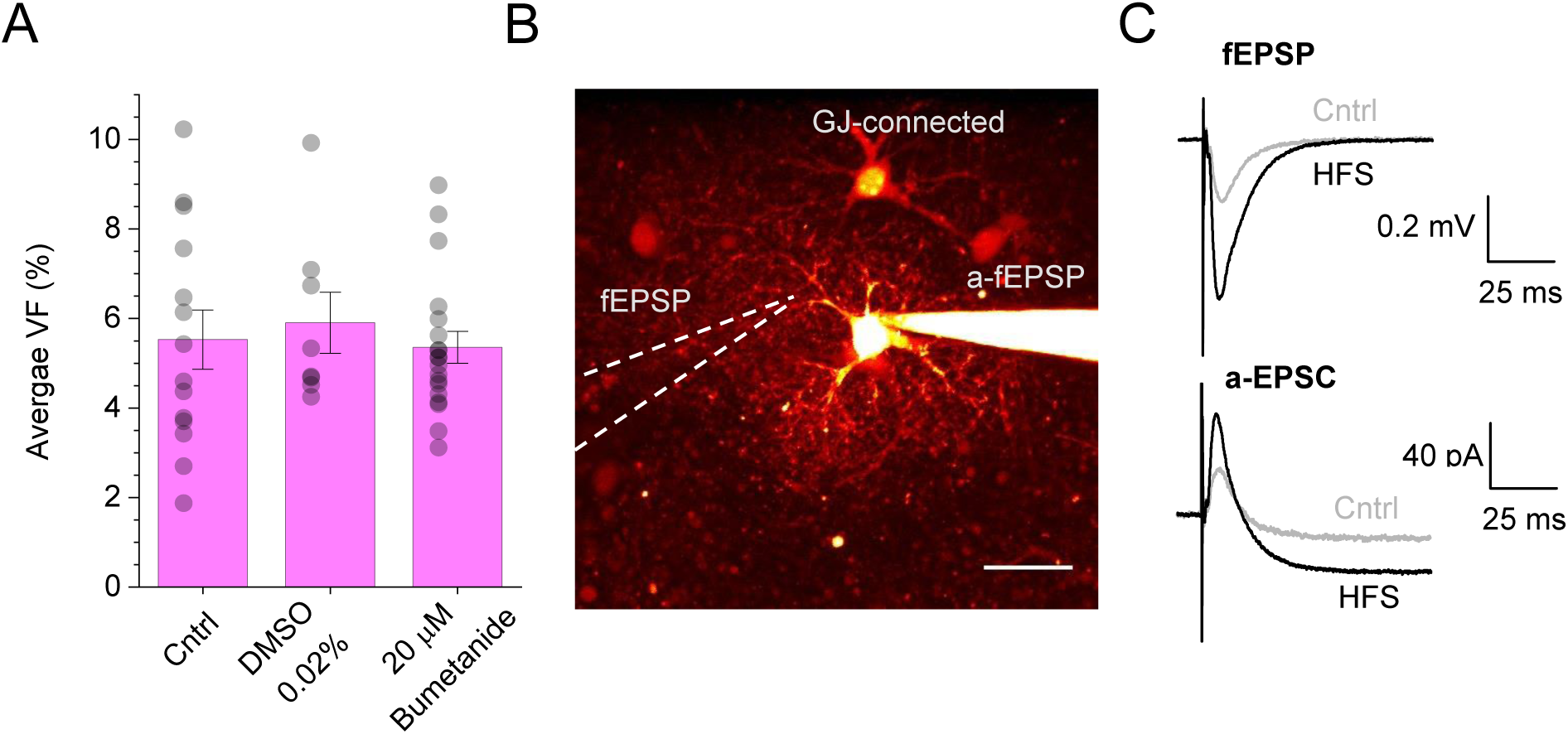
Control tests for astroglial VF in bumetanide and LTP induction with S3 peptide. (A) Astroglial PAP VF (%, mean ± SEM) measured in control conditions (Cntrl, 5.52 ± 0.66; n = 14), with the vehicle 0.02% DMSO (5.90 ± 0.68; n = 8), and with 20 µM bumetanide inside the cell (5.35 ± 0.36; n = 19). (B) Experimental arrangement in LTP experiments (including S3 peptide dialysis), in acute slices, area CA1 *s. radiatum*; Alexa Fluor 594 channel, λ_x_^2P^ = 800 nm. Locations of the whole-cell patch pipette (recording astroglial a-fEPSPs or a-EPSCs), extracellular electrode (recording fEPSPs), and gap junction connected astroglia (GJ-connected) are indicated. (C) Characteristic recordings before and ∼20 min after high-frequency stimulation (HFS) induced LTP using an extracellular electrode (fEPSP) and an astrocyte pipette (a-EPSC) (Henneberger and Rusakov, 2012), in the same peptide S3 experiment.

**Figure S4 (related to Figure 4).**
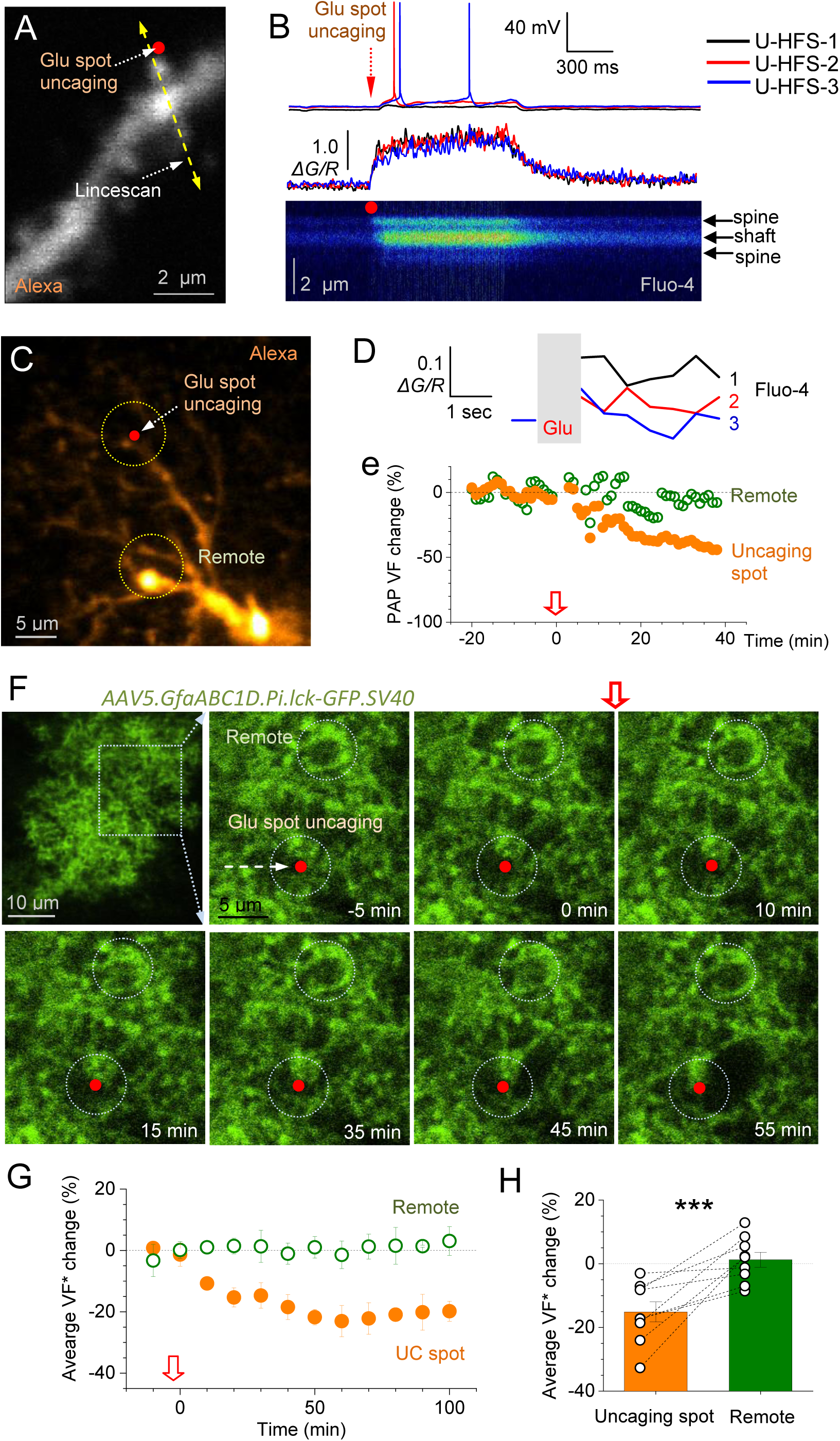
LTP induction by glutamate spot-uncaging induces postsynaptic depolarization and Ca^2+^ entry while triggering local PAP VF reduction. (A) Dendritic fragment, CA1 pyramidal cell (Alexa channel); red dot, MNI-glutamate uncaging spot; yellow arrow, linescan position; whole-cell current-clamp mode (V_m_ = - 60…65 mV in baseline conditions); uncaging at λ_u_^2P^ = 720 nm; Alexa imaging at λ_x_^2P^ = 840 nm. (B) Somatic voltage recording (upper traces) and postsynaptic Ca^2+^-sensitive linescan recording (bottom traces, Δ*G/R;* linescan, Fluo-4 channel) during LTP induction (100 x 1 ms laser pulses at 100Hz, three series 60 s apart), three traces superimposed (colour-coded; U-HFS-1/2/3); intermittent spikes can be seen in some traces. (C) Astrocyte fragment (averaged 9-section z-stack) depicting the uncaging spot (red dot) and two ROIs for PAP VF measurement; Alexa Fluor 594 channel. (D) Time course of Ca^2+^-sensitive fluorescence (averaged within ROI1), before and immediately after 2P spot-uncaging (grey segment), for series of pulses 1, 2, and 3, as shown in (B). (E) Relative change in PAP VF (%) in ROI1 and ROI2 shown in (C) during LTP induction (red arrow, onset). (F) One-cell example of PAP changes during LTP induction with glutamate spot-uncaging; the astrocyte-membrane bound GFP (transduction with AAV5.GfaABC1D.Pi.lck-GFP.SV40) provides readout of PAP (membrane) presence within ROI (PAP VF*, integrated brightness within ROI). Top left panel: astrocyte view; dotted rectangle, monitored area (10 µm deep z-stack average). Other panels: time-lapse images of the monitored area (single 2PE focal section, time stamps shown), during LTP induction (red arrow, onset); circles, ROIs for measuring PAP VF* near the uncaging spot and within a control / remote area (Remote), as indicated; acute hippocampal slice, *stratum radiatum*. (G) Time course of PAP VF* change relative to baseline (%, mean ± SEM, n = 8 slices) in the experiments illustrated in (F). (H) Summary of experiments shown in (F-G), for the PAP VF* changes over 25-35 min after LTP induction; bar graphs, mean ± SEM (n = 8); connected dots, individual experiments; ***, p < 0.001 (paired *t*-test).

**Figure S5 (related to Figure 5).**
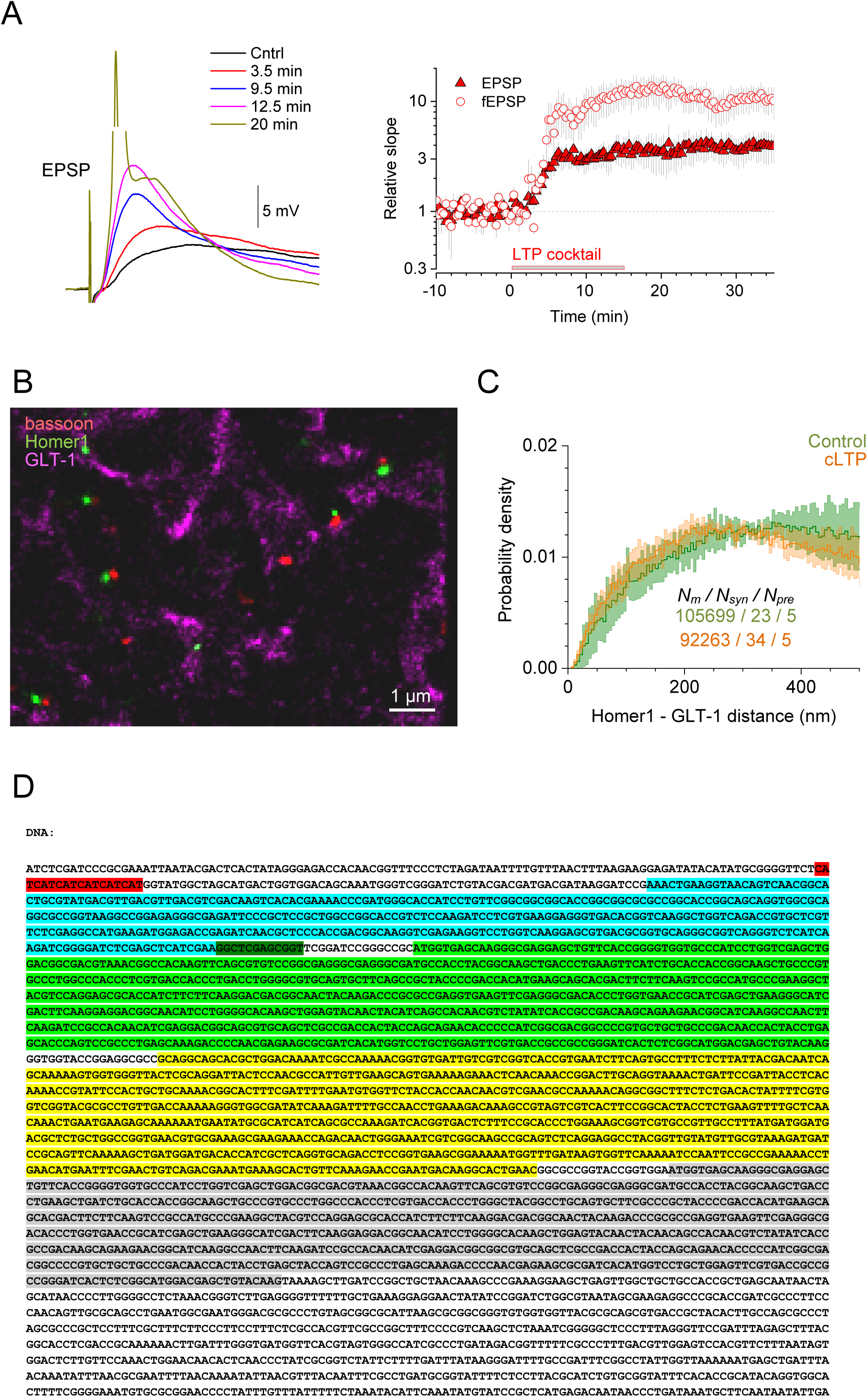
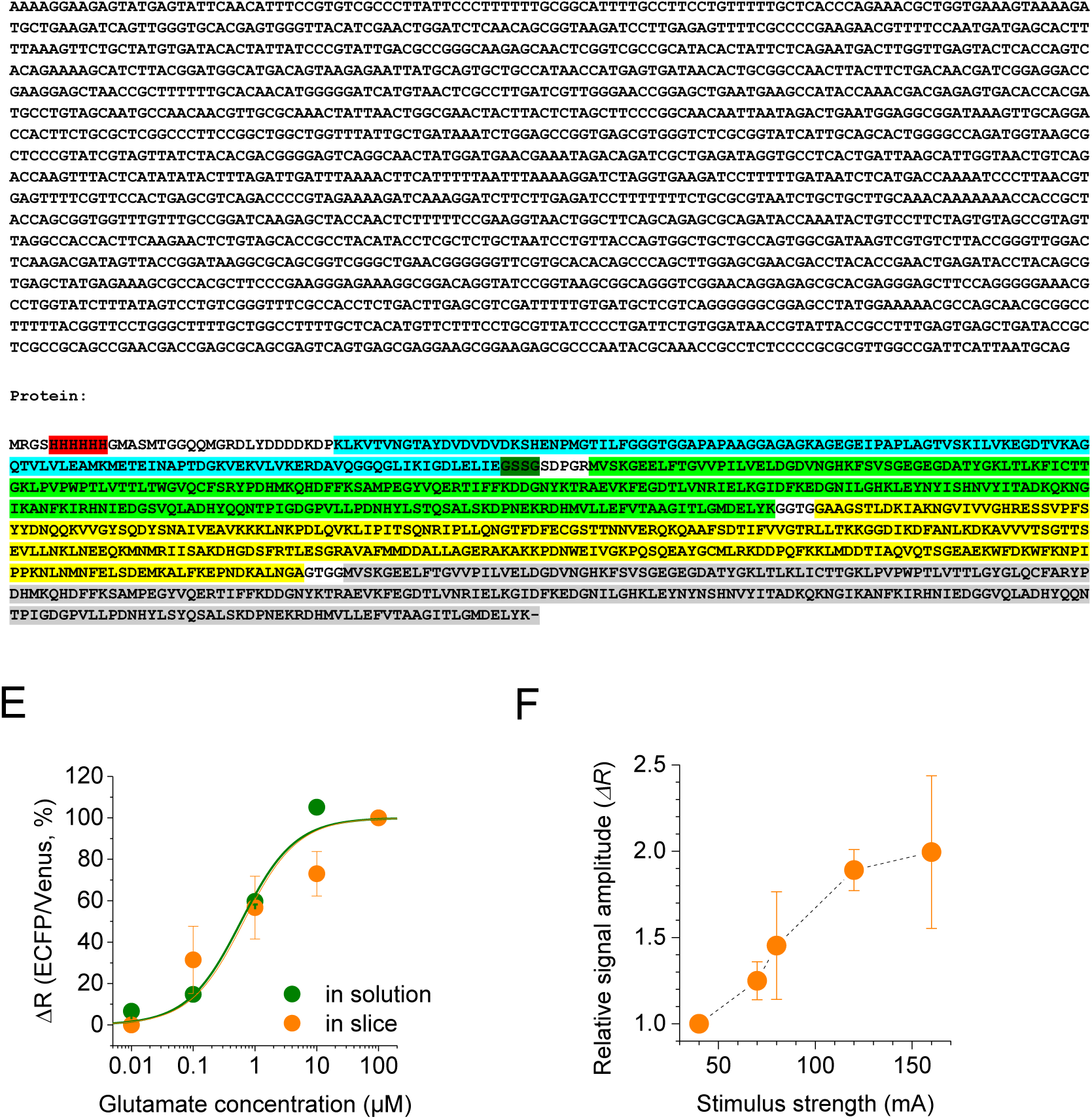
LTP-associated perisynaptic withdrawal of glutamate uptake system documented with super-resolution dSTORM and extracellular glutamate sensor bFLIPE600n. (A) Induction of chemical long-term potentiation (cLTP) in acute hippocampal slices. Traces, characteristic EPSPs (whole-cell current clamp) recorded in CA1 pyramidal cells in response to electrical stimulation of Schaffer collaterals, in control conditions (Cntrl), and at different time points after the onset of ‘LTP cocktail’ application (50 µM forskolin and 100 nM rolipram), as indicated. Graph, average fEPSP slope and whole-cell EPSPs relative to baseline (mean ± SEM, n = 8 slices) during cLTP induction. (B) 3D three-color dSTORM: a snapshot of molecular patterns for presynaptic bassoon (CF-568, red), postsynaptic Homer 1 (Atto-488, green), and glutamate transporter GLT-1 (Alexa-647, magenta), in the *stratum radiatum* (acute slices, 1-month-old rats, control conditions); images show 2D projections of 3D dSTORM molecular maps (∼4 µm deep stacks) in 30-µm acute slices; label brightness reflects molecular density, high-resolution raw data include single-molecule 3D co-ordinates. (C) Average distribution (probability density, mean ± SEM) of the nearest-neighbour distances (<500 nm) between GLT-1 and postsynaptic Homer1 molecules, in control and potentiated tissue (∼30 min after ‘chemical’ LTP induction; Methods), as indicated; summary for *N_m_* inter-molecular distances at *N_syn_* synapses from *N_pre_* individual slices; SEM reflects variance among *N_pre_* = 5 slices. (D) Glutamate sensor bFLIPE600n: DNA and protein sequences. Functionally relevant parts are colour coded, as follows: His-tag, Biotin-tag, ECFP, flexible linker (GSSG) after biotin tag, GltI, Venus. (E) Measuring glutamate sensitivity of bFLIPE600n immobilised in acute hippocampal slices. Glutamate levels were estimated by calculating the fluorescence intensity ratio *R* = ECFP/Venus and its changes (*ΔR*). Titration of bFLIPE600n was performed in free solution and acute slices (*ΔR* versus nominally zero glutamate; free solution: Kd = 596 nM, n = 3; acute slices Kd = 659 nM, n = 5). *In situ* measurements were performed in the presence of 10 µM NBQX, 50 µM D-APV, 1 µM TFB-TBOA, 1 µM TTX, 50 µM LY341495. Calibration curves, Hill equation approximation (Hill coefficient set to one). (F) The input-output evaluation bFLIPE600n sensitivity: glutamate transient amplitude versus stimulation intensity (mean ± SEM, n = 4) in CA1 *stratum radiatum*; *ΔR* signal is normalized to the baseline response at 40 µA stimulus.

**Figure S6 (related to Figure 6).**
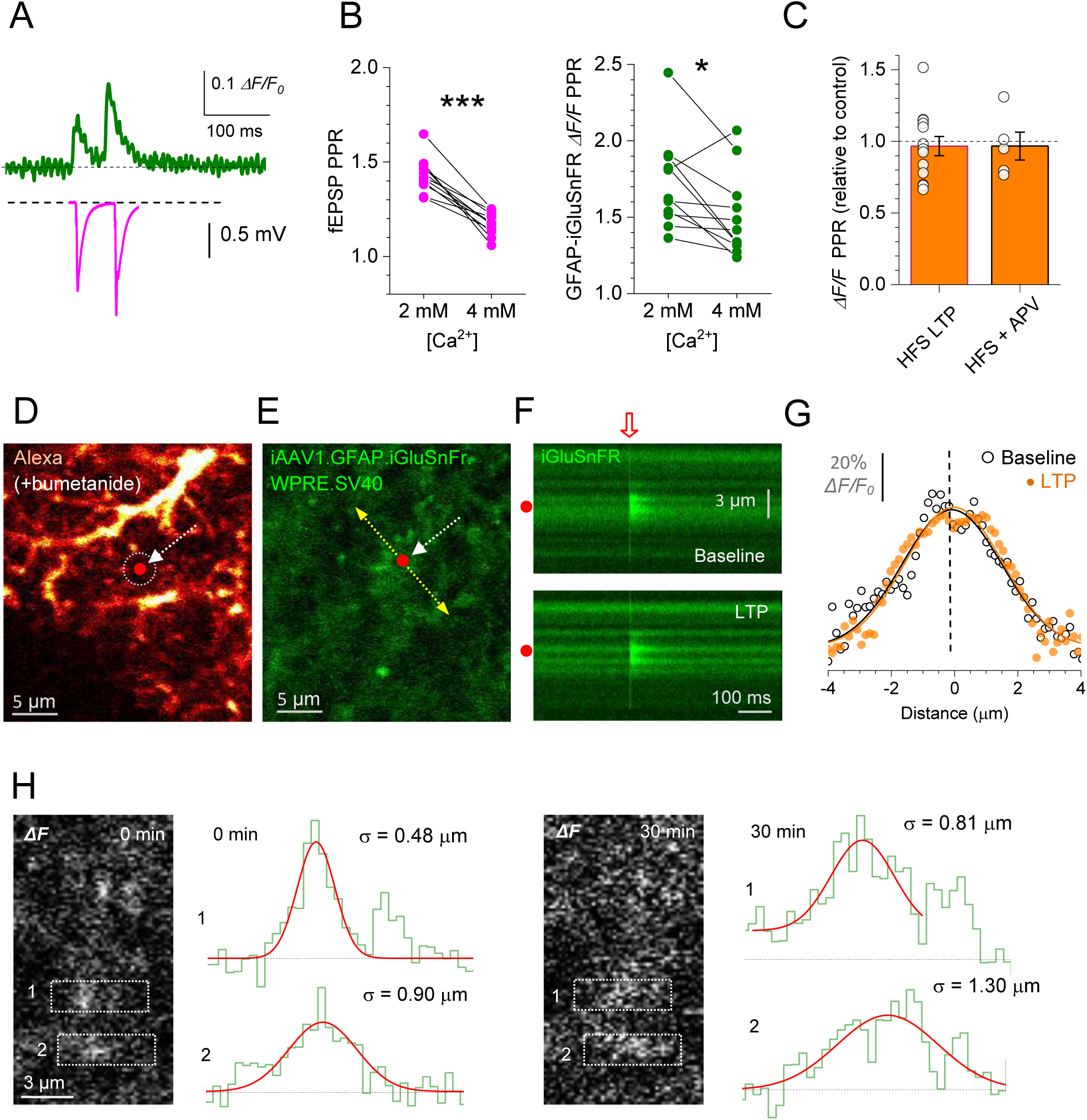
Monitoring extrasynaptic glutamate escape with the optical glutamate sensor iGluSnFR. (A) iGluSnFR fluorescence intensity traces (*ΔF/F_0_*) in response to paired-pulse stimuli (50 ms apart; top trace, average of 100 trials) faithfully reflect fEPSPs recorded in parallel (bottom trace). (B) Paired-pulse experiments (as in A) showing that increasing extracellular [Ca^2+^] reduces paired-pulse ratio (PPR) for both fEPSPs (left; p < 0.001) and optical iGluSnFR responses (right; p = 0.019; n = 11; paired *t*-tests). (C) HFS-induced LTP (n = 13), or HFS in the presence of 50 µM APV (n = 5), have no effect on the paired-pulse ratio (PPR) of iGluSnFR *ΔF/F* responses. (D) Fragment of the astrocyte (whole-cell loaded with 100 µM Alexa Fluor 594 and 20 µM bumetanide; λ_x_^2P^ = 910 nm; single optical section) illustrating the LTP induction protocol with spot-uncaging of glutamate (red dot; λ_u_^2P^ = 720 nm); dotted circle, ROI for monitoring local PAP VF during LTP induction. See Figure 6D for statistical summary. (E) Fragment in d shown in the iGluSnFR channel (λ_x_^2P^ = 910 nm); arrow, linescan position; other notations as in (D). (F) Examples of linescan traces (positioned as in E) recorded before and ∼20 min after the LTP induction spot-uncaging protocol (baseline and LTP, respectively). (G) Spatial glutamate-sensitive iGluSnFR fluorescence profiles (dots, individual pixel values) evoked by a 1 ms glutamate uncaging pulse, before (Baseline) and 20-25 min after LTP induction, as indicated, in the experiment illustrated in (D-F); zero abscissa, the uncaging spot position (red dot in D-E); black and orange solid lines, best-fit Gaussian approximation. See Figure 6d for statistical summary. (H) Example illustrating evaluation of the axonal glutamate signal spread in the experiment shown in Figure 6E-F, just before (0 min) and 30 min after LTP induction, as indicated. Image panels, grey-level images of the *ΔF = F - F_0_* signal (AAV9.hSynap. iGluSnFr.WPRE.SV40 fluorescence landscape obtained by subtracting the baseline landscape from that during five afferent stimuli at 20Hz); dotted rectangles, sampling (horizontal) segments to acquire brightness profiles associated with tentative axonal boutons 1 and 2, as indicated; plots, the corresponding brightness profiles (staircase, green) with the best-fit Gaussian (red); σ, Gaussian standard deviation (FWHM=2.35σ). Note that in the Bouton 1 Gaussian fitting ignores the neighbouring bouton (a second fluorescence peak to the right).

**Figure S7 (related to Figure 7).**
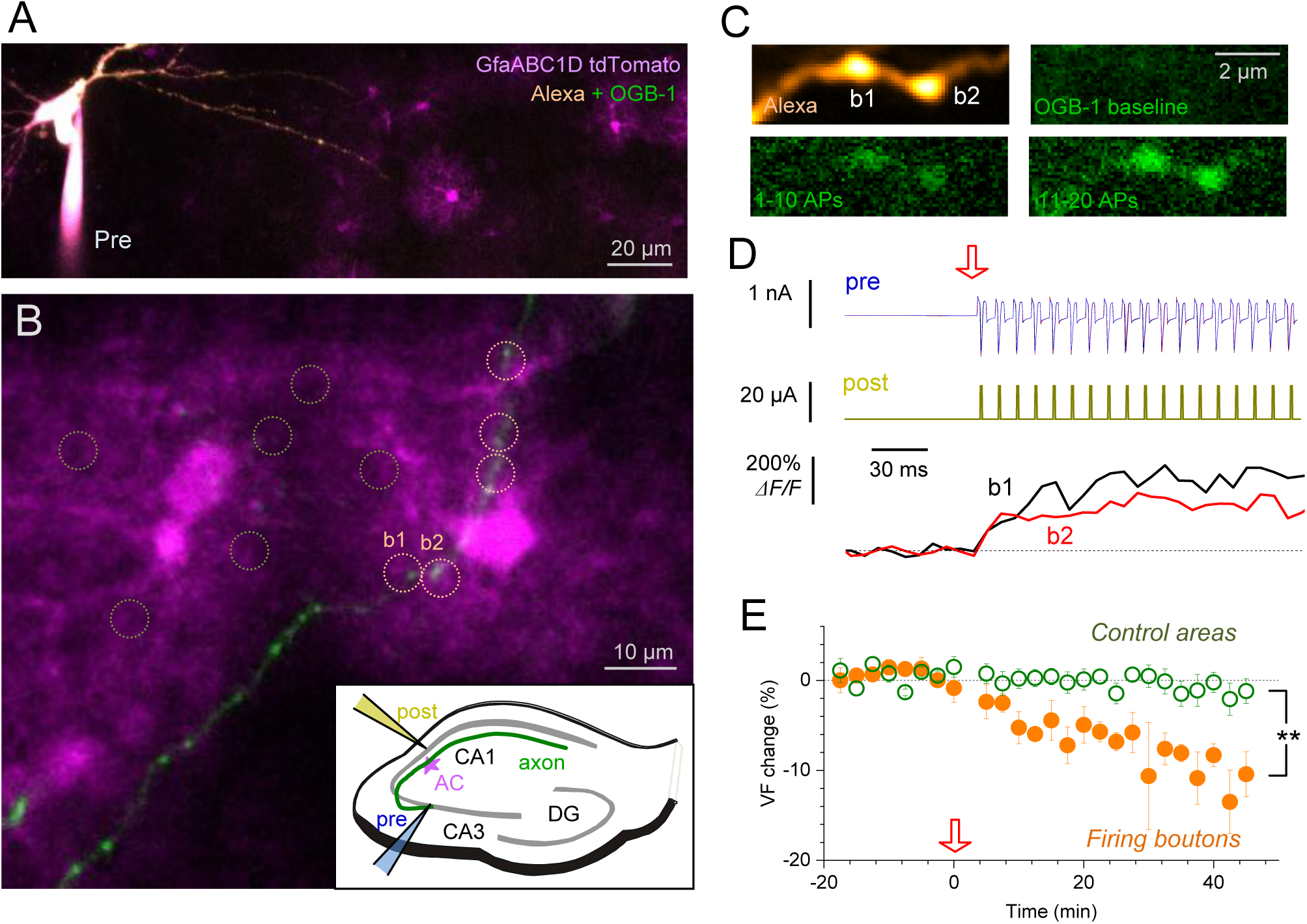
LTP pairing protocol for a single CA3 pyramidal cell axon leads to astroglial withdrawal near active axonal boutons. (A) CA3 pyramidal cell shown held in whole-cell (dialysed with 50 µM Alexa 594 and 200 µM OGB-1), with the axon proximal part seen traced into the field of astrocytes labelled with tdTomato (magenta); a 67 µm deep z-stack projection image (two-laser 2PE at λ_x_^2p^ = 800 nm and λ_x_^2p^ = 910 nm). (B) A distal fragment of the same axon (green) which crosses local CA1 astroglia (magenta); orange dotted circles, ROIs for PAP VF monitoring near five firing presynaptic boutons (see below for Ca^2+^ recordings in boutons b1 and b2); green dotted circles, examples of control astroglial areas devoid of the presynaptic axon; a 67 µm deep z-stack projection image for two-laser 2PE as in (A). Inset, experiment diagram of LTP induction protocol by pairing, depicting presynaptic cell held in whole-cell (as in A), a postsynaptic stimulating electrode placed in *stratum pyramidale* (yellow; to depolarize / fire postsynaptic cells), a single traced axon (green) crossing an astrocyte (AC, magenta). (C) Examples of two presynaptic axonal boutons (b1 and b2 in b), shown in the Alexa channel (red) and in the Ca^2+^ sensitive OGB-1 channel (green) in baseline conditions and following the initial burst of presynaptic action potentials Nos 1-10, and 11-20 (at 100 Hz), as indicated. (D) Examples of presynaptic whole-cell electrode recordings (blue, individual evoked spikes shown), postsynaptic extracellular electrode pulse control (dark yellow), and OGB-1 fluorescence in boutons b1 and b2 (shown in B-C), in the initial phase of the LTP induction protocol (3 x 1 s trains @ 100 Hz; red arrow, onset). (E) Time course of PAP VF changes (%, mean ± SEM) near firing axonal boutons (orange dots, the corresponding ROIs shown by orange circles in b; n = 5) and in control areas (ROI examples shown by green circles in b; n = 10) during LTP induction (red arrow, onset); **, p < 0.01 (two-sample *t*-test for 25-45 min interval post-induction); the data were routinely adjusted to account for the (small) experiment-wide tdTomato photobleaching, which was fitted by *P*(*t*) = 0.9723 + 0.00867 *e*^−*t/τ*^ with *τ* = 10.9 min.

**Figure S8 (related to Figure 8).**
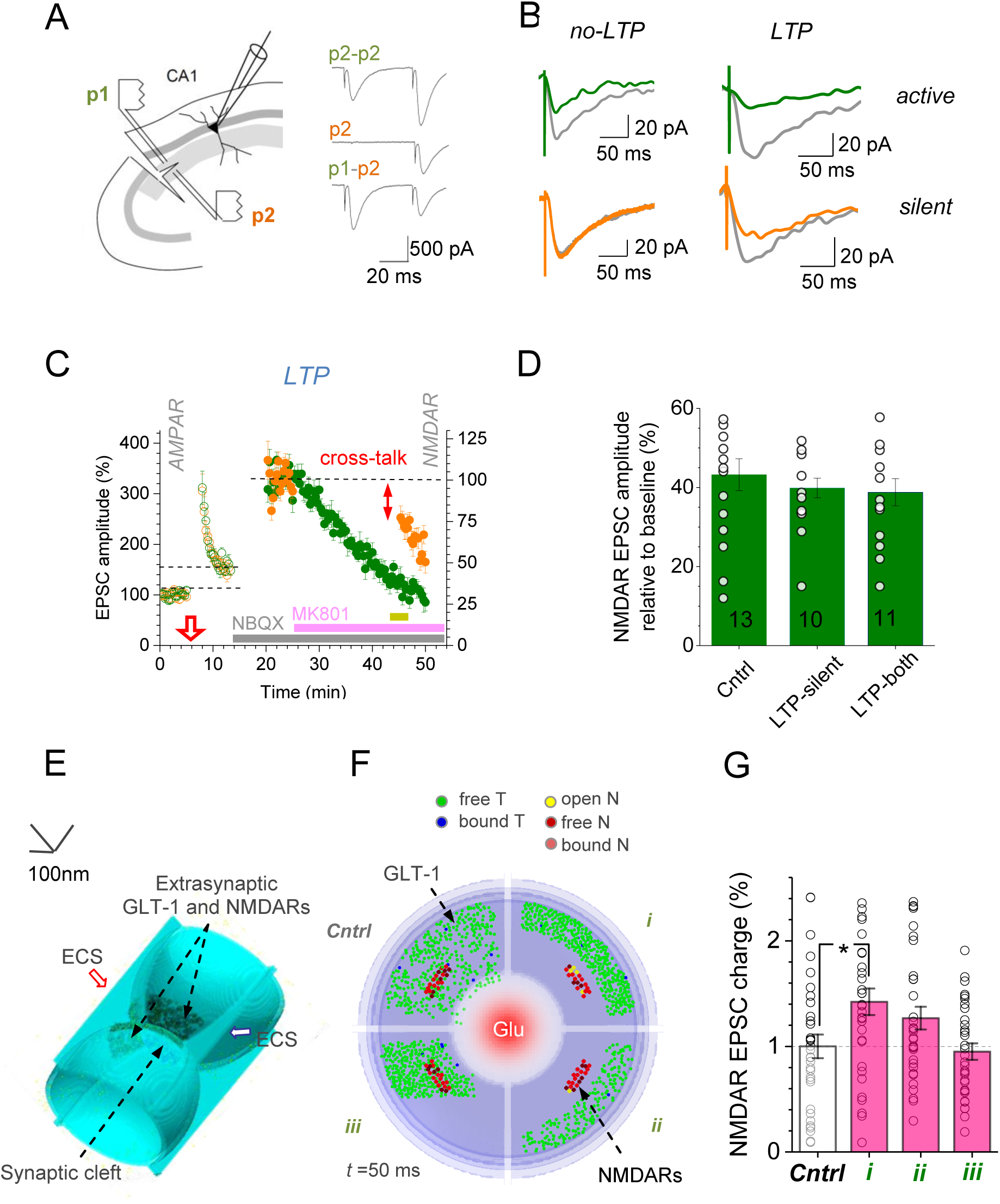
Electrophysiological probing of NMDAR-mediated inter-synaptic cross-talk and a biophysical plausibility test. (A) Experiment diagram (left), a previously established test (Scimemi et al., 2004) for presynaptic independence of two Schaffer collateral pathways (p1 and p2) converging onto a CA1 pyramidal cell. Traces, EPSCs show paired-pulse facilitation during p2-p2 stimulation but not across the pathways (p1-p2, modified from (Scimemi et al., 2004)). In the recorded sample (*n* = 54 slices), paired-pulse facilitation with a 50 ms interval was 75.4 ± 6.1 % (mean ± SEM; p < 0.001) in the same pathway and only 16.5 ± 2.9% across (difference at p < 0.001). (B) Characteristic one-cell examples (summaries shown in Figure 8A-B) of NMDAR EPSCs recorded in both pathways, in baseline conditions (grey) and after resuming stimulation of the silent (green) pathway; in baseline conditions (*no-LTP*) and with LTP induced in the active (orange) pathway during AMPA EPSC recording (*LTP*). (C) Summary of two-pathway experiments with LTP induced simultaneously in both (rather than one) pathways. Notations are as in Figure 8A; yellow segment, period over which the degree of NMDAR EPSC suppression by MK801, prior to the resumption of silent pathway stimulation, was measured. (D) The reduction of the NMDAR EPSC amplitude (mean ± SEM, sample size n shown) following application of MK801, prior to the resumption of test pathway stimulation (averaging time interval depicted by yellow segment in c), in experiments with no LTP, with LTP induced in the silent pathway (LTP-silent), and with LTP induced in both pathways (LTP-both), as indicated. See Figure 8A for the full time course data. No difference in the MK801-induced decay among the three conditions indicate no difference in release probabilities in the bulk of stimulated synapses, with or without LTP induction. (E) A 3D Monte-Carlo model depicting microenvironment of the CA3-CA1 synapses. Truncated hemispheres represent surfaces of the presynaptic bouton and the postsynaptic dendritic spine separated by the synaptic cleft and surrounded by extracellular space gaps (ECS). Extrasynaptic clusters of astroglial transporters (GLT-1, on the PAP surface) and NMDA receptors (on the surface of dendrites facing PAPs) are shown (Methods; (Zheng et al., 2008)). (F) Front view (cross-section) of the model shown in e. Four scenarios of astroglial GLT-1 relocation (green dot scatter) during LTP are shown in four different quadrants: *Cntrl* (random scatter), *i* (even withdrawal/shrinkage, same transporter numbers)*, ii* (even withdrawal, same transporter density, ∼50% drop in numbers), *and iii* (withdrawal to one side, same transporter numbers). Images illustrate a snapshot of NMDAR activation 50 ms following release of 3000 glutamate molecules from the centre (red shade); GLT-1 transporter (T) and NMDAR (N) states are color-coded, as indicated (Video 2). (G) Summary of Monte Carlo experiments (n = 32 runs) shown in (E-F). Average NMDAR-mediated charge transfer relative to Control (*Cntrl*), for the three types of change: *i* (mean ± SEM; 1.42 ± 0.13, p < 0.015 compared to control), *ii* (1.26 ± 0.11), and *iii* (0.95 ± 0.08); dots, data points at individual runs; note high variability due to a few NMDARs being activated stochastically per release. The data indicate that scenario *i* is most likely to correspond to the larger increase in extrasynaptic NMDAR activation after LTP induction. Note that the higher transporter density in scenario *i*, if compared to scenario *ii,* prolongs local dwell time of glutamate molecules, due to unbinding form transporters.

## Supplementary References

Henneberger, C., and Rusakov, D.A. (2012). Monitoring local synaptic activity with astrocytic patch pipettes. Nature Protocols 7, 2171–2179.

Nolte, C., Matyash, M., Pivneva, T., Schipke, C.G., Ohlemeyer, C., Hanisch, U.K., Kirchhoff, F., and Kettenmann, H. (2001). GFAP promoter-controlled EGFP-expressing transgenic mice: a tool to visualize astrocytes and astrogliosis in living brain tissue. Glia 33, 72–86.

Scimemi, A., Fine, A., Kullmann, D.M., and Rusakov, D.A. (2004). NR2B-containing receptors mediate cross talk among hippocampal synapses. J. Neurosci. 24, 4767–4777.

Zheng, K., Scimemi, A., and Rusakov, D.A. (2008). Receptor actions of synaptically released glutamate: the role of transporters on the scale from nanometers to microns. Biophys J 95, 4584–4596.

## REFERENCES

Adamsky, A., Kol, A., Kreisel, T., Doron, A., Ozeri-Engelhard, N., Melcer, T., Refaeli, R., Horn, H., Regev, L., Groysman, M., et al. (2018). Astrocytic Activation Generates De Novo Neuronal Potentiation and Memory Enhancement. Cell 174, 59–71 e14.

Aizawa, H., Wakatsuki, S., Ishii, A., Moriyama, K., Sasaki, Y., Ohashi, K., Sekine-Aizawa, Y., Sehara-Fujisawa, A., Mizuno, K., Goshima, Y., and Yahara, I. (2001). Phosphorylation of cofilin by LIM-kinase is necessary for semaphorin 3A-induced growth cone collapse. Nat Neurosci 4, 367–373.

Anders, S., Minge, D., Griemsmann, S., Herde, M.K., Steinhauser, C., and Henneberger, C. (2014). Spatial properties of astrocyte gap junction coupling in the rat hippocampus. Philos Trans R Soc Lond B Biol Sci 369, 20130600.

Arnth-Jensen, N., Jabaudon, D., and Scanziani, M. (2002). Cooperation between independent hippocampal synapses is controlled by glutamate uptake. Nature Neurosci 5, 325–331.

Bernardinelli, Y., Randall, J., Janett, E., Nikonenko, I., Konig, S., Jones, E.V., Flores, C.E., Murai, K.K., Bochet, C.G., Holtmaat, A., and Muller, D. (2014). Activity-dependent structural plasticity of perisynaptic astrocytic domains promotes excitatory synapse stability. Curr Biol 24, 1679–1688.

Bliss, T.V.P., Douglas, R.M., Errington, M.L., and Lynch, M.A. (1986). Correlation between Long-Term Potentiation and Release of Endogenous Amino-Acids from Dentate Gyrus of Anesthetized Rats. J Physiol 377, 391–408.

Bravo-Cordero, J.J., Magalhaes, M.A., Eddy, R.J., Hodgson, L., and Condeelis, J. (2013). Functions of cofilin in cell locomotion and invasion. Nat Rev Mol Cell Biol 14, 405–415.

Bushong, E.A., Martone, M.E., Jones, Y.Z., and Ellisman, M.H. (2002). Protoplasmic astrocytes in CA1 stratum radiatum occupy separate anatomical domains. J Neurosci 22, 183–192.

Carter, A.G., and Regehr, W.G. (2000). Prolonged synaptic currents and glutamate spillover at the parallel fiber to stellate cell synapse. J Neurosci 20, 4423–4434.

Chalifoux, J.R., and Carter, A.G. (2011). Glutamate Spillover Promotes the Generation of NMDA Spikes. J Neurosci 31, 16435–16446.

Coddington, L.T., Rudolph, S., Vande Lune, P., Overstreet-Wadiche, L., and Wadiche, J.I. (2013). Spillover-Mediated Feedforward Inhibition Functionally Segregates Interneuron Activity. Neuron 78, 1050–1062.

Danbolt, N.C. (2001). Glutamate uptake. Progr Neurobiol 65, 1–105.

Diamond, J.S., Bergles, D.E., and Jahr, C.E. (1998). Glutamate release monitored with astrocyte transporter currents during LTP. Neuron 21, 425–433.

Dittman, J.S., and Regehr, W.G. (1997). Mechanism and kinetics of heterosynaptic depression at a cerebellar synapse. J Neurosci 17, 9048–9059.

Dityatev, A., and Rusakov, D.A. (2011). Molecular signals of plasticity at the tetrapartite synapse. Curr Opin Neurobiol 21, 353–359.

Dityatev, A., and Schachner, M. (2003). Extracellular matrix molecules and synaptic plasticity. Nat Rev Neurosci 4, 456–468.

Endesfelder, U., and Heilemann, M. (2015). Direct stochastic optical reconstruction microscopy (dSTORM). Methods Mol Biol 1251, 263–276.

Epsztein, J., Lee, A.K., Chorev, E., and Brecht, M. (2010). Impact of spikelets on hippocampal CA1 pyramidal cell activity during spatial exploration. Science 327, 474–477.

Errington, M.L., Galley, P.T., and Bliss, T.V.P. (2003). Long-term potentiation in the dentate gyrus of the anaesthetized rat is accompanied by an increase in extracellular glutamate: real-time measurements using a novel dialysis electrode. Phil Trans Roy Soc ser B 358, 675–687.

Ethell, I.M., and Pasquale, E.B. (2005). Molecular mechanisms of dendritic spine development and remodeling. Prog Neurobiol 75, 161–205.

Filosa, A., Paixao, S., Honsek, S.D., Carmona, M.A., Becker, L., Feddersen, B., Gaitanos, L., Rudhard, Y., Schoepfer, R., Klopstock, T., et al. (2009). Neuron-glia communication via EphA4/ephrin-A3 modulates LTP through glial glutamate transport. Nat Neurosci 12, 1285–1292.

Florence, C.M., Baillie, L.D., and Mulligan, S.J. (2012). Dynamic Volume Changes in Astrocytes Are an Intrinsic Phenomenon Mediated by Bicarbonate Ion Flux. PLoS One 7.

Frick, A., Magee, J., and Johnston, D. (2004). LTP is accompanied by an enhanced local excitability of pyramidal neuron dendrites. Nat Neurosci 7, 126–135.

Gambino, F., Pages, S., Kehayas, V., Baptista, D., Tatti, R., Carleton, A., and Holtmaat, A. (2014). Sensory-evoked LTP driven by dendritic plateau potentials in vivo. Nature 515, 116–119.

Garzon-Muvdi, T., Schiapparelli, P., ap Rhys, C., Guerrero-Cazares, H., Smith, C., Kim, D.H., Kone, L., Farber, H., Lee, D.Y., An, S.S., et al. (2012). Regulation of brain tumor dispersal by NKCC1 through a novel role in focal adhesion regulation. PLoS Biol 10, e1001320.

Grosche, J., Matyash, V., Moller, T., Verkhratsky, A., Reichenbach, A., and Kettenmann, H. (1999). Microdomains for neuron-glia interaction: parallel fiber signaling to Bergmann glial cells. Nat Neurosci 2, 139–143.

Gundelfinger, E.D., Reissner, C., and Garner, C.C. (2016). Role of Bassoon and Piccolo in Assembly and Molecular Organization of the Active Zone. Front Synaptic Neuro 7.

Haas, B.R., and Sontheimer, H. (2010). Inhibition of the Sodium-Potassium-Chloride Cotransporter Isoform-1 reduces glioma invasion. Cancer Res 70, 5597–5606.

Habela, C.W., Ernest, N.J., Swindall, A.F., and Sontheimer, H. (2009). Chloride accumulation drives volume dynamics underlying cell proliferation and migration. J Neurophysiol 101, 750–757.

Haber, M., Zhou, L., and Murai, K.K. (2006). Cooperative astrocyte and dendritic spine dynamics at hippocampal excitatory synapses. J Neurosci 26, 8881–8891.

Hama, H., Kurokawa, H., Kawano, H., Ando, R., Shimogori, T., Noda, H., Fukami, K., Sakaue-Sawano, A., and Miyawaki, A. (2011). Scale: a chemical approach for fluorescence imaging and reconstruction of transparent mouse brain. Nat Neurosci 14, 1481–1488.

Harris, K.M., Jensen, F.E., and Tsao, B. (1992). Three-dimensional structure of dendritic spines and synapses in rat hippocampus (CA1) at postnatal day 15 and adult ages: implications for the maturation of synaptic physiology and long-term potentiation. J Neurosci 12, 2685–2705.

Harvey, C.D., and Svoboda, K. (2007). Locally dynamic synaptic learning rules in pyramidal neuron dendrites. Nature 450, 1195–U1193.

Heller, J.P., Michaluk, P., Sugao, K., and Rusakov, D.A. (2017). Probing nano-organization of astroglia with multi-color super-resolution microscopy. J Neurosci Res 95, 2159–2171.

Heller, J.P., and Rusakov, D.A. (2015). Morphological plasticity of astroglia: Understanding synaptic microenvironment. Glia 63, 2133–2151.

Henneberger, C., Papouin, T., Oliet, S.H., and Rusakov, D.A. (2010). Long-term potentiation depends on release of D-serine from astrocytes. Nature 463, 232–236.

Hires, S.A., Zhu, Y., and Tsien, R.Y. (2008). Optical measurement of synaptic glutamate spillover and reuptake by linker optimized glutamate-sensitive fluorescent reporters. Proc Natl Acad Sci U S A 105, 4411–4416.

Hirrlinger, J., Hulsmann, S., and Kirchhoff, F. (2004). Astroglial processes show spontaneous motility at active synaptic terminals in situ. Eur J Neurosci 20, 2235–2239.

Hoffmann, E.K., Lambert, I.H., and Pedersen, S.F. (2009). Physiology of Cell Volume Regulation in Vertebrates. Physiol Rev 89, 193–277.

Igarashi, H., Huber, V.J., Tsujita, M., and Nakada, T. (2011). Pretreatment with a novel aquaporin 4 inhibitor, TGN-020, significantly reduces ischemic cerebral edema. Neurol Sci 32, 113–116.

Isaacson, J.S. (1999). Glutamate spillover mediates excitatory transmission in the rat olfactory bulb. Neuron 23, 377–384.

Jensen, T.P., Zheng, K., Cole, N., Marvin, J.S., Looger, L.L., and Rusakov, D.A. (2019). Multiplex imaging relates quantal glutamate release to presynaptic Ca2+ homeostasis at multiple synapses in situ. Nature Communications 10, 1414.

Jones, T.A., and Greenough, W.T. (1996). Ultrastructural evidence for increased contact between astrocytes and synapses in rats reared in a complex environment. Neurobiology of Learning and Memory 65, 48–56.

Jourdain, P., Bergersen, L.H., Bhaukaurally, K., Bezzi, P., Santello, M., Domercq, M., Matute, C., Tonello, F., Gundersen, V., and Volterra, A. (2007). Glutamate exocytosis from astrocytes controls synaptic strength. Nature Neurosci 10, 331–339.

Kaila, K., Price, T.J., Payne, J.A., Puskarjov, M., and Voipio, J. (2014). Cation-chloride cotransporters in neuronal development, plasticity and disease. Nature Rev Neurosci 15, 637–654.

Kochlamazashvili, G., Henneberger, C., Bukalo, O., Dvoretskova, E., Senkov, O., Lievens, P.M., Westenbroek, R., Engel, A.K., Catterall, W.A., Rusakov, D.A., et al. (2010). The extracellular matrix molecule hyaluronic acid regulates hippocampal synaptic plasticity by modulating postsynaptic L-type Ca^2+^ channels. Neuron 67, 116–128.

Korogod, N., Petersen, C.C.H., and Knott, G.W. (2015). Ultrastructural analysis of adult mouse neocortex comparing aldehyde perfusion with cryo fixation. Elife 4.

Kullmann, D.M., and Asztely, F. (1998). Extrasynaptic glutamate spillover in the hippocampus: evidence and implications. Trends Neurosci 21, 8–14.

Kullmann, D.M., Erdemli, G., and Asztely, F. (1996). LTP of AMPA and NMDA receptor-mediated signals: evidence for presynaptic expression and extrasynaptic glutamate spill-over. Neuron 17, 461–474.

Lehre, K.P., and Danbolt, N.C. (1998). The number of glutamate transporter subtype molecules at glutamatergic synapses: Chemical and stereological quantification in young adult rat brain. J Neurosci 18, 8751–8757.

Lehre, K.P., and Rusakov, D.A. (2002). Asymmetry of glia near central synapses favors presynaptically directed glutamate escape. Biophys J 83, 125–134.

Liu, A., Zhou, Z.K., Dang, R., Zhu, Y.H., Qi, J.X., He, G.Q., Leung, C., Pak, D., Jia, Z.P., and Xie, W. (2016). Neuroligin 1 regulates spines and synaptic plasticity via LIMK1/cofilin-mediated actin reorganization. J Cell Biol 212, 449–463.

Llano, O., Smirnov, S., Soni, S., Golubtsov, A., Guillemin, I., Hotulainen, P., Medina, I., Nothwang, H.G., Rivera, C., and Ludwig, A. (2015). KCC2 regulates actin dynamics in dendritic spines via interaction with beta-PIX. J Cell Biol 209, 671–686.

Lozovaya, N.A., Kopanitsa, M.V., Boychuk, Y.A., and Krishtal, O.A. (1999). Enhancement of glutamate release uncovers spillover-mediated transmission by N-methyl-D-aspartate receptors in the rat hippocampus. Neurosci 91, 1321–1330.

Luscher, C., Malenka, R.C., and Nicoll, R.A. (1998). Monitoring glutamate release during LTP with glial transporter currents. Neuron 21, 435–441.

Lushnikova, I., Skibo, G., Muller, D., and Nikonenko, I. (2009). Synaptic potentiation induces increased glial coverage of excitatory synapses in CA1 hippocampus. Hippocampus 19, 753–762.

Manabe, T., and Nicoll, R.A. (1994). Long-Term Potentiation - Evidence Against an Increase in Transmitter Release Probability in the Ca1 Region of the Hippocampus. Science 265, 1888–1892.

Marvin, J.S., Borghuis, B.G., Tian, L., Cichon, J., Harnett, M.T., Akerboom, J., Gordus, A., Renninger, S.L., Chen, T.W., Bargmann, C.I., et al. (2013). An optimized fluorescent probe for visualizing glutamate neurotransmission. Nature Methods 10, 162–170.

Matsuzaki, M., Ellis-Davies, G.C., Nemoto, T., Miyashita, Y., Iino, M., and Kasai, H. (2001). Dendritic spine geometry is critical for AMPA receptor expression in hippocampal CA1 pyramidal neurons. Nat Neurosci 4, 1086–1092.

Matsuzaki, M., Honkura, N., Ellis-Davies, G.C., and Kasai, H. (2004). Structural basis of long-term potentiation in single dendritic spines. Nature 429, 761–766.

Medvedev, N., Popov, V., Henneberger, C., Kraev, I., Rusakov, D.A., and Stewart, M.G. (2014). Glia selectively approach synapses on thin dendritic spines. Philos Trans R Soc Lond B Biol Sci 369.

Medvedev, N.I., Popov, V.I., Rodriguez Arellano, J.J., Dallerac, G., Davies, H.A., Gabbott, P.L., Laroche, S., Kraev, I.V., Doyere, V., and Stewart, M.G. (2010). The N-methyl-D-aspartate receptor antagonist CPP alters synapse and spine structure and impairs long-term potentiation and long-term depression induced morphological plasticity in dentate gyrus of the awake rat. Neurosci 165, 1170–1181.

Megevand, P., Troncoso, E., Quairiaux, C., Muller, D., Michel, C.M., and Kiss, J.Z. (2009). Long-term plasticity in mouse sensorimotor circuits after rhythmic whisker stimulation. J Neurosci 29, 5326–5335.

Metcalf, D.J., Edwards, R., Kumarswami, N., and Knight, A.E. (2013). Test samples for optimizing STORM super-resolution microscopy. J Vis Exp.

Migliati, E., Meurice, N., DuBois, P., Fang, J.S., Somasekharan, S., Beckett, E., Flynn, G., and Yool, A.J. (2009). Inhibition of aquaporin-1 and aquaporin-4 water permeability by a derivative of the loop diuretic bumetanide acting at an internal pore-occluding binding site. Mol Pharmacol 76, 105–112.

Min, M.Y., Rusakov, D.A., and Kullmann, D.M. (1998). Activation of AMPA, kainate, and metabotropic receptors at hippocampal mossy fiber synapses: role of glutamate diffusion. Neuron 21, 561–570.

Min, R., and Nevian, T. (2012). Astrocyte signaling controls spike timing-dependent depression at neocortical synapses. Nat Neurosci 15, 746–753.

Mishra, A., Reynolds, J.P., Chen, Y., Gourine, A.V., Rusakov, D.A., and Attwell, D. (2016). Astrocytes mediate neurovascular signaling to capillary pericytes but not to arterioles. Nature Neurosci 19, 1619–1627.

Murai, K.K., Nguyen, L.N., Irie, F., Yamaguchi, Y., and Pasquale, E.B. (2003). Control of hippocampal dendritic spine morphology through ephrin-A3/EphA4 signaling. Nat Neurosci 6, 153–160.

Nagelhus, E.A., and Ottersen, O.P. (2013). Physiological roles of aquaporin-4 in brain. Physiol Rev 93, 1543–1562.

Nagerl, U.V., Eberhorn, N., Cambridge, S.B., and Bonhoeffer, T. (2004). Bidirectional activity-dependent morphological plasticity in hippocampal neurons. Neuron 44, 759–767.

Navarrete, M., and Araque, A. (2010). Endocannabinoids potentiate synaptic transmission through stimulation of astrocytes. Neuron 68, 113–126.

Nishida, H., and Okabe, S. (2007). Direct astrocytic contacts regulate local maturation of dendritic spines. J Neurosci 27, 331–340.

Okubo, Y., Sekiya, H., Namiki, S., Sakamoto, H., Iinuma, S., Yamasaki, M., Watanabe, M., Hirose, K., and Iino, M. (2010). Imaging extrasynaptic glutamate dynamics in the brain. Proc Natl Acad Sci U S A 107, 6526–6531.

Okumoto, S., Looger, L.L., Micheva, K.D., Reimer, R.J., Smith, S.J., and Frommer, W.B. (2005). Detection of glutamate release from neurons by genetically encoded surface-displayed FRET nanosensors. Proc Natl Acad Sci USA 102, 8740–8745.

Oliet, S.H.R., Piet, R., and Poulain, D.A. (2001). Control of glutamate clearance and synaptic efficacy by glial coverage of neurons. Science 292, 923–926.

Ostroff, L.E., Manzur, M.K., Cain, C.K., and Ledoux, J.E. (2014). Synapses lacking astrocyte appear in the amygdala during consolidation of Pavlovian threat conditioning. The Journal of comparative neurology 522, 2152–2163.

Otmakhov, N., Khibnik, L., Otmakhova, N., Carpenter, S., Riahi, S., Asrican, B., and Lisman, J. (2004). Forskolin-induced LTP in the CA1 hippocampal region is NMDA receptor dependent. J Neurophysiol 91, 1955–1962.

Panatier, A., Arizono, M., and Nagerl, U.V. (2014). Dissecting tripartite synapses with STED microscopy. Philos Trans R Soc Lond B Biol Sci 369, 20130597.

Panatier, A., Theodosis, D.T., Mothet, J.P., Touquet, B., Pollegioni, L., Poulain, D.A., and Oliet, S.H. (2006). Glia-derived D-serine controls NMDA receptor activity and synaptic memory. Cell 125, 775–784.

Panatier, A., Vallee, J., Haber, M., Murai, K.K., Lacaille, J.C., and Robitaille, R. (2011). Astrocytes are endogenous regulators of basal transmission at central synapses. Cell 146, 785–798.

Pascual, O., Casper, K.B., Kubera, C., Zhang, J., Revilla-Sanchez, R., Sul, J.Y., Takano, H., Moss, S.J., McCarthy, K., and Haydon, P.G. (2005). Astrocytic purinergic signaling coordinates synaptic networks. Science 310, 113–116.

Pereira, A.C., Lambert, H.K., Grossman, Y.S., Dumitriu, D., Waldman, R., Jannetty, S.K., Calakos, K., Janssen, W.G., McEwen, B.S., and Morrison, J.H. (2014). Glutamatergic regulation prevents hippocampal-dependent age-related cognitive decline through dendritic spine clustering. Proc Natl Acad Sci U S A 111, 18733–18738.

Perez-Alvarez, A., Navarrete, M., Covelo, A., Martin, E.D., and Araque, A. (2014). Structural and functional plasticity of astrocyte processes and dendritic spine interactions. J Neurosci 34, 12738–12744.

Peters, A., and Kaiserman-Abramof, I.R. (1970). The small pyramidal neuron of the rat cerebral cortex. The perikaryon, dendrites and spines. Am J Anat 127, 321–355.

Popov, V., Medvedev, N.I., Davies, H.A., and Stewart, M.G. (2005). Mitochondria form a filamentous reticular network in hippocampal dendrites but are present as discrete bodies in axons: a three-dimensional ultrastructural study. J Comp Neurol 492, 50–65.

Popov, V.I., Davies, H.A., Rogachevsky, V.V., Patrushev, I.V., Errington, M.L., Gabbot, P.L.A., Bliss, T.V.P., and Stewart, M.G. (2004). Remodelling of synaptic morphology but unchanged synaptic density during late phase long-term potentiation (LTP): A serial section electron micrograph study in the dentate gyrus in the anaesthetised rat. Neurosci 128, 251–262.

Porter, J.T., and McCarthy, K.D. (1997). Astrocytic neurotransmitter receptors in situ and in vivo. Prog Neurobiol 51, 439–455.

Reeves, A.M., Shigetomi, E., and Khakh, B.S. (2011). Bulk loading of calcium indicator dyes to study astrocyte physiology: key limitations and improvements using morphological maps. J Neurosci 31, 9353–9358.

Reynolds, J.P., Zheng, K., and Rusakov, D.A. (2018). Multiplexed calcium imaging of single-synapse activity and astroglial responses in the intact brain. Neurosci Lett.

Rusakov, D.A. (2001). The role of perisynaptic glial sheaths in glutamate spillover and extracellular Ca^2+^ depletion. Biophys J 81, 1947–1959.

Rusakov, D.A. (2015). Disentangling calcium-driven astrocyte physiology. Nature Rev Neurosci 16, 226–233.

Rusakov, D.A., Harrison, E., and Stewart, M.G. (1998). Synapses in hippocampus occupy only 1-2% of cell membranes and are spaced less than half-micron apart: a quantitative ultrastructural analysis with discussion of physiological implications. Neuropharmacol 37, 513–521.

Rusakov, D.A., and Kullmann, D.M. (1998). Extrasynaptic glutamate diffusion in the hippocampus: ultrastructural constraints, uptake, and receptor activation. J Neurosci 18, 3158–3170.

Rusakov, D.A., Kullmann, D.M., and Stewart, M.G. (1999). Hippocampal synapses: do they talk to their neighbours? Trends Neurosci 22, 382–388.

Russell, J.M. (2000). Sodium-potassium-chloride cotransport. Physiol Rev 80, 211–276.

Santello, M., Bezzi, P., and Volterra, A. (2011). TNFalpha controls glutamatergic gliotransmission in the hippocampal dentate gyrus. Neuron 69, 988–1001.

Savtchenko, L.P., Bard, L., Jensen, T.P., Reynolds, J.P., Kraev, I., Medvedev, N., Stewart, M.G., Henneberger, C., and Rusakov, D.A. (2018). Disentangling astroglial physiology with a realistic cell model in silico. Nature Communications 9, 3554.

Savtchenko, L.P., and Rusakov, D.A. (2005). Extracellular diffusivity determines contribution of high-versus low-affinity receptors to neural signaling. Neuroimage 25, 101–111.

Savtchenko, L.P., Sylantyev, S., and Rusakov, D.A. (2013). Central synapses release a resource-efficient amount of glutamate. Nature Neurosci 16, 10–U163.

Scanziani, M., Salin, P.A., Vogt, K.E., Malenka, R.C., and Nicoll, R.A. (1997). Use-dependent increases in glutamate concentration activate presynaptic metabotropic glutamate receptors. Nature 385, 630–634.

Schiapparelli, P., Guerrero-Cazares, H., Magana-Maldonado, R., Hamilla, S.M., Ganaha, S., Goulin Lippi Fernandes, E., Huang, C.H., Aranda-Espinoza, H., Devreotes, P., and Quinones-Hinojosa, A. (2017). NKCC1 Regulates Migration Ability of Glioblastoma Cells by Modulation of Actin Dynamics and Interacting with Cofilin. EBioMedicine, 10.1016/j.ebiom.2017.1006.1020.

Scimemi, A., Fine, A., Kullmann, D.M., and Rusakov, D.A. (2004). NR2B-containing receptors mediate cross talk among hippocampal synapses. J Neurosci 24, 4767–4777.

Shen, H.W., Scofield, M.D., Boger, H., Hensley, M., and Kalivas, P.W. (2014). Synaptic glutamate spillover due to impaired glutamate uptake mediates heroin relapse. J Neurosci 34, 5649–5657.

Shepherd, G.M.G., and Harris, K.M. (1998). Three-dimensional structure and composition of CA3 -> CA1 axons in rat hippocampal slices: Implications for presynaptic connectivity and compartmentalization. J Neurosci 18, 8300–8310.

Shigetomi, E., Jackson-Weaver, O., Huckstepp, R.T., O’Dell, T.J., and Khakh, B.S. (2013). TRPA1 channels are regulators of astrocyte basal calcium levels and long-term potentiation via constitutive D-serine release. J Neurosci 33, 10143–10153.

Smith, A.C.W., Scofield, M.D., Heinsbroek, J.A., Gipson, C.D., Neuhofer, D., Roberts-Wolfe, D.J., Spencer, S., Garcia-Keller, C., Stankeviciute, N.M., Smith, R.J., et al. (2017). Accumbens nNOS Interneurons Regulate Cocaine Relapse. J Neurosci 37, 742–756.

Sykova, E., and Nicholson, C. (2008). Diffusion in brain extracellular space. Physiol Rev 88, 1277–1340.

Szapiro, G., and Barbour, B. (2007). Multiple climbing fibers signal to molecular layer interneurons exclusively via glutamate spillover. Nature Neurosci 10, 735–742.

Tanaka, M., Shih, P.Y., Gomi, H., Yoshida, T., Nakai, J., Ando, R., Furuichi, T., Mikoshiba, K., Semyanov, A., and Itohara, S. (2013). Astrocytic Ca2+ signals are required for the functional integrity of tripartite synapses. Molecular Brain 6.

Thrane, A.S., Rappold, P.M., Fujita, T., Torres, A., Bekar, L.K., Takano, T., Peng, W.G., Wang, F.S., Thrane, V.R., Enger, R., et al. (2011). Critical role of aquaporin-4 (AQP4) in astrocytic Ca2+ signaling events elicited by cerebral edema. Proc Natl Acad Sci USA 108, 846–851.

Tonnesen, J., Inavalli, V.V.G.K., and Nagerl, U.V. (2018). Super-resolution imaging of the extracellular space in living brain tissue. Cell 172, 1108–1121.

Tonnesen, J., Katona, G., Rozsa, B., and Nagerl, U.V. (2014). Spine neck plasticity regulates compartmentalization of synapses. Nat Neurosci 17, 678–685.

Tonnesen, J., Nadrigny, F., Willig, K.I., Wedlich-Soldner, R., and Nagerl, U.V. (2011). Two-color STED microscopy of living synapses using a single laser-beam pair. Biophys J 101, 2545–2552.

Tradtrantip, L., Jin, B.J., Yao, X., Anderson, M.O., and Verkman, A.S. (2017). Aquaporin-Targeted Therapeutics: State-of-the-Field. Adv Exp Med Biol 969, 239–250.

Tsvetkov, E., Shin, R.M., and Bolshakov, V.Y. (2004). Glutamate uptake determines pathway specificity of long-term potentiation in the neural circuitry of fear conditioning. Neuron 41, 139–151.

Ventura, R., and Harris, K.M. (1999). Three-dimensional relationships between hippocampal synapses and astrocytes. J Neurosci 19, 6897–6906.

Vogt, K.E., and Nicoll, R.A. (1999). Glutamate and gamma-aminobutyric acid mediate a heterosynaptic depression at mossy fiber synapses in the hippocampus. Proc Natl Acad Sci USA 96, 1118–1122.

Volterra, A., Liaudet, N., and Savtchouk, I. (2014). Astrocyte Ca^2+^ signalling: an unexpected complexity. Nat Rev Neurosci 15, 327–335.

Watkins, S., and Sontheimer, H. (2011). Hydrodynamic cellular volume changes enable glioma cell invasion. J Neurosci 31, 17250–17259.

Wenzel, J., Lammert, G., Meyer, U., and Krug, M. (1991). The influence of long-term potentiation on the spatial relationship between astrocyte processes and potentiated synapses in the dentate gyrus neuropil of rat-brain. Brain Res 560, 122–131.

Whitfield, J.H., Zhang, W.H., Herde, M.K., Clifton, B.E., Radziejewski, J., Janovjak, H., Henneberger, C., and Jackson, C.J. (2015). Construction of a robust and sensitive arginine biosensor through ancestral protein reconstruction. Protein Science 24, 1412–1422.

Yasuda, R., Sabatini, B.L., and Svoboda, K. (2003). Plasticity of calcium channels in dendritic spines. Nat Neurosci 6, 948–955.

Zheng, K., Bard, L., Reynolds, J.P., King, C., Jensen, T.P., Gourine, A.V., and Rusakov, D.A. (2015). Time-resolved imaging reveals heterogeneous landscapes of nanomolar Ca^2+^ in neurons and astroglia. Neuron 88, 277–288.

